# Awakening a latent carbon fixation cycle in *Escherichia coli*

**DOI:** 10.1101/2020.05.18.102244

**Authors:** Ari Satanowski, Beau Dronsella, Elad Noor, Bastian Vögeli, Hai He, Philipp Wichmann, Tobias J. Erb, Steffen N. Lindner, Arren Bar-Even

## Abstract

Carbon fixation is one of the most important biochemical processes. Most natural carbon fixation pathways are thought to have emerged from enzymes that originally performed other metabolic tasks. Can we recreate the emergence of a carbon fixation pathway in a heterotrophic host by recruiting only endogenous enzymes? In this study, we address this question by systematically analyzing possible carbon fixation pathways composed only of *Escherichia coli* native enzymes. We identify the GED (<u>G</u>nd-<u>E</u>ntner-<u>D</u>oudoroff) cycle as the simplest pathway that can operate with high thermodynamic driving force. This autocatalytic route is based on reductive carboxylation of ribulose 5-phosphate (Ru5P) by 6-phosphogluconate dehydrogenase (Gnd), followed by reactions of the Entner-Doudoroff pathway, gluconeogenesis, and the pentose phosphate pathway. We demonstrate the *in vivo* feasibility of this new-to-nature pathway by constructing *E. coli* gene deletion strains whose growth on pentose sugars depends on the GED shunt, a linear variant of the GED cycle which does not require the regeneration of Ru5P. Several metabolic adaptations, most importantly the increased production of NADPH, assist in establishing sufficiently high flux to sustain this growth. Our study exemplifies a trajectory for the emergence of carbon fixation in a heterotrophic organism and demonstrates a synthetic pathway of biotechnological interest.

## Introduction

The ability to assimilate inorganic carbon into biomass sets a clear distinction between autotrophic primary producers and the heterotrophs depending on them for the supply of organic carbon. Most primary production occurs via the ribulose bisphosphate (RuBP) cycle – better known as the Calvin-Benson cycle – used in bacteria, algae, and plants ^1^. Five other carbon fixation pathways are known to operate in various bacterial and archaeal lineages ^2, 3^ and also contribute to primary production ^1^. Recent studies have made considerable progress in establishing carbon fixation pathways in heterotrophic organisms, with the long-term goal of achieving synthetic autotrophy, which could pave the way towards sustainable bioproduction schemes rooted in CO2 and renewable energy ^4, 5^. Most notably, overexpression of phosphoribulokinase and Rubisco, followed by long-term evolution, enabled the industrial hosts *Escherichia coli* ^6, 7^ and *Pichia pastoris*^8^ to synthesize all biomass from CO_2_ via the RuBP cycle. Also, overexpression of enzymes of the 3-hydroxypropionate bicycle established the activity of different modules of this carbon fixation pathway in *E. coli* ^9^. While such studies help us to gain a deeper understanding of the physiological changes required to adapt a heterotrophic organism to autotrophic growth, they do not, however, shed light on the origin of the carbon fixation pathways themselves.

Besides the reductive acetyl-CoA pathway and the reductive TCA cycle – both of which are believed to have originated early in the evolution of metabolism ^10^ – the other carbon fixation routes are thought to have evolved by recruiting enzymes from other metabolic pathways. For example, Rubisco – the carboxylating enzyme of the RuBP cycle – probably evolved from a non-CO_2_-fixing ancestral enzyme, thus emerging in a non-autotrophic context ^11^. Similarly, acetyl-CoA carboxylase likely originated as a key component of fatty acid biosynthesis before being recruited into carbon fixation pathways in several prokaryotic lineages ^2, 3^. The limited number of natural carbon fixation pathways might indicate that the recruitment of endogenous enzymes to support carbon fixation is a rather exceptional event. To understand this process better we aimed to recreate it in a heterotrophic bacterium.

Here, we use a computational approach to comprehensively search for all thermodynamically feasible carbon fixation pathways that rely solely on endogenous *E. coli* enzymes. We identify a promising candidate route – the GED cycle – that is expected to enable carbon fixation with minimal reactions and with a high thermodynamic driving force. This synthetic route combines reductive carboxylation of ribulose 5-phosphate (Ru5P) with the Entner–Doudoroff (ED) pathway, gluconeogenesis, and the pentose phosphate pathway. The GED cycle mirrors the structure of the RuBP cycle and is expected to outperform it in terms of ATP-efficiency. We demonstrate that overexpression of key pathway enzymes together with small modifications of the endogenous metabolic network enable growth via the “GED shunt” – a linear route that requires the key reactions of the GED cycle, including the carboxylation step, for the biosynthesis of (almost) all biomass building blocks. Our findings indicate the feasibility of recruiting endogenous enzymes to establish a novel carbon fixation pathway and pave the way for future establishment of synthetic autotrophy based on new-to-nature pathways.

## Results

### Systematic search for latent carbon fixation pathways in *E. coli*

To identify possible carbon fixation pathways that can be established using only native *E. coli* enzymes ^12^, we used the genome-scale metabolic model of this bacterium ^13^. We assumed that all reactions are reversible and then used an algorithm to systematically uncover all possible combinations of enzymes, the net reaction of which use CO_2_ and cofactors (e.g., ATP, NAD(P)H) as sole substrates to produce pyruvate, a reference product commonly used to compare carbon fixation pathways ^2-4^ (Methods and Supplementary Text). The pathways were then analyzed thermodynamically: for each pathway, we calculated the Max-min Driving Force (MDF), representing the smallest driving force among all pathway reactions after optimizing metabolite concentrations within a physiological range (Methods and Supplementary Text) ^14^. We assumed an elevated CO_2_ concentration of 20% (200 mbar), which is easily attainable in microbial cultivation within an industrial context and further characterizes the natural habitats of *E. coli*, e.g., the mammalian gut ^15, 16^. The MDF criterion enabled us to discard thermodynamically infeasible routes (having MDF<0) and to compare the feasible pathways according to their energetic driving force, which directly affects their kinetics ^14^.

Using this approach, we identified multiple carbon fixation pathways based on endogenous *E. coli* enzymes (see Supplementary Text for a full list). We ranked the pathways according to two key criteria that can be calculated for each of them in a straightforward manner: their MDF and the number of enzymes they require (preferring fewer enzymes, see Methods and Supplementary Text). Pathways ranked high in terms of these criteria are expected to be simpler to establish and to operate more robustly under fluctuating physiological conditions. The pathway that was ranked highest (Fig. 1A) had the lowest number of enzymes while still supporting a high MDF (> 3 kJ/mol, such that reverse enzyme flux is minimal ^14^). This cycle is based on the reductive carboxylation of ribulose 5-phosphate (Ru5P) by 6-phosphogluconate (6PG) dehydrogenase (Gnd). The carboxylation product, 6PG, is then metabolized by the enzymes of the Entner–Doudoroff (ED) pathway – 6PG dehydratase (Edd) and 2-keto-3-deoxygluconate 6-phosphate aldolase (Eda) – to produce glyceraldehyde 3-phosphate (GAP) and pyruvate (Fig. 1A). Pyruvate is subsequently converted to GAP via native gluconeogenesis, and GAP is metabolized via the pentose phosphate pathway to regenerate Ru5P, thus completing the cycle. We termed this pathway the GED (Gnd-Entner-Doudoroff) cycle, according to its key enzymes that serve to connect CO_2_ fixation to central metabolism.

**Figure 1.**
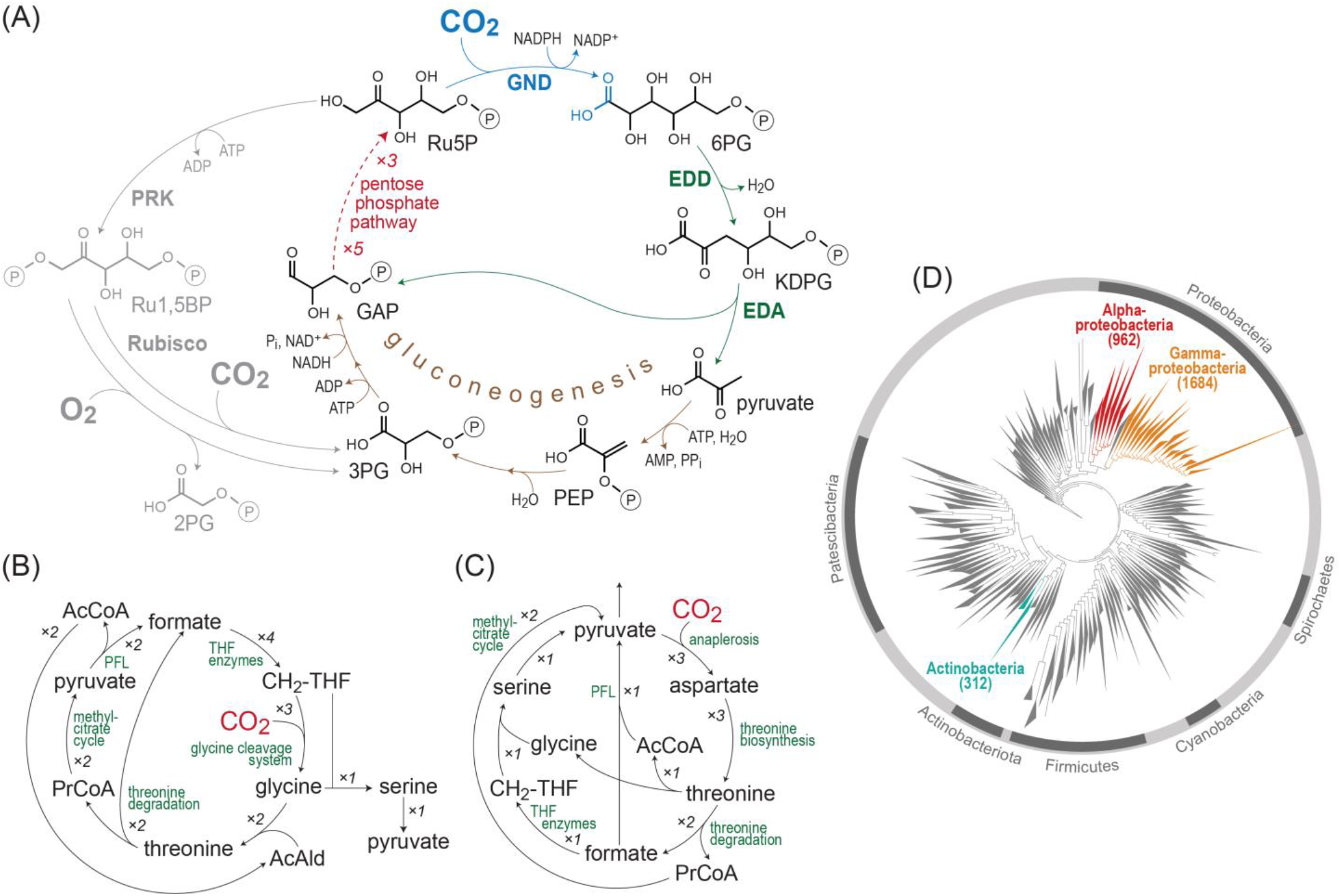
*In silico* identification of latent carbon fixation pathways. (A) The GED cycle, the simplest carbon fixation pathway that can be generated from *E. coli* endogenous enzymes. The pathway is based on the reductive carboxylation of ribulose 5-phosphate (Ru5P) to 6-phosphogluconate (6PG) by Gnd, followed by the activity of the Entner–Doudoroff pathway (enzymes Edd and Eda) to produce pyruvate and glyceraldehyde 3-phosphate (GAP). Pyruvate is converted to GAP via gluconeogenesis and the intermediates phosphoenolpyruvate (PEP) and 3-phosphoglycerate (3PG). To close the cycle, GAP is recycled to Ru5P via the pentose phosphate pathway. The GED cycle mirrors the ribulose bisphosphate cycle (i.e., Calvin-Benson cycle), which is shown in grey. (B,C) Two other general archetypes of carbon fixation pathways that were computationally uncovered. Both are based on integrated cycles, which together reduce CO_2_ to formate and then assimilate formate into pyruvate. AcCoA corresponds to acetyl-CoA and PrCoA corresponds to propionyl-CoA. CH_2_-THF corresponds to methylene-THF. (D) A phylogenetic tree of bacteria, showing the three phyla that harbor all key enzymes of the GED cycle (shown in color). Numbers in parentheses correspond to the number of species in which the key pathway enzymes were found.

Our computational analysis identified multiple variants of the GED cycle (Supplementary Text). However, as these are unnecessarily more complex than the simple GED cycle design, we decided not to consider them further. We also identified numerous pathways that do not require Gnd. These Gnd-independent pathways share a common carbon fixation strategy in which a sub-cycle converts CO_2_ to formate, where the release of formate is catalyzed by an oxygen-sensitive oxoacid formate-lyase ^17^ (Supplementary Text). Formate is then assimilated via one of several variants of the reductive glycine pathway ^18, 19^ (see Fig. 1B for an example), the activity of which was recently demonstrated in *E. coli* ^20, 21^. Alternatively, formate is assimilated via variants of the serine cycle, or, more precisely, the previously suggested serine-threonine cycle ^22^ (see Fig. 1C for an example). While these formate-dependent pathways are interesting, their high oxygen sensitivity, as well as general complexity, make them less attractive. Therefore, for further investigation, we decided to focus on the GED cycle.

### Properties of the GED cycle and its enzymes

The GED cycle mirrors the structure of the canonical RuBP cycle, where phosphoribulokinase and Rubisco are replaced with Gnd, Edd, Eda, and gluconeogenic enzymes (Fig. 1). Similarly to the RuBP cycle, the GED cycle is autocatalytic and any one of its intermediates can be used as a product to be diverted towards biosynthesis of cellular building blocks ^23^. Production of pyruvate, a key biosynthetic building block, is more ATP-efficient via the GED cycle than via the RuBP cycle: while the former pathway requires 6 ATP molecules to generate pyruvate, the latter pathway needs 7 ATP molecules (not accounting for further losses due to Rubisco’s oxygenation reaction and the resulting photorespiration). Indeed, using Flux Balance Analysis, we found that, compared to the RuBP cycle, the GED cycle is consistently expected to support higher yields of biomass and multiple products that are derived from pyruvate or acetyl-CoA, including ethanol, lactate, isobutanol, 2,3-butanediol, acetone, butyrate, n-butanol, citrate, itaconate, 2-ketoglutarate, and levulinic acid (Supplementary Fig. S1 and Methods).

All enzymes of the GED cycle are known to carry flux in the direction required for pathway activity, with the exception of Gnd. The Gibbs energy of the Gnd reaction indicates that it should be fully reversible under elevated CO_2_: Δ_r_G’^m^ ≈ −1.5 kJ/mol in the oxidative decarboxylation direction (pH 7.5, ionic strength of 0.25 M, [CO_2_] = 200 mbar, and 1 mM concentration of the other reactants ^24^). Indeed, similar oxidative decarboxylation enzymes are known to support reductive carboxylation, for example, the malic enzyme ^25-27^ and isocitrate dehydrogenase ^28, 29^. While sporadic studies have reported that some Gnd variants support the reductive carboxylation of Ru5P *in vitro* ^30-34^, a comprehensive kinetic characterization of this activity in bacterial Gnd variants is lacking. More importantly, it remains unclear whether this reaction could operate under physiological conditions, where the concentrations of substrates and products are constrained; that is, substrate concentrations are not necessarily saturating and product concentrations are non-negligible.

First, we measured the kinetics of *E. coli* Gnd. We found Gnd to have a rather high k_cat_ in the reductive carboxylation direction, approaching 6 s^-1^ (5.9 ± 0.2 s^-1^ with Ru5P as substrate, Table 1 and Supplementary Fig. S2), about twice as high as the k_cat_ of most plant Rubisco variants ^35^. The affinity of Gnd towards CO_2_ is high enough to enable saturation under elevated CO_2_ concentrations: KM = 0.9 ± 0.1 mM (Table 1 and Supplementary Fig. S2) which is equivalent to ~3% CO_2_ in the headspace (at ambient pressure). Notably, these kinetic parameters are substantially better than those previously reported for a eukaryotic Gnd variant (k_cat_ ~ 1 s^-1^ and K_M_(CO_2_) ≥ 15 mM ^30^, ^31^).

**Table 1.** Kinetic parameters of *E. coli* Gnd in the reductive and oxidative directions. Fitted Michaelis-Menten curves are shown in Supplementary Fig. S2. See Methods for a detailed description of the kinetic characterization.

| | $k_{cat}$ (s<sup>-1</sup>) | $K_M$ (mM) |
| --- | --- | --- |
| *Reductive carboxylation* |  |  |
| ribulose 5-phosphate | 5.9 ± 0.2 | 2.8 ± 0.2 |
| NADPH | 5.2 ± 0.2 | 1.0 ± 0.1 |
| CO<sub>2</sub> | 4.7 ± 0.2 | 0.9 ± 0.1 |
| *Oxidative decarboxylation* |  |  |
| 6-phosphogluconate | 49 ± 1 | 0.035 ± 0.002 |
| NADP<sup>+</sup> | 46 ± 1 | 0.012 ± 0.003 |

To check how prevalent the potential of carbon fixation via the GED cycle is, we performed a phylogenetic analysis to identify bacteria that harbor its key enzymes (Methods). We found that the enzymes of the GED cycle are ubiquitous in alpha-proteobacteria (962 species), gamma-proteobacteria (1684), and actinobacteria (312) (Fig. 1D). The species of these phyla might therefore be prime candidates to search for the carbon fixation activity of the GED cycle. However, in other bacterial lineages, the combined occurrence of the GED cycle enzymes is quite rare, with only 10 other species harboring all key enzymes.

### Selection for the activity of the GED shunt within a Δ*rpe* context

Engineering *E. coli* for autotrophic growth via the GED cycle would be a challenging task requiring considerable metabolic adaptation of the host; for example, establishing a delicate balance between the metabolic fluxes within the cycle and those diverging out of the cycle (as found in previous efforts to establish the RuBP cycle in *E. coli* ^6, 7^). Hence, to check the feasibility of the cycle, we focused on establishing growth via the “GED shunt”, representing a segment of the full cycle which consists of reductive carboxylation by Gnd and the subsequent ED pathway (blue reactions in Fig. 2A; for a similar approach see ref. ^36, 37^). As we show below, growth via this linear shunt requires the activity of most enzymes of the GED cycle, but relies on a pentose substrate rather than regeneration of Ru5P.

**Figure 2.**
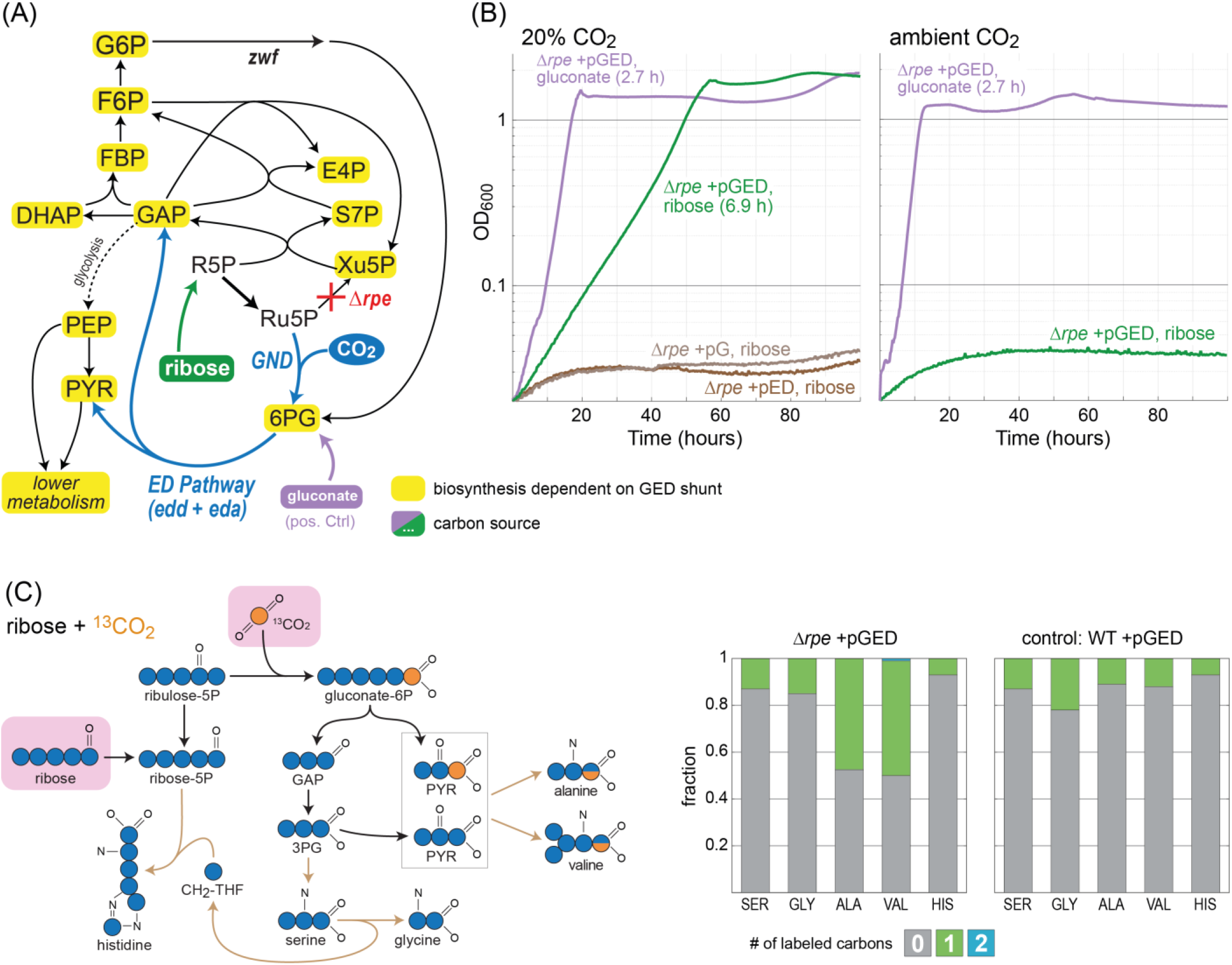
Activity of the GED shunt in a Δ*rpe* strain. (A) Design of the Δ*rpe* selection scheme. Ribose can be assimilated only via the activity of the GED shunt, where the biosynthesis of almost all biomass building blocks is dependent on the pathway (marked in yellow). Growth on gluconate (violet) is not dependent on reductive carboxylation via Gnd and thus serves as a positive control. Reaction directionalities are shown as predicted by Flux Balance Analysis. (B) Growth of a Δ*rpe* strain on ribose (20mM) as sole carbon source is dependent on elevated CO_2_ concentration (20%, i.e., 200 mbar) and overexpression of *gnd, edd*, and *eda* (pGED). Overexpression of only *gnd* (pG) or only *edd* and *eda* (pED) failed to establish growth (less than two doublings). Cultivation at ambient CO_2_ also failed to achieve growth. Values in parentheses indicate doubling times. Curves represent the average of technical duplicates, which differ from each other by less than 5%. Growth experiments were repeated independently three times to ensure reproducibility. (C) Cultivation on ^13^CO_2_ confirms operation of the GED shunt. On the left, a prediction of the labeling pattern of key amino acids is shown. The observed labeling fits the prediction and differs from the WT control cultivated under the same conditions. Labeling of amino acids in the WT strain stems from the natural occurrence of ^13^C as well as from reactions that exchange cellular carbon with CO_2_, e.g., the glycine cleavage system and anaplerotic/cataplerotic cycling. Values represent averages of two independent cultures that differ from each other by less than 10%. Abbreviations: 3PG, 3-phosphoglycerate; ALA, Alanine; GAP, glyceraldehyde-3-phosphate; GLY, Glycine; HIS, Histidine; PYR, pyruvate; SER, Serine; VAL, Valine.

First, we generated an *E. coli* strain deleted in the gene encoding for ribulose 5-phosphate 3-epimerase (Δ*rpe*). This strain cannot grow on ribose as a sole carbon source, as ribose 5-phosphate (R5P) cannot be converted to xylulose 5-phosphate, thus blocking the pentose phosphate pathway (Fig 2A) ^38, 39^. Activity of Gnd, Edd, and Eda should restore growth by enabling conversion of R5P to GAP and pyruvate, from which all cellular building blocks can be derived (Fig. 2A). This would enable direct selection for the activity of the GED shunt.

We found that overexpression of Gnd, Edd, and Eda from a plasmid (pGED) enabled the growth of the Δ *rpe* strain on ribose only under elevated CO_2_ concentration (green lines in Fig. 2B). The observed growth rate via the GED shunt is almost half of that obtained with gluconate, which requires no carboxylation by Gnd and thus serves as a positive control (doubling times 6.9 h and 2.8 h, respectively). While Gnd, Edd, and Eda are all present in the genome of *E. coli*, their native expression level is too low to enable sufficient activity of the GED shunt: overexpression of Gnd alone (pG) or of Edd and Eda alone (pED) did not support growth (less than two doublings, brown lines in Fig. 2B).

To confirm that growth indeed proceeds via the GED shunt, we performed a ^13^C-labeling experiment. We cultivated the Δ*rpe* +pGED strain with unlabeled ribose and ^13^CO_2_, and measured the labeling pattern of five proteinogenic amino acids – serine, glycine, alanine, valine, and histidine. The results confirm the activity of the GED shunt (Fig. 2C): (i) since ^13^CO_2_ is incorporated as the carboxylic carbon of 6PG, GAP is completely unlabeled and hence serine and glycine that are derived from it are unlabeled; (ii) pyruvate is generated both from GAP and directly from Eda activity (Fig. 2C), such that about half of the pyruvate molecules are unlabeled and half are labeled once at their carboxylic carbon – as a result, half of the alanine and valine molecules are labeled (during valine biosynthesis two pyruvate molecules are condensed and one carboxylic carbon is lost as CO_2_).

These results confirm that, upon overexpression of Gnd, Edd, and Eda, the GED shunt is sufficiently active to provide the cell with almost all cellular building blocks as well as energy (by complete oxidation of pyruvate via the TCA cycle). As mentioned above, the growth of this strain requires the simultaneous activity of most enzymes on the GED cycle, including those of glycolysis and the pentose phosphate pathway; for example, net production of erythrose 4-phosphate (E4P) from ribose requires the combined activity of Gnd, the ED pathway, and enzymes of the pentose phosphate pathway.

### A Δ*tktAB* context requires additional metabolic adaptations to enable growth via the GED shunt

To check whether the operation of the GED shunt is robust, we decided to use another metabolic background to select for its activity. We deleted the genes encoding for both isozymes of transketolase (Δ*tktAB*). This strain, in which the non-oxidative pentose phosphate pathway is effectively abolished, cannot grow when provided with a pentose as sole carbon source ^36, 40^. Furthermore, as E4P cannot be synthesized in this strain, small amounts of essential cellular components, the biosynthesis of which is E4P-dependent, need to be added to the media ^36, 40^: phenylalanine, tyrosine, tryptophan, shikimate, pyridoxine, 4-aminobenzoate, 4-hydroxybenzoate, and 2,3-dihydroxybenzoate (referred to as E4P-supplements, see Methods).

As with the Δ*rpe* strain, we expected overexpression of Gnd, Edd, and Eda to enable growth on a pentose substrate such as xylose (supplemented with E4P-supplements) (Fig. 3A). However, we failed to obtain growth even at an elevated CO_2_ concentration (less than two doublings, green lines in Fig. 3B). This is in line with previous findings that seemingly small differences in the design of metabolic growth selection schemes (e.g. the choice of deleted enzymes) can lead to substantially dissimilar metabolic behaviors ^36, 41^.

**Figure 3.**
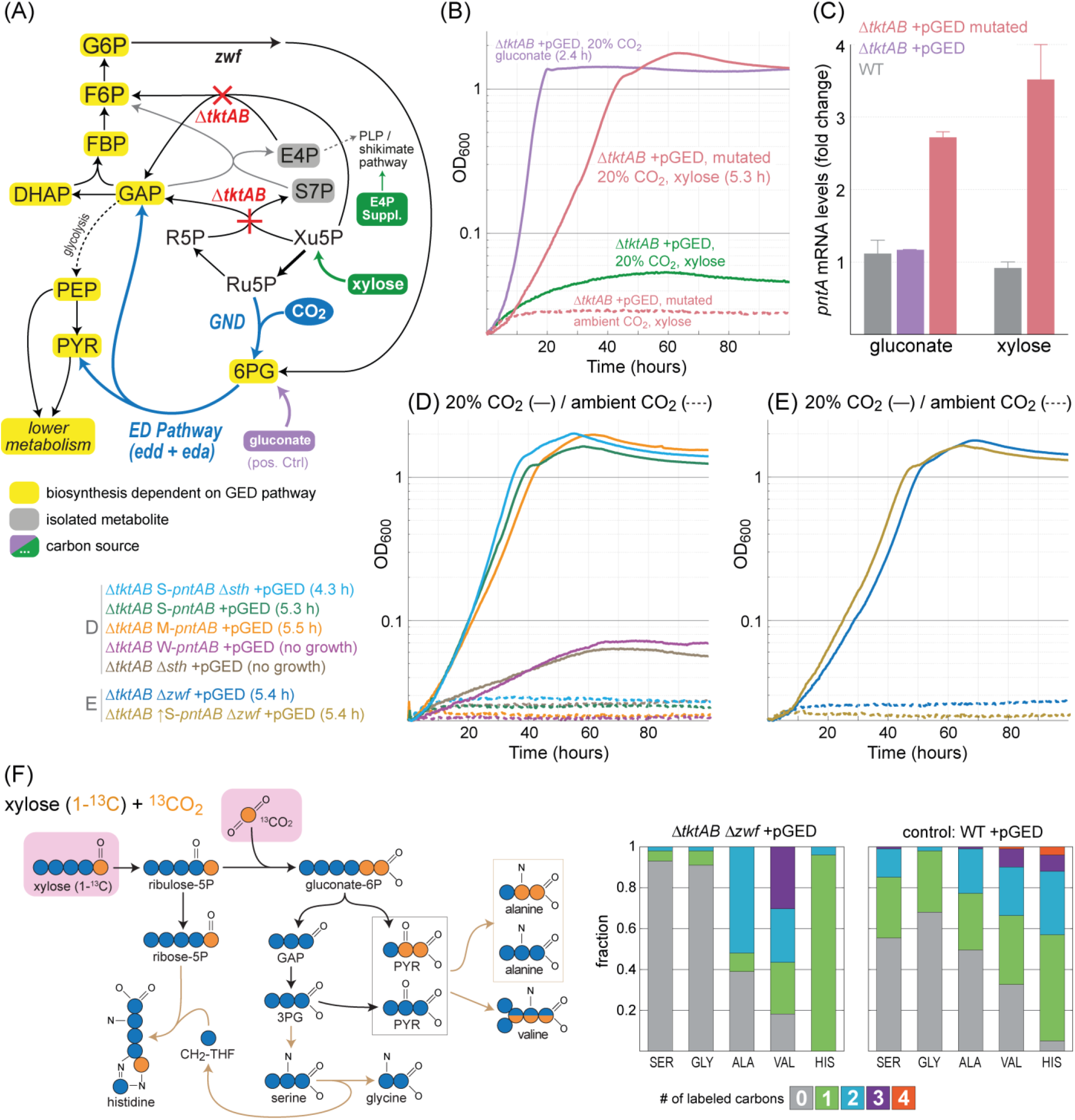
Activity of the GED shunt in a Δ*tktAB* strain. (A) Design of the Δ*tktAB* selection scheme. Xylose can be assimilated only via the GED shunt. E4P supplements are provided as the Δ*tkt*AB strain cannot synthesize erythrose 4-phosphate. Growth on gluconate is not dependent on reductive carboxylation by Gnd and thus serves as a positive control. Reaction directionalities are shown as predicted by Flux Balance Analysis. (B) Growth on xylose upon overexpression of *gnd, edd*, and *eda* (pGED) was achieved only after mutation and was still dependent on elevated CO_2_ concentration. Values in parentheses indicate doubling times. Curves represent the average of technical duplicates, which differ from each other by <5%. Growth experiments were repeated independently three times to ensure reproducibility. (C) Expression analysis by quantitative RT-PCR revealed that the transcript level of *pntA* increased ~3 fold in the mutated strain. Values correspond to the average of two independent experiments; error bars indicating the standard errors. Gluconate and xylose indicate carbon sources. (D) Genomic overexpression of *pntAB* using medium (M) or strong (S) promoter, but not weak (W) promoter, supported growth of a Δ*tkt*AB pGED strain on xylose (legend to the left). (E) Deletion of glucose 6-phosphate dehydrogenase (Δ*zwf*) supported growth of a Δ*tkt*AB pGED strain on xylose (legend to the left). (F) ^13^C-labeling experiments confirm the operation of the GED shunt. Cells were cultivated with xylose (1-^13^C) and ^13^CO_2_. Observed labeling fits the expected pattern and differs from that of a WT strain cultured under the same conditions. Results from additional labeling experiments are shown in Supplementary Fig. S3. Abbreviations: 3PG, 3-phospho-glycerate; ALA, Alanine; GAP, glyceraldehyde-3-phosphate; GLY, Glycine; HIS, Histidine; PYR, pyruvate; SER, Serine; VAL, Valine.

Following the failure to obtain GED shunt-dependent growth within the Δ*tktAB* context, we sought to harness adaptive evolution to provide us with information on further cellular adaptations required for the activity of the synthetic route. Towards this aim, we inoculated the Δ*tktAB* +pGED strain into multiple test-tubes with xylose and E4P-supplements and incubated them for an extended period of time at 37°C and 20% CO_2_. After ~2 weeks, the culture in several of these test-tubes showed apparent growth. When the cells from the growing cultures were transferred to fresh selective medium, we observed immediate growth, indicating that genetic adaptation had occurred. Yet, only one of these cultures showed robust growth on the selective medium, whereas that of the others was less reproducible and highly sensitive to the exact conditions, preculture, and inoculation. An isolated single clone from this robust culture grew, under elevated CO_2_ concentration, with a doubling time of 5.3 h (red lines in Fig. 3B).

We sequenced the genome of the mutated strain and identified a single mutation (compared to the parental strain): the mobile element IS5 ^42^ was inserted 104 base pairs upstream of the *pntAB* operon (Supplementary Data), which encodes for the membrane-bound transhydrogenase that plays a key role in supplying the cell with NADPH ^43, 44^. Insertion of the IS5 mobile element is well-known to occur in adaptive evolution experiments, increasing the expression levels of the downstream genes ^45, 46^. Indeed, we found that the transcription of *pntA* increased ~3-fold in the mutated strain compared to the non-mutated parent and WT strains (Fig. 3C). The contribution of this mutation to the activity of the GED shunt can be easily explained, as it increases the generation of NADPH required for the reductive carboxylation of Ru5P by Gnd (Fig. 1).

To confirm that increased *pntAB* expression indeed enables GED shunt-dependent growth, we replaced the native promoter of *pntAB* (within the unmutated strain) with three previously characterized constitutive promoters: weak (W-*pntAB*), medium (M-*pntAB*), and strong (S-*pntAB*), with relative strengths of 1:10:20, respectively ^47^. We found that while the weak promoter failed to support growth (less than two doublings, purple line in Fig. 3D), the medium and strong promoters supported growth with a similar doubling time to the mutated strain, ~5.4 h, at elevated CO_2_ concentrations (orange and green lines in Fig. 3D). This indicates that sustaining sufficiently high expression of *pntAB* suffices to enable the activity of the GED shunt within a Δ*tktAB* metabolic context.

We wondered whether other metabolic manipulations that target NADPH homeostasis could also enable the growth of the Δ*tktAB* +pGED strain. We tested the deletion of the gene encoding for the soluble transhydrogenase (Δ*sth*), as this enzyme is known to provide a strong sink for NADPH ^43, 44^. Yet, the Δ*tktAB Δsth* +pGED strain did not grow on xylose even at an elevated CO_2_ concentration (less than two doublings, brown lines in Fig. 3D). Furthermore, deletion of *sth* in the Δ*tktAB S-pntAB* +pGED strain improved its growth only marginally (light blue line in Fig. 3D). Taken together, it seems that the soluble transhydrogenase has only a minor effect on NADPH availability within this metabolic context.

We hypothesized that alongside NADPH availability, competing sources of 6PG could play a key role in determining the feasibility of the GED shunt. Specifically, 6PG is natively produced by the oxidative pentose phosphate pathway, which thus provides a metabolic push against the reductive activity of Gnd. Moreover, the activity of glucose 6-phosphate 1-dehydrogenase (encoded by *zwf*), the first enzyme of the oxidative pentose phosphate pathway, has been reported to increase under conditions of high NADPH demand (i.e. upon depletion of cellular NADPH levels) ^48, 49^. Hence, we wondered whether deletion of *zwf* could remove a barrier for reductive carboxylation by Gnd and thus assist the activity of the GED shunt. We found that this is indeed the case, where the Δ*tktAB Δzwf* +pGED strain was able to grow under elevated CO_2_ concentrations with a doubling time of 5.4 h (blue line in Fig. 3E). Deleting *zwf* in the Δ*tktAB S-pntAB* +pGED strain did not improve growth (dark yellow line in Fig. 3E), suggesting that the effects of NADPH and 6PG availability are not additive or that a different bottleneck is limiting growth.

To provide an unequivocal confirmation for the activity of the GED shunt within the Δ*tktAB* metabolic context we performed several ^13^C-labeling experiments (Fig. 3F and Supplementary Fig. S3). When the Δ*tktAB Δzwf* +pGED strain was fed with both ^13^CO_2_ and 1-^13^C-xylose, we expected the GED shunt to produce unlabeled GAP and twice labeled pyruvate (Fig. 3F). Hence, serine and glycine, which are derived directly from GAP, should be unlabeled while about half of pyruvate (derived from GAP metabolism) should be unlabeled while the other half (generated directly by Eda activity) should be twice labeled. This should lead to half of the alanine being unlabeled and half twice labeled while the labeling of valine should roughly follow a 1:1:1:1 pattern (unlabeled: one labeled: twice labeled: trice labeled). The observed labeling confirms these expected patterns (Fig. 3F). Histidine, the carbons of which originate from R5P and the β-carbon of serine, is labeled once as expected (Fig. 3F). The labeling patterns we observe upon feeding with unlabeled xylose and ^13^CO_2_ (Supplementary Fig. S3A) as well as upon feeding with 5-^13^C-xylose and unlabeled CO_2_ (Supplementary Fig. S3B) further confirm that growth of the Δ*tktAB Δzwf* +pGED strain indeed takes place exclusively via the GED shunt.

### Growth via the GED shunt in a strain that could support cyclic flux

While the Δ*rpe* and Δ*tktAB* strains were useful selection platforms to test the activity of the GED shunt, they are, in a sense, ‘metabolic dead-ends’. This is because the activities of both ribulose-phosphate 3-epimerase (Rpe) and transketolase (Tkt) are essential for the operation of the full GED cycle, that is, for the regeneration of Ru5P from GAP. To address this problem, we aimed to construct a strain which keeps all necessary enzymes of the GED cycle intact, while still allowing to select for the activity of the GED shunt, i.e., preventing utilization of a pentose substrate as sole carbon source via the canonical pentose phosphate pathway. Such a strain would enable a smooth transition from GED shunt-dependent growth on a pentose substrate towards autotrophic growth via the GED cycle.

We therefore constructed a strain deleted in all enzymes that can metabolize fructose 6-phosphate, directly or indirectly, into a downstream glycolytic intermediate (Δ*pfkAB* Δ*fsaAB* Δ*fruK*) or channel it into the oxidative pentose phosphate pathway (Δ*zwf*). The latter gene deletion should also support the activity of the GED shunt, as was shown above within the Δ*tktAB* context. The strain containing all of these deletions, which we term ΔPZF, establishes a uni-directional block within the pentose phosphate pathway. That is, growth on a pentose substrate is not possible due to the accumulation of fructose 6-phosphate that prevents further conversion of pentose phosphates into GAP (Fig. 4A) ^7^. In contrast, flux in the opposite direction, as required for the GED cycle, can still occur, since fructose 1,6-bisphosphate can be dephosphorylated to fructose 6-phosphate which is then used to regenerate Ru5P.

**Figure 4.**
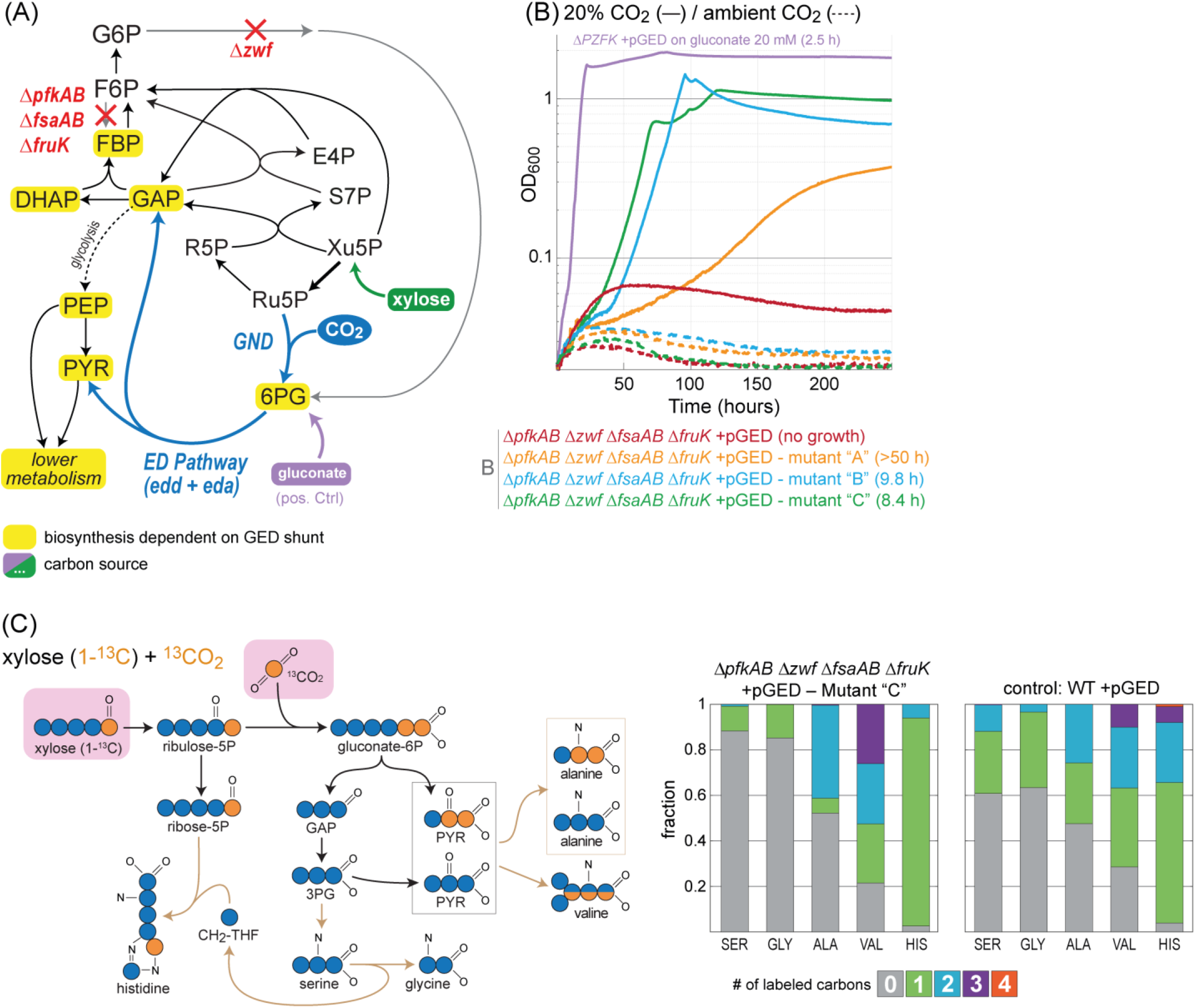
Activity of the GED shunt in a Δ*PZF* strain which enables a smooth transition into a GED cycle. (A) Design of the ΔPZF (ΔpfkAB Δzwf ΔfsaAB Δ*fruK*) selection scheme. Xylose can be assimilated only via the activity of the GED shunt, where the biosynthesis of most biomass building blocks is dependent on the pathway (marked in yellow). Growth on gluconate (violet) is not dependent on reductive carboxylation via Gnd and thus serves as a positive control. Reaction directionalities are shown as predicted by Flux Balance Analysis. (B) Growth on xylose upon overexpression of *gnd, edd*, and *eda* (pGED) was achieved only after adaptive evolution and was dependent on elevated CO_2_ concentration. Values in parentheses indicate doubling times. Curves represent the average of technical quadruplicates, which differ from each other by <5%. Growth experiments were repeated independently three times to ensure reproducibility. (C) ^13^C-labeling experiments confirm the operation of the GED shunt (mutant ‘C’). Cells were cultivated with xylose (1-^13^C) and ^13^CO_2_. Observed labeling fits the expected pattern and differs from that of a WT strain cultured under the same conditions. Results from additional labeling experiments are shown in Supplementary Fig. S4. Abbreviations: 3PG, 3-phosphoglycerate; ALA, Alanine; GAP, glyceraldehyde-3-phosphate; GLY, Glycine; HIS, Histidine; PYR, pyruvate; SER, Serine; VAL, Valine.

To establish growth of the ΔPZF strain on xylose via the GED shunt, we overexpressed Gnd, Edd, and Eda. However, transforming the ΔPZF strain with pGED failed to support growth on xylose even at elevated CO_2_ (less than two doublings, red line in Fig. 4B). Hence, we again harnessed natural selection and performed short-term evolution by incubating the strain in multiple test-tubes for an extended period of time in xylose minimal medium at 37°C and 20% CO_2_. Within 6-8 days, three parallel cultures started growing. Isolated clones from two of these three mutant cultures displayed a fairly high growth rate (doubling time of 8-10 hours, green and blue lines in Fig. 4B), while clones from the third culture showed considerably slower growth (doubling time > 50 h, orange line in Fig. 4B). We conducted ^13^C-labeling experiments using the fastest-growing strain (ΔPZF +pGED mutant “C”) and found that it displayed a labeling pattern almost identical to that of the Δ*tktAB* Δ*zwf* +pGED strain described above, thus confirming growth via the GED shunt (Fig. 4C and Supplementary Fig. S4). We sequenced the genomes of the mutant strains, compared them to the parental strain, and discovered several mutations (Supplementary Table 1). All isolated colonies from the two fast-growing cultures shared an identical mutation at the start of an L-leucyl-tRNA (*leuX*) and, in most colonies, *avtA*, encoding for valine-pyruvate aminotransferase, had mutated. While the exact contribution of these mutations to the growth phenotype remains elusive and could be further investigated, the isolated strains provide a promising starting point for evolution of the full GED cycle.

## Discussion

Our computational analysis identified multiple carbon fixation pathways that are based solely on endogenous *E. coli* enzymes. A key factor for the success of this analysis was to ignore the rather arbitrary dichotomic classification of reactions as reversible or irreversible as suggested by metabolic models. Instead, we first identified potential pathways based on pure stoichiometric analysis, and then calculated the thermodynamic feasibility and driving force of each of the candidate routes. This enabled us to uncover potential carbon fixation pathways that were not identified before ^12^. Indeed, the GED cycle itself was previously ignored as Gnd was considered to be an irreversible decarboxylating enzyme. As we have shown here, however, Gnd can catalyze the carboxylation reaction quite efficiently, with a k_cat_ almost double that of a typical plant Rubisco ^35^. This finding is similar to a recent study that found that citrate synthase – which is usually thought to be irreversible – can catalyze citrate cleavage, thus enabling carbon fixation via a unique variant of the reductive TCA cycle ^50, 51^. These examples indicate that we should revise our dogmatic interpretation of enzyme reversibility and instead adopt a more quantitative approach to understand reaction directionality.

Previous studies have suggested multiple synthetic carbon fixation pathways that could surpass the natural routes in terms of resource use efficiency, thermodynamics, and/or kinetics ^52, 53^. The most advanced of these pathways is the CETCH cycle ^53^ that combines segments of the 3-hydroxypropionate/4-hydroxybutyrate cycle ^54^ and the ethylmalonyl-CoA pathway ^55^. The CETCH cycle was assembled *in vitro* using enzymes from nine organisms and optimized in several rounds of enzyme engineering ^53, 56^. However, the *in vivo* implementation of this synthetic pathway, as well as of other previously suggested routes, is highly challenging due to its complexity and requirement for considerable rewiring of central metabolic fluxes. The GED cycle provides a favorable alternative to these routes, as the fluxes it requires mostly correspond to native gluconeogenesis and the pentose phosphate pathway. Hence, the establishment of carbon fixation via the GED cycle might be less demanding and more likely to succeed.

We demonstrated the feasibility of carbon fixation via the GED cycle by establishing growth via the GED shunt – a linear pathway variant that requires the key pathway reactions to provide (almost) all biomass building blocks and cellular energy without regenerating the substrate Ru5P (Fig. 2A and 3A). In line with previous studies, we found that changing the metabolic context can have a dramatic effect on the activity of a metabolic module under selection: the GED shunt was able to directly support growth of a Δ*rpe* strain but not of a Δ*tktAB* strain or a ΔPZF strain, even though in all of them the pentose phosphate pathway is disrupted. Despite this, a short-term adaptation was able to restore the growth of the latter two strains. Within the Δ*tktAB* strain, we demonstrated that either an increase in the supply of the substrate (e.g., NADPH, via *pntAB* overexpression) or a decrease in the availability of the product (e.g., 6PG, via *zwf* deletion) is sufficient to enable Gnd-dependent carboxylation and growth via the GED shunt.

The GED shunt might have biotechnological advantages on its own. By using Flux Balance Analysis, we found that rerouting the utilization of sugar substrates via the GED shunt is expected to increase the yield of various commercially interesting products, such as acetate, pyruvate, acetone, citrate, itaconate, and levulinic acid (Fig. 5, Supplementary Fig. S5, and Methods). This is attributed to the fact that the biosynthesis of these compounds from sugar feedstocks results in the production of excess reducing power, which the GED shunt can utilize to fix CO_2_ and generate more product. We note that this assimilation of CO_2_ also serves to compensate for the carbon released during the oxidation of pyruvate to acetyl-CoA, thus addressing a common challenge for the production of value-added chemicals derived from acetyl-CoA ^57, 58^. Further supply of reducing power by adding auxiliary substrates such as hydrogen or formate ^59^ can make the GED shunt advantageous over glycolysis for even more reduced products, such as ethanol, lactate, n-butanol, and isobutanol (Fig. 5, Supplementary Fig. S5, and Methods). While the RuBP shunt – a linear version of the RuBP cycle, which channels Ru5P via Rubisco – can also increase fermentative yield of some products ^60-62^, the GED shunt always outperforms it due to a lower ATP requirement (Fig. 5 and Supplementary Fig. S5).

**Figure 5.**
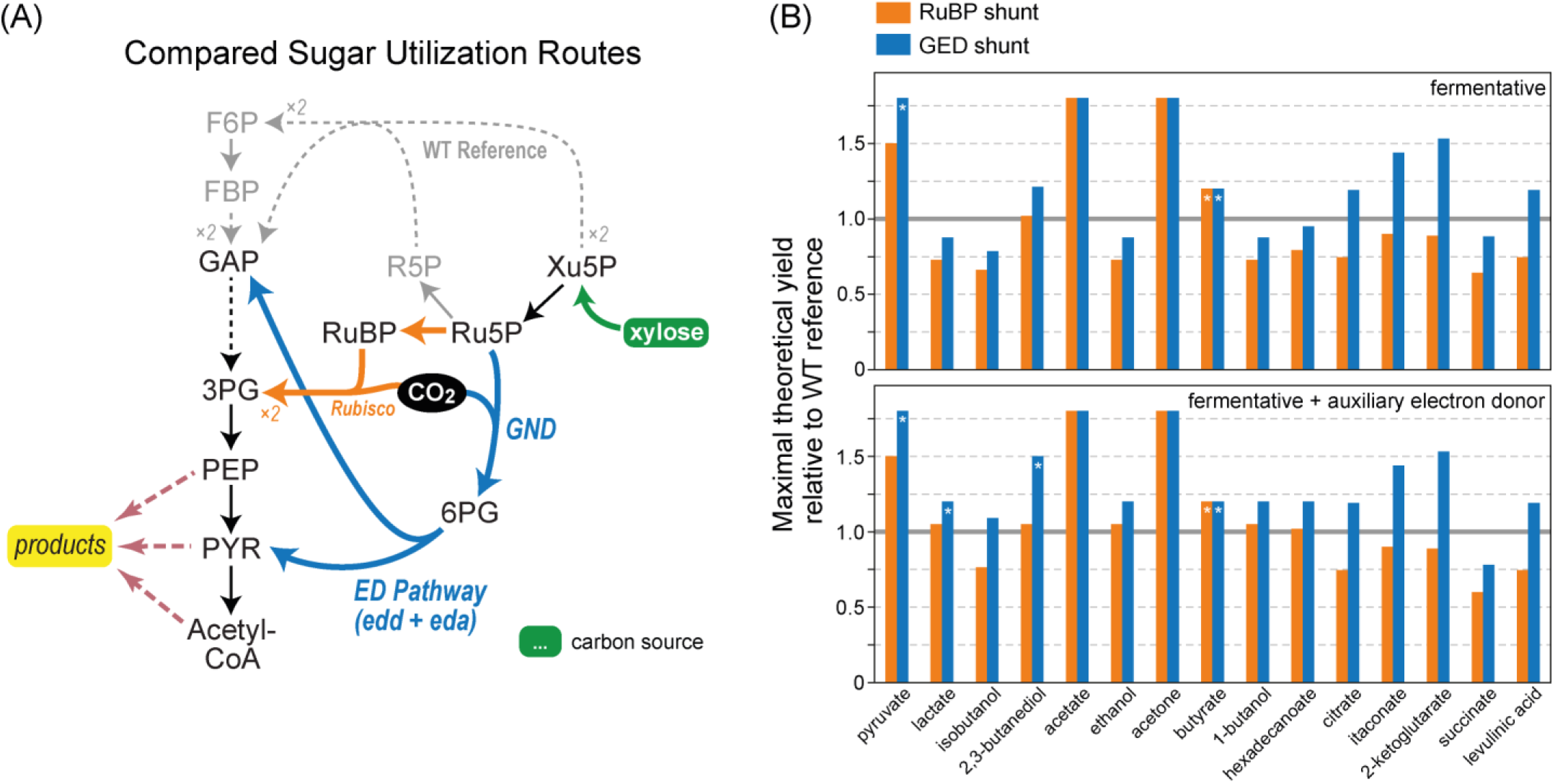
Rerouting sugar fermentation via the GED shunt can increase product yields. In this example, we chose xylose as the sugar substrate, due to its growing relevance as a renewable feedstock derived from lignocellulosic wastes. An additional analysis for glucose as the fermentation feedstock is shown in Supplementary Fig. S5. (A) An overview comparing xylose utilization routes: Canonical utilization (gray) via the pentose phosphate pathway and glycolysis; the GED shunt (blue); and a linear pathway based on carboxylation by Rubisco (“RuBP shunt”, orange). (B) Maximal theoretical yields of 15 fermentation products are shown relative to the WT reference, as calculated by Flux Balance Analysis. Presented values are normalized to the product yield from canonical sugar utilization (pentose phosphate pathway and glycolysis). Most products required the secretion of other organic compounds (e.g. acetate, formate) to achieve balancing of reducing equivalents or support ATP biosynthesis. Asterisks denote products that could be produced without byproducts.

A previous study has established the RuBP cycle in *E. coli*, demonstrating that this heterotrophic bacterium can be modified to grow autotrophically with CO_2_ as a sole carbon source ^6, 7^. However, to our knowledge, the current study is the first one in which the capacity for net carbon fixation was explored *in vivo* using only endogenous enzymes of a heterotrophic host, thus shedding light on the emergence of novel carbon fixation pathways. Importantly, the establishment of the RuBP cycle in *E. coli* required long-term adaptive evolution of the microbe under selective conditions, which modulated the partitioning of metabolic fluxes between carbon fixation and biosynthetic pathways ^23, 63^. We expect that autotrophic growth via the GED cycle can be achieved in a similar manner. The ΔPZF strain serves as an ideal starting point for such a future evolution experiment, as its growth is dependent on the activity of the GED shunt while it still harbors all necessary enzymes to run the GED cycle. The gradual evolution of autotrophic growth via the GED cycle would be achieved via the additional expression of formate dehydrogenase as an energy-supplying module and long-term cultivation with limiting amounts of xylose and saturating amounts of CO_2_ and formate ^6, 7^.

As the GED cycle is composed of ubiquitous enzymes that are widespread throughout major bacterial phyla, it is tempting to speculate that this route naturally operates in yet unexplored microorganisms. Especially promising are α-proteobacteria, γ-proteobacteria, and actinobacteria, which contain many species that harbor all key enzymes of the pathway (Fig. 1D). Such bacteria could evolve autotrophic growth by recruiting the enzymes of the GED cycle if exposed to the appropriate selective conditions – for example, lack of organic carbon sources and availability of energy sources such as inorganic electron donors (e.g., hydrogen).

The GED cycle could be used to replace the RuBP cycle in plants, algae, and bacteria ^64^, requiring relatively modest changes to the endogenous metabolic structure of carbon fixation. Replacing the RuBP cycle in chemolithotrophic bacteria of biotechnological significance, e.g., *Cupriavidus necator* ^65^, would be relatively straightforward as cultivating these microorganisms on elevated CO_2_ is a common practice, thus avoiding the rate limitation associated with the low affinity of Gnd to CO_2_. Such an engineered microorganism could support higher product yields when cultivated under autotrophic conditions as pyruvate and acetyl-CoA, from which most value-added chemicals are derived, are produced more efficiently via the GED cycle than via the RuBP cycle (Supplementary Fig. S1). Furthermore, as the carboxylating activity of Gnd would be enhanced by the carbon concentrating mechanisms of algae ^66^ and cyanobacteria ^67^, engineering these organisms to use the GED cycle could be advantageous. However, to facilitate the establishment of the GED cycle in higher plants, the affinity of Gnd towards CO_2_ would have to be improved, e.g., via the rational engineering of CO_2_ binding sites ^68^, as successfully demonstrated recently in a proof-of-principle study ^69^. Alternatively, replacing *E. coli* Gnd with a variant that has a considerably higher k_cat_ (~100 s^-1^) could compensate for the low affinity towards CO_2_ (i.e., achieving k_cat_/KM at least as high as that of plant Rubisco). As some variants of similar reductive carboxylation enzymes – isocitrate dehydrogenase and the malic enzyme – incorporate CO_2_ with k_cat_ surpassing 100 s^-1 70,^ identifying a Gnd variant supporting such a high carboxylation rate might be feasible. Engineering such an optimized GED cycle into crop plants could boost agricultural productivity, thus addressing one of our key societal challenges ^64^.

## Methods

### Identifying carbon fixation cycles in *E. coli* using a constraint-based model

In order to find all possible carbon fixation cycles using *E. coli* endogenous enzymes, we used an approach similar to the one we have previously developed ^52^. In this previous study, all reactions found in the KEGG database ^71^ (denoted the universal stoichiometric matrix) were used to design potential CO_2_ fixating pathways using a Mixed-Integer Linear Programming (MILP) approach. Here, we focused only on enzymes present in the most recent genome-scale metabolic reconstruction of *E. coli: i*ML1515 ^13^. We further added thermodynamic constraints in order to exclude infeasible pathways and to rank the feasible ones based on their Max-min Driving Force (MDF) ^14^. The source code and all input and output files required for repeating this analysis can be found on GitLab (https://gitlab.com/elad.noor/path-designer/) and are available under an open-source license (MIT).

We used the COBRApy package to identify carbon fixation pathways ^72^. First, we removed all exchange and transport reactions and kept only the strictly cytoplasmic ones. Then, we added reactions that allow the free flow of electrons and energy (by regenerating ATP, NADH, and NADPH) as well as inorganic compounds (protons, water, oxygen, and ammonia). Finally, we defined an optimization problem, where the set of reactions in each pathway should overall convert 3 moles of CO_2_ to one mole of pyruvate. The objective function of this optimization is a combination of the MDF ^14^ and the minimum number of reactions in each pathway. The standard approach for multi-objective optimization is maximization of a linear combination of the two functions. Here, we maximized the MDF (in units of *RT*) minus the number of reactions. We note that changing the relative weight between these two objectives did not change our main result, namely that the GED pathway is Pareto-optimal. A more detailed list of changes to the model, and the formal description of the optimization problem can be found in the Supplementary Text.

We used the framework described above to iterate the space of possible solutions, i.e., all pathways that have a positive MDF. This requirement ensures that the pathway is thermodynamically feasible given our constraints, i.e. all of its reactions can operate simultaneously without violating the second law of thermodynamics. Covering this space exhaustively would require many days of computer time and is not useful since most of these solutions are unnecessarily complicated. The Pareto-optimal solutions are those that are not dominated by any other pathway in both objectives simultaneously (i.e. compared to any other pathways, they either have either a higher MDF or a lower number of reactions). We found only two such pathways: the GED cycle and a pathway which uses the reverse glycine cleavage system as its carboxylating mechanism (similar to the one shown in Fig. 1B). If we remove all the oxygen-sensitive reactions (specifically, PFL and OBTFL) then the GED cycle is the only Pareto-optimal pathway, i.e. it has the minimum number of steps and the highest MDF possible. For more details on the method, and for the list of all identified pathways, see the Supplementary Text.

### Phylogenetic analysis

In order to assess how many bacterial genomes contain the genes necessary for GED, we used AnnoTree - a web tool for visualization of genome annotations in prokaryotes (http://annotree.uwaterloo.ca/) ^73^. AnnoTree generates a phylogenetic tree and highlights genomes that include all of the KEGG orthologies ^71^ selected in the query. Since most annotations are homology based, and therefore not always precise, we restricted our search to the enzymes that are most indicative of the GED pathway and cannot be easily replaced by a metabolic bypass (e.g., transaldolase could be replaced by sedoheptulose 1,7-bisphosphate aldolase and phosphatase):

- K00033 – 6-phosphogluconate dehydrogenase (EC:1.1.1.44 1.1.1.343): the enzymatic keystone of the GED cycle.
- K01690 – 6-phosphogluconate dehydratase [EC:4.2.1.12]: indicative of the Entner–Doudoroff pathway.
- K00615 – transketolase [EC:2.2.1.1]: indicative of the pentose phosphate pathway.
- K00927 – phosphoglycerate kinase [EC:2.7.2.3]: indicative of gluconeogenesis (as glycolysis can operate irreversibly via a non-phosphorylating glyceraldehyde 3-phosphate dehydrogenase).

We chose the family level as the tree resolution, since it provides a good balance between number of leaves and branching diversity. For the sake of readability of Fig. 1D, we applied the *auto collapse clades* option in iTOL with the setting BRL < 1.

### Yield estimation via Flux Balance Analysis

Flux Balance Analysis was conducted in Python with COBRApy ^72^. We used the most updated *E. coli* genome-scale metabolic network *i*ML1515 ^13^ with several curations and changes: (i) transhydrogenase (THD2pp) translocates one proton instead of two ^74^; (ii) homoserine dehydrogenase (HSDy) produces homoserine from aspartate-semialdehyde irreversibly instead of reversibly ^75^; (iii) GLYCK (glycerate-3P producing glycerate kinase) and POR5 (pyruvate synthase) were removed from the model as their existence in *E. coli* is highly disputable; (iv) the exchange reactions of Fe2+, H2S, methionine and cysteine were removed to prevent respiration with inorganic Fe or S as electron acceptors in simulated anaerobic conditions; (v) the threonine cleavage routes were blocked by deleting threonine aldolase (THRA), threonine dehydrogenase (THRD), and glycine C-acetyltransferase (GLYAT); (vi) two unrealistic pathways for the conversion of pentoses to acetyl-CoA were removed by deleting reactions DRPA and PAI2T; (vii) the pyrroloquinoline quinone (PQQ)-dependent glucose dehydrogenase (GLCDpp) was removed from the model; (viii) pyruvate formate lyase (PFL) and 2-oxobutanoate formate lyase (OBTFL) were deactivated only under aerobic conditions. We further removed the ATP maintenance reaction (ATPM) due to the fact that, rather than estimating growth rate, we used FBA to estimate the maximal yield. We used the model with these modifications as a ‘wild-type’ reference.

The Gnd reaction was changed to be reversible as part of the GED cycle/shunt; phosphoribulokinase (PRK) and Rubisco (RBPC) reactions were added to create the RuBP cycle/shunt. In order to use hydrogen as a proxy electron donor, hydrogen dehydrogenase reaction (H2DH, irreversible) was added to the model. The GED cycle and RuBP cycle were analyzed by assuming CO_2_ as the sole (unconstrained) carbon source, with hydrogen import as a constrained source of electrons and energy. Given that the GAP dehydrogenase (GAPDH) of the RuBP cycle is canonically considered NADPH-dependent, the according reaction of the model was modified to utilize this co-factor (only for analysis of the cycles, not the shunts). We note that this removes a positive bias, which would have caused yields of the GED cycle to be estimated higher. To analyze yields of the GED shunt and RuBP shunt, these linear variants were created by blocking the reactions of phosphofructokinase (PFK), S7P-reacting phosphofructokinase (PFK_3), fructose 6-phosphate aldolase (F6PA), glucose 6-phosphate dehydrogenase (G6PDH2r), and fructose-bisphosphatase (FBP). Then, xylose or glucose were assumed as constrained carbon source together with unconstrained CO_2_ (similar to ref. ^12^).The full code, including changes to the model and the new reactions of each production route can be found at https://gitlab.com/elad.noor/path-designer/-/tree/master/co2_fixation/FBA.

### Strains and genomic modifications

All strains used in this study are listed in Table 2. Gene deletions and growth experiments were performed in strains derived from *E.coli* SIJ488 ^76^, a strain derived from wild-type *E.coli* MG1655. The SIJ488 strain contains inducible genes for λ-Red recombineering (Red-recombinase and flippase) integrated into its genome to increase ease-of-use for multiple genomic modifications ^76^. Gene deletion strains were either taken from previous studies (Table 2) or generated by P1 phage transduction as described before ^77^. Strains from the Keio collection carrying single gene deletions with a kanamycin-resistance gene (KmR) as selective marker were used as donor strains ^78^. Strains that had acquired the desired deletion were selected by plating on appropriate antibiotics (Kanamycin, Km) and confirmed by determining the size of the respective genomic locus via PCR (oligonucleotide sequences shown in Supplementary Table 2). To remove the selective marker, flippase was induced in a fresh culture grown to OD_600_ ~ 0.2 by adding 50mM L-Rhamnose and cultivating for ~4h at 30°C. Loss of the antibiotic resistance was confirmed by identifying individual colonies that only grew on LB in absence of the respective antibiotic and by PCR of the genomic locus.

**Table 2.**
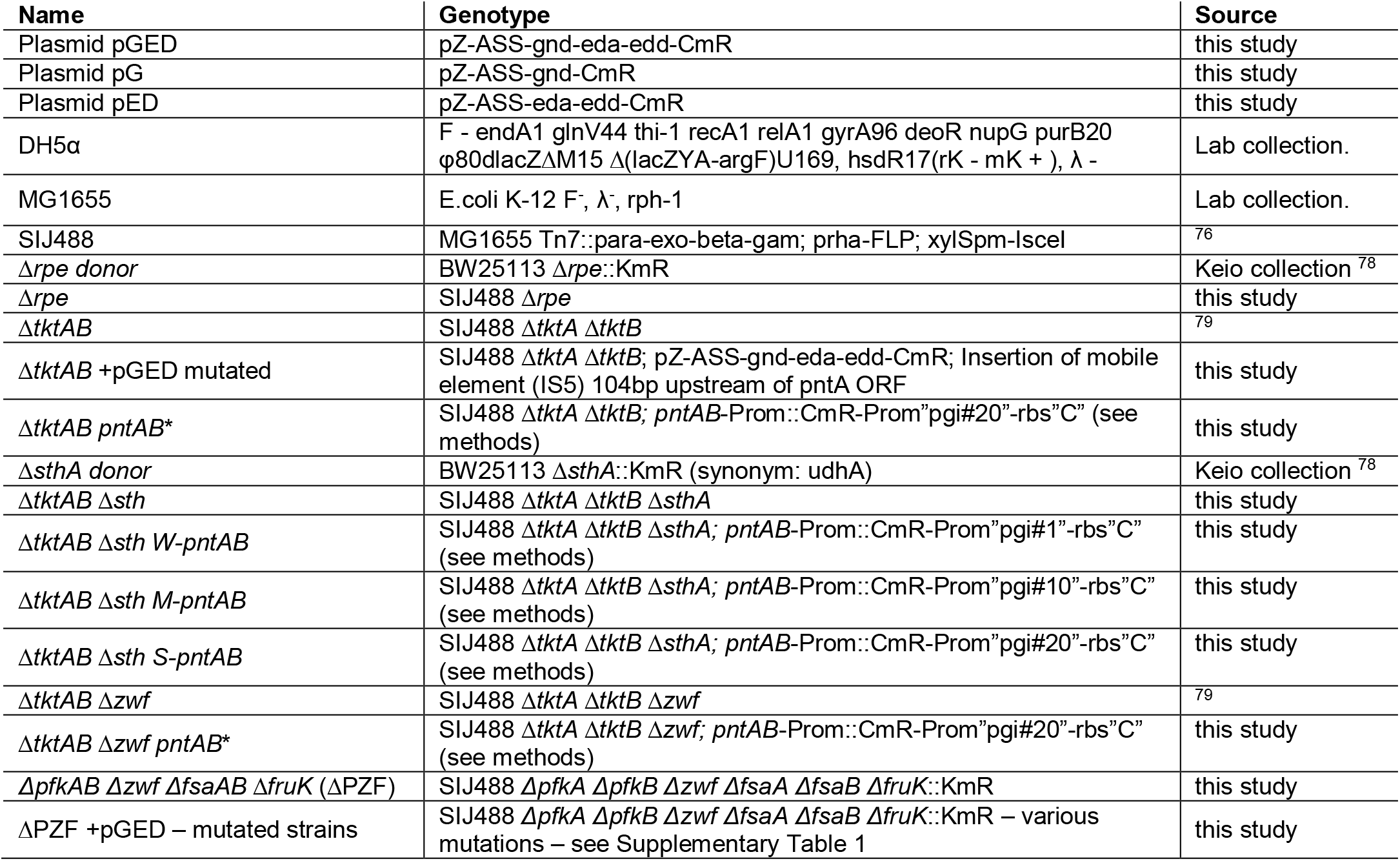
List of strains and expression plasmids used in this study. KmR and CmR denote kanamycin or chloramphenicol resistance markers, respectively.

For genomic overexpression of the *pntAB* operon, its promoter region was edited using a method based on λ-Red recombineering ^77^. We replaced the native *pntAB* promoter region (spanning 443bp upstream of the *pntA* start codon) by a weak (pgi#1), moderate (pgi#10), or strong (pgi#20) constitutive promoter ^42, 78^ and a medium-strength ribosome binding site (RBS “C”: AAGTTAAGAGGCAAGA ^79^), downstream of a CmR-cassette for selection (for resulting sequence, see Supplementary Table 2). For this purpose, the CmR cassette was amplified from plasmid pKD3 ^77^ with the primers CmR-1 and CmR-2 followed by overlap extension PCR to combine it with the promoter and RBS amplified from plasmid pGED with the primers PromW-Fwd, PromM-Fwd or PromS-Fwd, respectively, and pntA-Prom2. The construct was inserted into a pJET1.2 cloning-vector (Thermo Scientific, Dreieich, Germany) and confirmed by Sanger sequencing. Linear dsDNA donors for λ-Red recombineering were generated by amplification with primers pntA-Prom1 and pntA-Prom2. DNA (400ng) was transformed into the desired strains by electroporation after fresh culturing to OD_600_ ~ 0.3 and induction of recombinase enzymes by addition of 15mM L-Arabinose for 45 minutes. Confirmation of the engineered promoter region, ancestral background deletions and removal of the selective marker were performed as described above for the P1 transduction method. The *pntAB* promoter locus was additionally verified by Sanger sequencing (LGC Genomics, Berlin, DE) after amplification with pntA-V1 and pntA-V2.

### Construction of pGED, pG, and pED vectors

Cloning was carried out in *E. coli* DH5α. The native *E.coli* genes encoding 6-phosphogluconate dehydrogenase (*gnd*, UniProt: P00350), KHG/KDPG aldolase (*eda*, Uniprot: P0A955) and phosphogluconate dehydratase (*edd*, UniProt: P0ADF6) were amplified from *E.coli* MG1655 genomic DNA with high-fidelity Phusion Polymerase (Thermo Scientific, Dreieich, Germany) using primers listed in Supplementary Table 2. Silent mutations were introduced to remove relevant restriction sites in *gnd* (C292T to remove a PstI site) and *eda* (G196A to remove a PvuI site and C202T to remove a PstI site). Assembly of synthetic operons and expression plasmids was performed as described before ^47, 80^. In brief, genes were first inserted individually into a “pNivC” vector ^80^ downstream of a ribosomal binding site (RBS “C”, AAGTTAAGAGGCAAGA). For synthetic operons, multiple genes were assembled in pNivC vectors using BioBrick restriction enzymes (Fast-Digest: BcuI, XhoI, SalI, NheI; Thermo Scientific, Dreieich, Germany). The generated operons were excised from the pNivC vector by restriction with EcoRI and NheI (Fast Digest, Thermo Scientific, Dreieich, Germany) and inserted into a pZ-ASS vector ^47^ (p15**A** medium-copy origin of replication, **s**treptomycin resistance for expression under the control of the constitutive **s**trong promoter “*pgi* #20 ^81^). The order of genes in the operons was *gnd, eda, edd* for pGED; and *eda, edd* for pED. Constructed vectors were confirmed by Sanger sequencing (LGC Genomics, Berlin, DE). The software Geneious 8 (Biomatters, New Zealand) was used for *in silico* cloning and sequence analysis.

### Culture conditions & growth experiments

For routine culturing of *E. coli* strains, LB medium was used (5 g/l yeast extract, 10 g/l tryptone, 10 g/l NaCl). Antibiotics were added when appropriate at the following concentrations: Kanamycin 50 μg/ml; Chloramphenicol 30 μg/ml; Ampicillin 100 μg/ml; Streptomycin 100 μg/ml. Growth assays were performed in M9 minimal medium (47.8 mM Na_2_HPO_4_, 22 mM KH_2_PO_4_, 8.6 mM NaCl, 18.7 mM NH_4_Cl, 2 mM MgSO_4_ and 100 μM CaCl_2_), supplemented with trace elements (134 μM EDTA, 31 μM FeCl_3_·6H_2_O, 6.2 μM ZnCl_2_, 0.76 μM CuCl_2_·2H_2_O, 0.42 μM CoCl_2_-2H_2_O, 1.62 μM H_3_BO_3_, 0.081 μM MnCl_2_·4H_2_O). Carbon sources were added as described in the text at a concentration of 20 mM. No antibiotics were used in growth experiments, except in precultures. When elevated CO_2_ was required, cultures were grown in an orbital shaker set to maintain 37°C and an atmosphere of 20% CO_2_ mixed with air. Growth on strain deleted in *tktAB* required further supplementation of “E4P Supplements” ^36, 40^: 1 mM shikimic acid, 1 μM pyridoxine, 250 μM tyrosine, 500 μM phenylalanine, 200 μM tryptophan, 6 μM 4-aminobenzoic acid, 6 μM 4-hydroxybenzoic acid and 50 μM 2,3-dihydroxybenzoic acid.

Precultures for growth experiments were generally grown in M9 medium with 20 mM gluconate as carbon source (“relaxing conditions”). Antibiotics were added to the precultures if appropriate but omitted for growth experiments. Cells from the preculture were washed three times in M9 medium without carbon source and inoculated to a starting OD_600_ of 0.02 into M9 media with the final carbon sources as detailed in the text. 96-well plates (Nunclon Delta Surface, Thermo Scientific, Dreieich, Germany) were filled with 150μl culture and covered with 50 μl mineral oil (Merck, Darmstadt, Germany) to avoid evaporation while allowing gas exchange. Aerobic growth was monitored in technical duplicates or triplicates at 37°C in a BioTek Epoch 2 Microplate Spectrophotometer (BioTek, Bad Friedrichshall, Germany) by absorbance measurements (600nm) of each well every ~10 minutes with intermittent orbital and linear shaking. Blank measurements were subtracted and OD_600_ measurements were normalized to “cuvette OD_600_” values with the following equation previously established empirically for the instruments: OD_cuvette_ = OD_plate_/0.23. When elevated CO_2_ was required, the atmosphere was maintained at 20% CO_2_ mixed with 80% air by placing the plate reader inside a Kuhner ISF1-X incubator shaker (Kuhner, Birsfelden, Switzerland).

### Isolation and sequencing of a ΔtktAB +pGED mutant capable of growing via the GED shunt

Tube cultures (batch growth) of 4 mL selective minimal medium (M9 + E4P supplements + 20mM xylose) were inoculated to an OD_600_ of 0.05 (~1.5×10^7^ cells) and monitored during prolonged incubation at 37° and 20% CO_2_ (up to 3 weeks). Several cultures reached OD_600_ values above 1.0 after ~2 weeks. Single colonies were isolated from these cultures by dilution streak from liquid cultures onto LB medium with chloramphenicol (to maintain the pGED plasmid). Individual clones were then re-assayed for immediate growth (observable OD_600_ increase within 48h) on selective liquid minimal medium (M9+E4Ps+Xylose+20% CO_2_).

Genomic DNA was extracted using the GeneJET genomic DNA purification Kit (Thermo Scientific, Dreieich, Germany) from 2×10^9^ cells of an overnight culture in LB medium supplied with chloramphenicol (to maintain the pGED plasmid). Construction of PCR-free libraries for single-nucleotide variant detection and generation of 150 bp paired-end reads on an Illumina HiSeq 3000 platform were performed by the Max-Planck Genome Centre (Cologne, DE). Reads were mapped to the reference genome of *E.coli* MG1655 ^82^ (GenBank accession no. U00096.3) using the software Geneious 8 (Biomatters, New Zealand). Using algorithms supplied by the software package, we identified single-nucleotide variants (with >50% prevalence in all mapped reads) and searched for regions with coverage deviating more than 2 standard deviations from the global median coverage. Confirmation of the *pntA* promoter locus in the Δ*tkt* + pGED mutant was performed by Sanger Sequencing of a PCR product from amplification of the respective locus with high-fidelity Phusion Polymerase (Thermo Scientific, Dreieich, Germany). Sanger sequencing was performed by LGC Genomics (Berlin, DE).

### Expression analysis by reverse transcriptase quantitative PCR (RT-qPCR)

In order to determine mRNA levels, total RNA was extracted from growing cells in exponential phase (OD_600_ 0.5-0.6) on M9 minimal medium with 20mM carbon source (gluconate or xylose, and E4P supplements) in presence of 20% CO_2_. Total RNA was purified with the RNeasy Mini Kit (Qiagen, Hilden, Germany) as instructed by the manufacturer. In brief, ~5×10^8^ cells (1 ml of OD_600_ 0.5) were mixed with 2 volumes of RNAprotect Bacteria Reagent (Qiagen, Hilden, Germany) and pelleted, followed by enzymatic lysis, on-column removal of genomic DNA with RNase-free DNase (Qiagen, Hilden, Germany) and spin-column-based purification of RNA. Integrity and concentration of the isolated RNA were determined by NanoDrop and gel electrophoresis. Reverse transcription to synthesize cDNA was performed on 1 μg RNA with the qScript cDNA Synthesis Kit (QuantaBio, Beverly, MA USA). Quantitative real-time PCR was performed in technical triplicates using the Maxima SYBR Green/ROX qPCR Master Mix (Thermo Scientific, Dreieich, Germany). An input corresponding to 3.125 ng total RNA was used per reaction. Non-specific amplification products were excluded by melting curve analysis. The gene encoding 16S rRNA (*rrsA*) was chosen as a well-established reference transcript for expression normalization ^83^. Two alternative primer pairs for amplification of *pntA* were tested and the primer pair showing highest specificity and amplification efficiency was chosen (Supplementary Table 2). Negative control assays with direct input of RNA (without previous reverse transcription) confirmed that residual genomic DNA contributed to less than 5% of the signal (ΔCt between +RT/-RT samples >4 for all). Differences in expression levels were calculated according to the 2^-ΔΔCT^ method ^84, 85^. Reported data represents the 2^-ΔΔCT^ value that was calculated for each sample individually relative to the first biological wild-type replicate.

### Stationary ^13^C isotopic labelling of proteinogenic amino acids

For isotope tracing, cells were cultured in 3 mL M9 medium supplied with the labeled/unlabeled carbon sources described in the main text. For ^13^CO_2_ labeling, the experiment was performed in a 10 L desiccator that was first purged twice of the contained ambient air with a vacuum pump and refilled with an atmosphere of 80% air and 20% ^13^CO_2_ (Cambridge Isotope Laboratories Inc., MA USA). All cultures were inoculated to an OD_600_ 0.02 and grown at 37°C until stationary phase. Then, ~10^9^ cells (1 mL of culture with OD_600_ = 1) were pelleted, washed once with ddH_2_O and hydrolyzed in 1 mL hydrochloric acid (6M) at 95°C for a duration of 24h. Subsequently, the acid was evaporated by heating at 95°C and the hydrolyzed biomass was re-suspended in ddH_2_O.

Hydrolyzed amino acids were separated using ultra-performance liquid chromatography (Acquity, Waters, Milford, MA, USA) using a C18-reversed-phase column (Waters, Eschborn, Germany) as previously described ^86^. Mass spectra were acquired using an Exactive mass spectrometer (Thermo Scientific, Dreieich, Germany). Data analysis was performed using Xcalibur (Thermo Scientific, Dreieich, Germany). Prior to analysis, amino-acid standards (Merck, Darmstadt, Germany) were analyzed under the same conditions in order to determine typical retention times.

### Purification and kinetic characterization of *E.coli* Gnd

Proteins were expressed from *E.coli* BL21-AI strains (Invitrogen) carrying appropriate plasmids for expression of *E.coli* Gnd or *E.coli* RpiA (ribose-5-phosphate isomerase) which were taken from the ASKA collection ^87^. Expression was induced overnight at 30°C in TB medium (24 g/L yeast extract, 12 g/L tryptone, 4 mL/L glycerol, 17 mM KH_2_PO_4_, 72 mM K_2_HPO_4_) by addition of 0.5 mM IPTG and 2.5 mM arabinose upon reaching an OD_600_ of 1. Cells were lysed in 500 mM NaCl, 20 mM Tris-HCl pH 6.9 by sonication. After centrifugation (1h at 30,000 *g*), proteins were purified on an ÄKTA start system (GE Healthcare) by HisTrap Purification (GE Healthcare, Illinois, USA) as instructed by the manufacturer, using a wash step with 18% Buffer B (500mM NaCl, 20mM TrisHCl pH 6.9, 500mM imidazole). Desalting was performed in 100mM NaCl, 20 mM Tris-HCl pH 6.9 and enzymes were stored at −20°C in desalting buffer with 20% glycerol.

Kinetic assays were carried out on a Carry-60 UV-vis spectrometer (Agilent, Ratingen, Germany) at 30°C using a 1 mm quartz cuvette (Hellma). All assays were carried out in 100 mM TrisHCl buffer at pH 8 following consumption or production, respectively, of NADPH at 340 nm (ε_340 nm_ = 6.2 cm^-1^ mM^-1^). The reductive carboxylation parameters for Gnd were determined with assays containing 2.4 mM NADPH, 16 mM ribose 5-phosphate, 1.5 M KHCO3 and 140 μM RpiA (with varying concentrations of the substrate under investigation). The assays were preincubated for 2 min and started with the addition of 750 nM of freshly diluted Gnd. Carbonic anhydrase was used to confirm that CO_2_ equilibration was not rate-limiting in these assays. Isomerization of ribose 5-phosphate to ribulose 5-phosphate was confirmed not to be rate-limiting. The concentration of ribulose 5-phosphate was calculated from the equilibrium constant of the isomerization reaction: Keq = 0.458 (eQuilibrator ^24^). The kinetic parameters of the oxidative decarboxylation were determined with assays containing either 800 μM NADP^+^ (for 6-phosphogluconate parameters) or 200 μM 6-phosphogluconate (for NADP+ parameters). Assays were started with the addition of 7.5 nM Gnd. All points were measured in triplicates and each Michaelis-Menten curve was determined using at least 15 measurements.

## Acknowledgements

The authors thank Änne Michaelis for assistance with LC-MS analysis of amino acids; Nicole Paczia, Stefano Donati and Hannes Link for metabolite analysis; Selcuk Aslan for assistance with expression analysis; Lorenz Heck for assistance in molecular biology work; and Charlie Cotton and Nico Claassens for critical reading of the manuscript. This study was funded by the Max Planck Society.

## Author Contributions

S.N.L. and A.B.-E. conceived the study.

A.S., B.D., S.N.L., and A.B.-E. designed the experiments.

A. S., B.D., P.W., and S.N.L. performed the *in vivo* experiments.

E.N. and H.H. performed the *in silico* experiments.

B. V., and T.E. performed the *in vitro* experiments.

A.S., B.D., S.N.L., and A.B.-E. analyzed the results.

A.S., B.D., E.N., S.N.L., and A.B.-E. wrote the manuscript with contributions from all authors.

## Data Availability

Accession codes for protein sequences and structures are shown in the text. The data that support the findings of this study are available from the corresponding authors upon request.

## Code Availability

The code used in this study can be found at https://gitlab.com/elad.noor/path-designer/.

## References

1. Hugler, M. & Sievert, S.M. Beyond the Calvin cycle: autotrophic carbon fixation in the ocean. Ann Rev Mar Sci 3, 261–289 (2011).

2. Berg, I.A. Ecological Aspects of the Distribution of Different Autotrophic CO2 Fixation Pathways. Appl Environ Microbiol 77, 1925–1936 (2011).

3. Bar-Even, A., Noor, E. & Milo, R. A survey of carbon fixation pathways through a quantitative lens. J Exp Bot 63, 2325–2342 (2012).

4. Claassens, N.J., Sousa, D.Z., Dos Santos, V.A., de Vos, W.M. & van der Oost, J. Harnessing the power of microbial autotrophy. Nat Rev Microbiol 14, 692–706 (2016).

5. Satanowski, A. & Bar-Even, A. A one-carbon path for fixing CO2. EMBO Rep 21, e50273 (2020).

6. Antonovsky, N. et al. Sugar Synthesis from CO2 in Escherichia coli. Cell 166, 115–125 (2016).

7. Gleizer, S. et al. Conversion of Escherichia coli to Generate All Biomass Carbon from CO2. Cell 179, 1255–1263 e1212 (2019).

8. Gassler, T. et al. The industrial yeast Pichia pastoris is converted from a heterotroph into an autotroph capable of growth on CO2. Nat Biotechnol 38, 210–216 (2020).

9. Mattozzi, M., Ziesack, M., Voges, M.J., Silver, P.A. & Way, J.C. Expression of the sub-pathways of the Chloroflexus aurantiacus 3-hydroxypropionate carbon fixation bicycle in E. coli: Toward horizontal transfer of autotrophic growth. Metab Eng 16, 130–139 (2013).

10. Braakman, R. & Smith, E. The emergence and early evolution of biological carbon-fixation. PLoS Comput Biol 8, e1002455 (2012).

11. Erb, T.J. & Zarzycki, J. A short history of RubisCO: the rise and fall (?) of Nature’s predominant CO2 fixing enzyme. Curr Opin Biotechnol 49, 100–107 (2018).

12. Hadicke, O., von Kamp, A., Aydogan, T. & Klamt, S. OptMDFpathway: Identification of metabolic pathways with maximal thermodynamic driving force and its application for analyzing the endogenous CO2 fixation potential of Escherichia coli. PLoS Comput Biol 14, e1006492 (2018).

13. Monk, J.M. et al. iML1515, a knowledgebase that computes Escherichia coli traits. Nat Biotechnol 35, 904–908 (2017).

14. Noor, E. et al. Pathway thermodynamics highlights kinetic obstacles in central metabolism. PLoS Comput Biol 10, e1003483 (2014).

15. Levitt, M.D. Volume and composition of human intestinal gas determined by means of an intestinal washout technic. N Engl J Med 284, 1394–1398 (1971).

16. Montalto, M., Di Stefano, M., Gasbarrini, A. & Corazza, G.R. Intestinal gas metabolism. Dig Liver Dis Supp 3, 27–29 (2009).

17. Hesslinger, C., Fairhurst, S.A. & Sawers, G. Novel keto acid formate-lyase and propionate kinase enzymes are components of an anaerobic pathway in *Escherichia coli* that degrades L-threonine to propionate. Mol Microbiol 27, 477–492 (1998).

18. Bar-Even, A., Noor, E., Flamholz, A. & Milo, R. Design and analysis of metabolic pathways supporting formatotrophic growth for electricity-dependent cultivation of microbes. Biochim Biophys Acta 1827, 1039–1047 (2013).

19. Cotton, C.A., Claassens, N.J., Benito-Vaquerizo, S. & Bar-Even, A. Renewable methanol and formate as microbial feedstocks. Curr Opin Biotechnol 62, 168–180 (2020).

20. Yishai, O., Bouzon, M., Doring, V. & Bar-Even, A. In Vivo Assimilation of One-Carbon via a Synthetic Reductive Glycine Pathway in Escherichia coli. ACS synthetic biology 7, 2023–2028 (2018).

21. Kim, S. et al. Growth of E. coli on formate and methanol via the reductive glycine pathway. Nat Chem Biol (2020).

22. Bar-Even, A. Formate Assimilation: The Metabolic Architecture of Natural and Synthetic Pathways. Biochemistry 55, 3851–3863 (2016).

23. Barenholz, U. et al. Design principles of autocatalytic cycles constrain enzyme kinetics and force low substrate saturation at flux branch points. eLife 6 (2017).

24. Flamholz, A., Noor, E., Bar-Even, A. & Milo, R. eQuilibrator--the biochemical thermodynamics calculator. Nucleic Acids Res 40, D770–775 (2012).

25. Kwon, Y.D., Kwon, O.H., Lee, H.S. & Kim, P. The effect of NADP-dependent malic enzyme expression and anaerobic C4 metabolism in Escherichia coli compared with other anaplerotic enzymes. J Appl Microbiol 103, 2340–2345 (2007).

26. Zelle, R.M., Harrison, J.C., Pronk, J.T. & van Maris, A.J. Anaplerotic role for cytosolic malic enzyme in engineered *Saccharomyces cerevisiae* strains. Appl Environ Microbiol 77, 732–738 (2011).

27. Pound, K.M. et al. Substrate-enzyme competition attenuates upregulated anaplerotic flux through malic enzyme in hypertrophied rat heart and restores triacylglyceride content: attenuating upregulated anaplerosis in hypertrophy. Circ Res 104, 805–812 (2009).

28. Kanao, T., Kawamura, M., Fukui, T., Atomi, H. & Imanaka, T. Characterization of isocitrate dehydrogenase from the green sulfur bacterium Chlorobium limicola. A carbon dioxide-fixing enzyme in the reductive tricarboxylic acid cycle. Eur J Biochem 269, 1926–1931 (2002).

29. Igamberdiev, A.U. & Gardestrom, P. Regulation of NAD- and NADP-dependent isocitrate dehydrogenases by reduction levels of pyridine nucleotides in mitochondria and cytosol of pea leaves. Biochim Biophys Acta 1606, 117–125 (2003).

30. Villet, R.H. & Dalziel, K. The nature of the carbon dioxide substrate and equilibrium constant of the 6-phosphogluconate dehydrogenase reaction. Biochem J 115, 633–638 (1969).

31. Villet, R.H. & Dalziel, K. Studies of 6-phosphogluconate dehydrogenase from sheep liver. 1. Kinetics of the reductive carboxylation reaction. Eur J Biochem 27, 244–250 (1972).

32. Silverberg, M. & Dalziel, K. 6-Phospho-D-gluconate dehydrogenase from sheep liver. Methods Enzymol 41, 214–220 (1975).

33. Berdis, A.J. & Cook, P.F. Overall kinetic mechanism of 6-phosphogluconate dehydrogenase from Candida utilis. Biochemistry 32, 2036–2040 (1993).

34. Hanau, S., Montin, K., Cervellati, C., Magnani, M. & Dallocchio, F. 6-Phosphogluconate dehydrogenase mechanism: evidence for allosteric modulation by substrate. J Biol Chem 285, 21366–21371 (2010).

35. Flamholz, A.I. et al. Revisiting trade-offs between rubisco kinetic parameters. Biochemistry 58, 3365–3376 (2019).

36. He, H., Edlich-Muth, C., Lindner, S.N. & Bar-Even, A. Ribulose Monophosphate Shunt Provides Nearly All Biomass and Energy Required for Growth of E. coli. ACS synthetic biology 7, 1601–1611 (2018).

37. Meyer, F. et al. Methanol-essential growth of Escherichia coli. Nature communications 9, 1508 (2018).

38. Lyngstadaas, A., Sprenger, G.A. & Boye, E. Impaired growth of an Escherichia coli rpe mutant lacking ribulose-5-phosphate epimerase activity. Biochim Biophys Acta 1381, 319–330 (1998).

39. Chen, C.T. et al. Synthetic methanol auxotrophy of Escherichia coli for methanol-dependent growth and production. Metab Eng 49, 257–266 (2018).

40. Zhao, G. & Winkler, M.E. An Escherichia coli K-12 tktA tktB mutant deficient in transketolase activity requires pyridoxine (vitamin B6) as well as the aromatic amino acids and vitamins for growth. J Bacteriol 176, 6134–6138 (1994).

41. Schada von Borzyskowski, L. et al. Replacing the Ethylmalonyl-CoA Pathway with the Glyoxylate Shunt Provides Metabolic Flexibility in the Central Carbon Metabolism of Methylobacterium extorquens AM1. ACS synthetic biology 7, 86–97 (2018).

42. Kroger, M. & Hobom, G. Structural analysis of insertion sequence IS5. Nature 297, 159–162 (1982).

43. Sauer, U., Canonaco, F., Heri, S., Perrenoud, A. & Fischer, E. The soluble and membrane-bound transhydrogenases UdhA and PntAB have divergent functions in NADPH metabolism of Escherichia coli. J Biol Chem 279, 6613–6619 (2004).

44. Lindner, S.N. et al. NADPH-Auxotrophic E. coli: A Sensor Strain for Testing in Vivo Regeneration of NADPH. ACS synthetic biology 7, 2742–2749 (2018).

45. Schnetz, K. & Rak, B. IS5: a mobile enhancer of transcription in Escherichia coli. Proc Natl Acad Sci U S A 89, 1244–1248 (1992).

46. Zhang, Z. & Saier, M.H., Jr. A novel mechanism of transposon-mediated gene activation. PLoS Genet 5, e1000689 (2009).

47. Wenk, S., Yishai, O., Lindner, S.N. & Bar-Even, A. An Engineering Approach for Rewiring Microbial Metabolism. Methods Enzymol 608, 329–367 (2018).

48. Christodoulou, D. et al. Reserve Flux Capacity in the Pentose Phosphate Pathway Enables Escherichia coli’s Rapid Response to Oxidative Stress. Cell Syst 6, 569–578 e567 (2018).

49. Ralser, M. et al. Dynamic rerouting of the carbohydrate flux is key to counteracting oxidative stress. Journal of biology 6, 10 (2007).

50. Mall, A. et al. Reversibility of citrate synthase allows autotrophic growth of a thermophilic bacterium. Science 359, 563–567 (2018).

51. Nunoura, T. et al. A primordial and reversible TCA cycle in a facultatively chemolithoautotrophic thermophile. Science 359, 559–563 (2018).

52. Bar-Even, A., Noor, E., Lewis, N.E. & Milo, R. Design and analysis of synthetic carbon fixation pathways. Proc Natl Acad Sci U S A 107, 8889–8894 (2010).

53. Schwander, T., Schada von Borzyskowski, L., Burgener, S., Cortina, N.S. & Erb, T.J. A synthetic pathway for the fixation of carbon dioxide in vitro. Science 354, 900–904 (2016).

54. Berg, I.A., Kockelkorn, D., Buckel, W. & Fuchs, G. A 3-hydroxypropionate/4-hydroxybutyrate autotrophic carbon dioxide assimilation pathway in Archaea. Science 318, 1782–1786 (2007).

55. Erb, T.J. et al. Synthesis of C5-dicarboxylic acids from C2-units involving crotonyl-CoA carboxylase/reductase: the ethylmalonyl-CoA pathway. Proc Natl Acad Sci U S A 104, 10631–10636 (2007).

56. Miller, T.E. et al. Light-powered CO2 fixation in a chloroplast mimic with natural and synthetic parts. Science 368, 649–654 (2020).

57. Bogorad, I.W., Lin, T.S. & Liao, J.C. Synthetic non-oxidative glycolysis enables complete carbon conservation. Nature 502, 693–697 (2013).

58. Aslan, S., Noor, E. & Bar-Even, A. Holistic bioengineering: rewiring central metabolism for enhanced bioproduction. Biochem J 474, 3935–3950 (2017).

59. Babel, W. The Auxiliary Substrate Concept: From simple considerations to heuristically valuable knowledge. Eng. Life Sci. 9, 285–290 (2009).

60. Guadalupe-Medina, V. et al. Carbon dioxide fixation by Calvin-Cycle enzymes improves ethanol yield in yeast. Biotechnology for biofuels 6, 125 (2013).

61. Zhuang, Z.Y. & Li, S.Y. Rubisco-based engineered Escherichia coli for in situ carbon dioxide recycling. Bioresour Technol 150, 79–88 (2013).

62. Tseng, I.T. et al. Exceeding the theoretical fermentation yield in mixotrophic Rubisco-based engineered Escherichia coli. Metab Eng 47, 445–452 (2018).

63. Herz, E. et al. The genetic basis for the adaptation of E. coli to sugar synthesis from CO2. Nature communications 8, 1705 (2017).

64. Bar-Even, A. Daring metabolic designs for enhanced plant carbon fixation. Plant science: an international journal of experimental plant biology 273, 71–83 (2017).

65. Chakravarty, J. & Brigham, C.J. Solvent production by engineered Ralstonia eutropha: channeling carbon to biofuel. Appl Microbiol Biotechnol 102, 5021–5031 (2018).

66. Mackinder, L.C. et al. A repeat protein links Rubisco to form the eukaryotic carbon-concentrating organelle. Proc Natl Acad Sci U S A 113, 5958–5963 (2016).

67. Mangan, N.M., Flamholz, A., Hood, R.D., Milo, R. & Savage, D.F. pH determines the energetic efficiency of the cyanobacterial CO2 concentrating mechanism. Proc Natl Acad Sci U S A 113, E5354–5362 (2016).

68. Stoffel, G.M.M. et al. Four amino acids define the CO2 binding pocket of enoyl-CoA carboxylases/reductases. Proc Natl Acad Sci U S A 116, 13964–13969 (2019).

69. Bernhardsgrutter, I. et al. Awakening the Sleeping Carboxylase Function of Enzymes: Engineering the Natural CO2-Binding Potential of Reductases. J Am Chem Soc 141, 9778–9782 (2019).

70. Cotton, C.A., Edlich-Muth, C. & Bar-Even, A. Reinforcing carbon fixation: CO2 reduction replacing and supporting carboxylation. Curr Opin Biotechnol 49, 49–56 (2018).

71. Kanehisa, M. & Goto, S. KEGG: kyoto encyclopedia of genes and genomes. Nucleic Acids Res 28, 27–30 (2000).

72. Ebrahim, A., Lerman, J.A., Palsson, B.O. & Hyduke, D.R. COBRApy: COnstraints-Based Reconstruction and Analysis for Python. BMC Syst Biol 7, 74 (2013).

73. Mendler, K. et al. AnnoTree: visualization and exploration of a functionally annotated microbial tree of life. Nucleic Acids Res 47, 4442–4448 (2019).

74. Bizouarn, T., van Boxel, G.I., Bhakta, T. & Jackson, J.B. Nucleotide binding affinities of the intact proton-translocating transhydrogenase from Escherichia coli. Biochim Biophys Acta 1708, 404–410 (2005).

75. He, H., Höper, R., Dodenhöft, M., Marlière, P. & Bar-Even, A. An optimized methanol assimilation pathway relying on promiscuous formaldehyde-condensing aldolases in E. coli. Metab Eng 60, 1–13 (2020).

76. Jensen, S.I., Lennen, R.M., Herrgard, M.J. & Nielsen, A.T. Seven gene deletions in seven days: Fast generation of Escherichia coli strains tolerant to acetate and osmotic stress. Scientific reports 5, 17874 (2015).

77. Thomason, L.C., Costantino, N. & Court, D.L. E. coli genome manipulation by P1 transduction. Curr Protoc Mol Biol Chapter 1, Unit 1 17 (2007).

78. Baba, T. et al. Construction of Escherichia coli K-12 in-frame, single-gene knockout mutants: the Keio collection. Mol Syst Biol 2, 2006–2008 (2006).

79. Krusemann, J.L. et al. Artificial pathway emergence in central metabolism from three recursive phosphoketolase reactions. FEBS J 285, 4367–4377 (2018).

80. Zelcbuch, L. et al. Spanning high-dimensional expression space using ribosome-binding site combinatorics. Nucleic Acids Res 41, e98 (2013).

81. Braatsch, S., Helmark, S., Kranz, H., Koebmann, B. & Jensen, P.R. Escherichia coli strains with promoter libraries constructed by Red/ET recombination pave the way for transcriptional fine-tuning. Biotechniques 45, 335–337 (2008).

82. Hayashi, K. et al. Highly accurate genome sequences of Escherichia coli K-12 strains MG1655 and W3110. Mol Syst Biol 2, 2006 0007 (2006).

83. Rocha, D.J., Santos, C.S. & Pacheco, L.G. Bacterial reference genes for gene expression studies by RT-qPCR: survey and analysis. Antonie Van Leeuwenhoek 108, 685–693 (2015).

84. Livak, K.J. & Schmittgen, T.D. Analysis of relative gene expression data using real-time quantitative PCR and the 2-ΔΔCT method. methods 25, 402–408 (2001).

85. Schmittgen, T.D. & Livak, K.J. Analyzing real-time PCR data by the comparative C T method. Nature protocols 3, 1101 (2008).

86. Giavalisco, P. et al. Elemental formula annotation of polar and lipophilic metabolites using 13C, 15N and 34S isotope labelling, in combination with high-resolution mass spectrometry. Plant J 68, 364–376 (2011).

87. Kitagawa, M. et al. Complete set of ORF clones of Escherichia coli ASKA library (a complete set of E. coli K-12 ORF archive): unique resources for biological research. DNA Res 12, 291–299 (2005).

