## Supplementary Information for "Awakening a latent carbon fixation cycle in *Escherichia coli*"

- Supplementary Figures S1-S5
- Supplementary Tables S1-S2
- Supplementary Data
- Supplementary Text

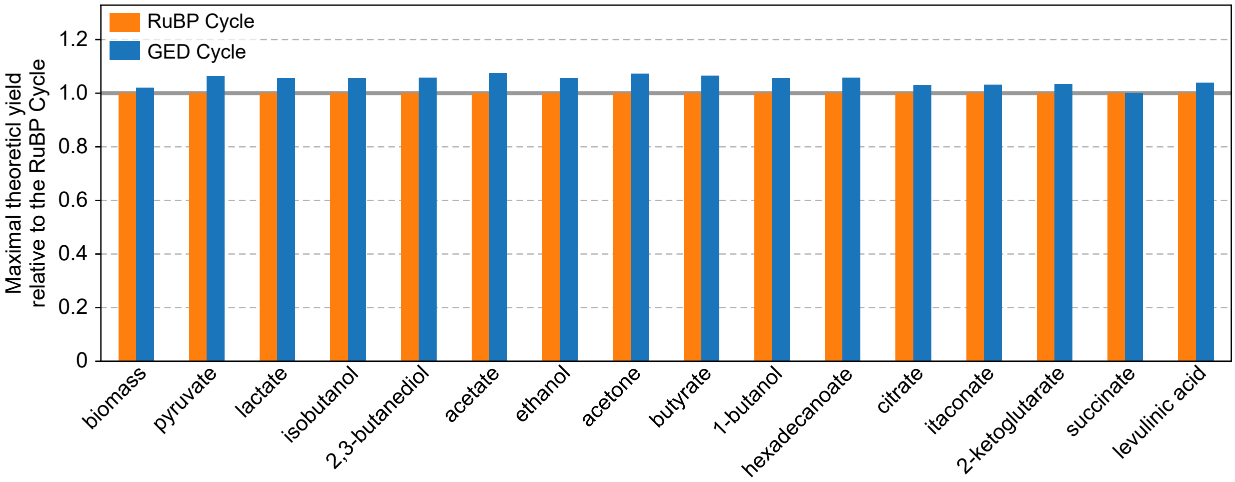


**Supplementary Figure S1:** The GED cycle is predicted to achieve higher yields of biomass and various commercially relevant products when compared to the RuBP cycle. Maximal theoretical yields of biomass and 15 selected products are shown, as calculated by Flux Balance Analysis. Presented values are normalized to the product yield from the RuBP cycle.

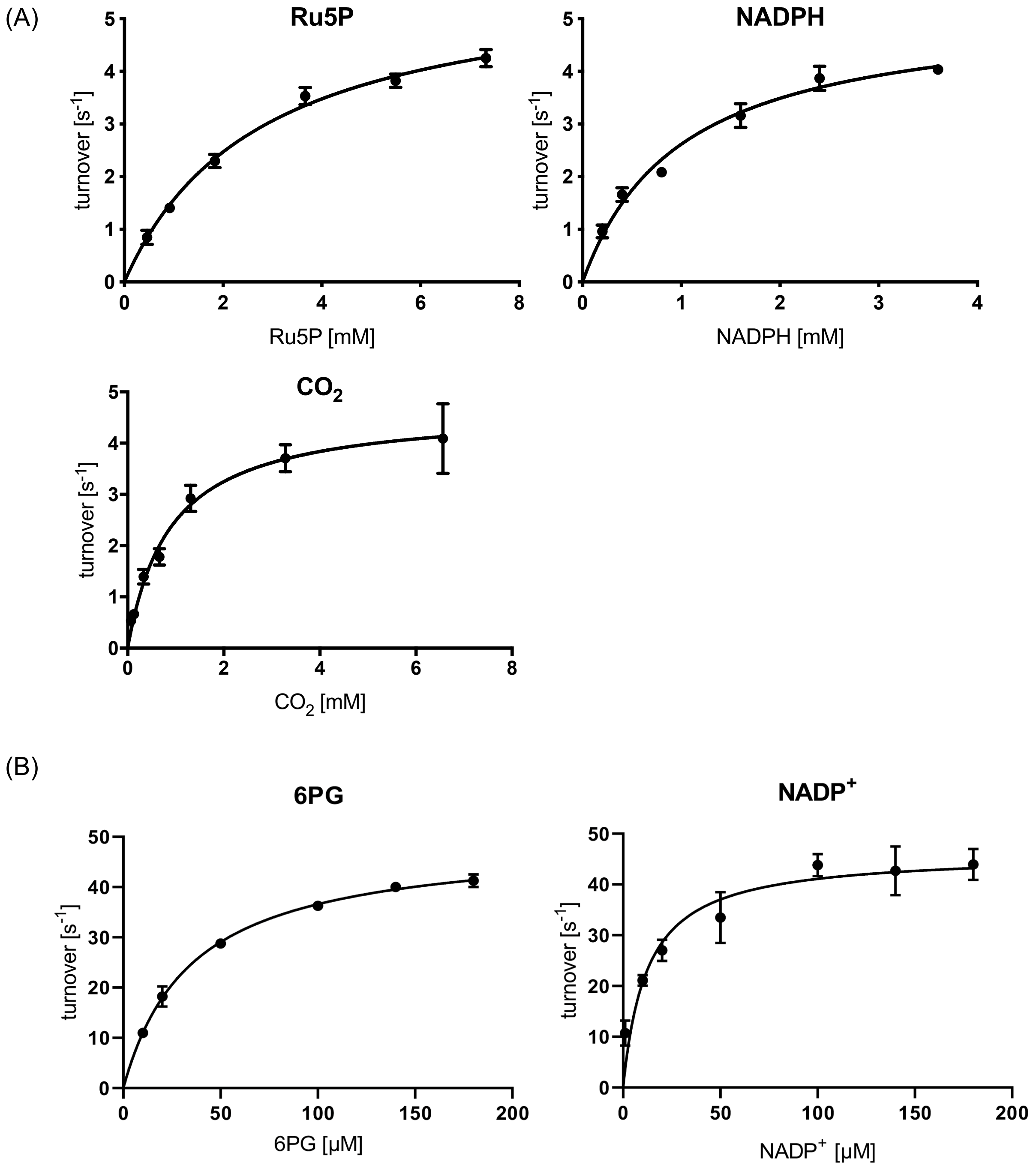


**Supplementary Figure S2:** Michaelis-Menten kinetics of (A) reductive carboxylation and (B) oxidative decarboxylation by *E.coli* Gnd. The data are summarized in the main text.

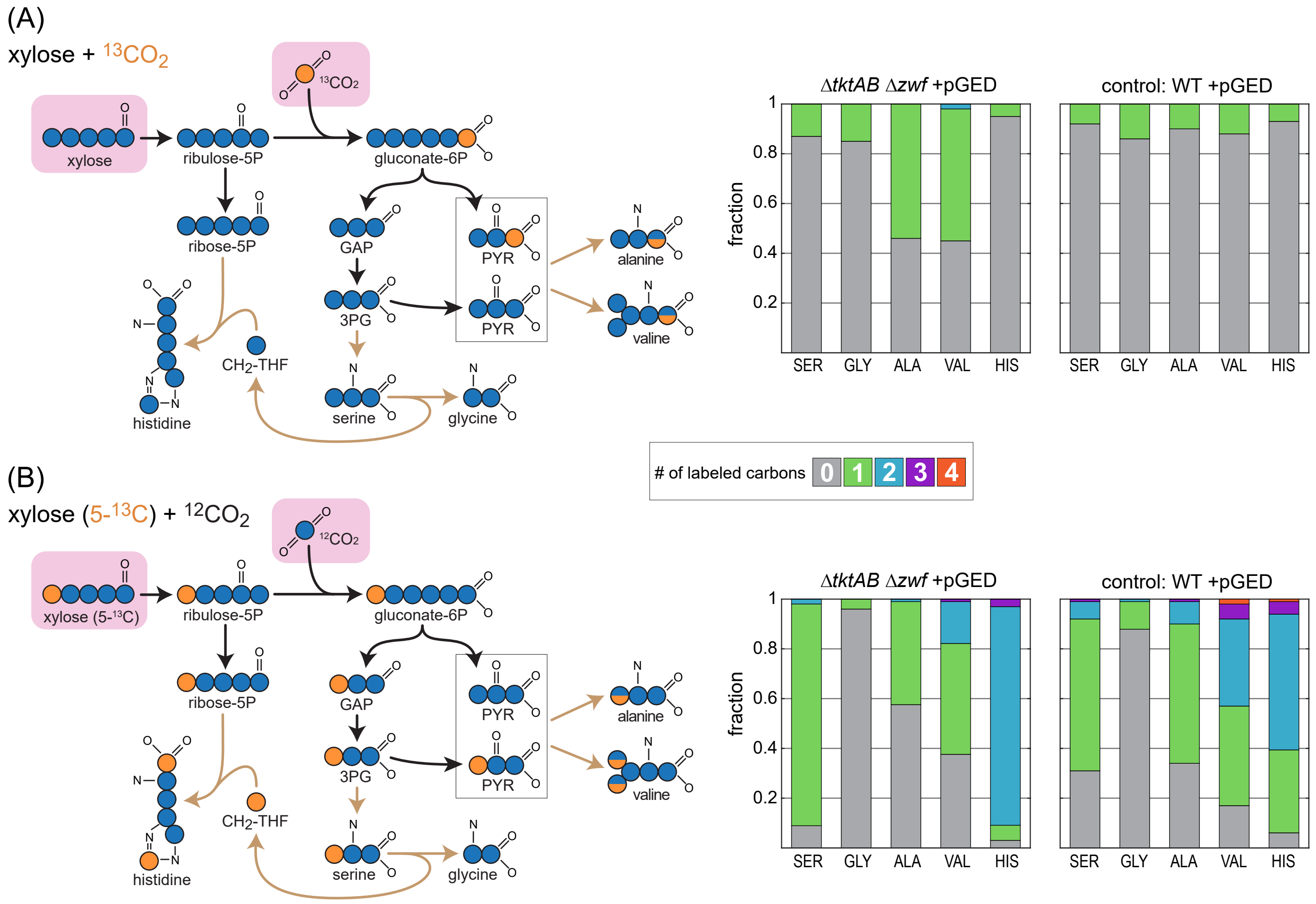


**Supplementary Figure S3:** $^{13}$C-labeling experiments confirm the operation of the GED shunt in the Δ*tktAB* (Δ*zwf*) strain. (A) Cultivation with unlabeled xylose + $^{13}$CO2. (B) Cultivation with xylose (5-$^{13}$C) + unlabeled $CO_2$. On the left, the expected labeling distributions for the respective feed-stocks are shown. Observed labeling fits the expected pattern and differs from the labeling in a WT strain, which serves as a control. Abbreviations: ALA, Alanine; GLY, Glycine; HIS, Histidine; SER, Serine; VAL, Valine.

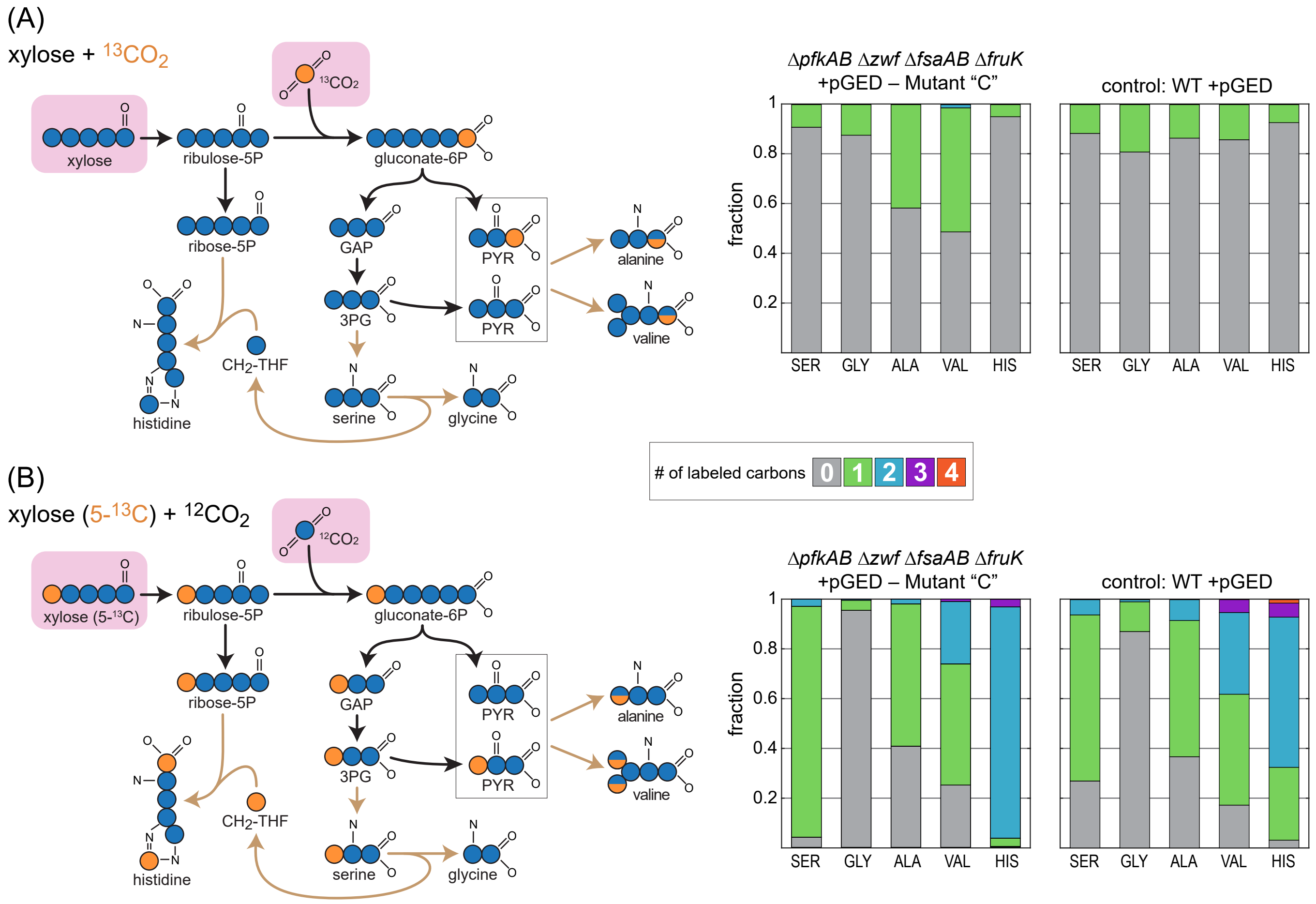


**Supplementary Figure S4:** $^{13}$C-labeling experiments confirm the operation of the GED shunt in a ΔPZF strain ("ΔPZF +pGED Mutant C1"). (A) Cultivation with unlabeled xylose + $^{13}$CO2. (B) Cultivation with xylose (5-$^{13}$C) + unlabeled $CO_2$. On the left, the expected labeling distributions for the respective feedstocks are shown. Observed labeling fits the expected pattern and differs from the labeling in a WT strain, which serves as a control. Abbreviations: ALA, Alanine; GLY, Glycine; HIS, Histidine; SER, Serine; VAL, Valine.

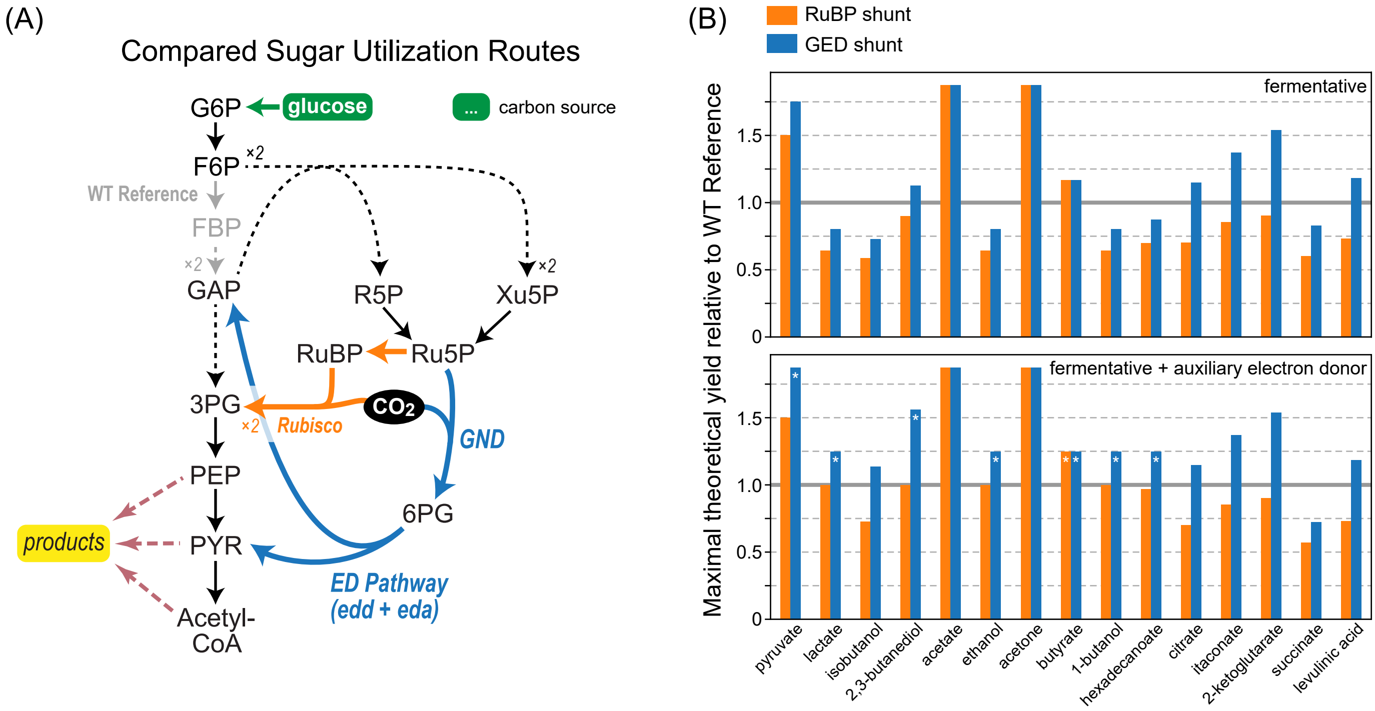


**Supplementary Figure S5:** Rerouting glucose fermentation via the GED shunt is expected to increase product yields. (A) An overview comparing glucose utilization routes: Canonical utilization (gray) via glycolysis (Embden-Meyerhof-Parnas pathway); the GED shunt (blue); and a linear pathway based on carboxylation by Rubisco ("RuBP shunt", orange). (B) Maximal theoretical yields of 15 fermentation products are shown with or without an additional electron donor, as calculated by Flux Balance Analysis. Presented values are normalized to the product yield from canonical sugar utilization (glycolysis). Most products are predicted to require the secretion of other organic compounds (e.g. acetate, formate) to achieve balancing of reducing equivalents or support ATP biosynthesis. Asterisks denote products that could be produced without byproducts.

**Supplementary Table 1:** Mutations identified in the evolved ΔPZF strains.

| Locus / Gene Name | Position (in NC_000913.3) | Annotation | Mutation Type | Reference bases | Variant bases | Description/Comments | Ancestral Strain (ΔpfkAB Δzwf ΔfsaAB) | Parental Strain (ΔPZF +pGED) ΔpfkAB Δzwf ΔfsaAB ΔfruK +pGED | ΔPZF +pGED - Mutant A1 | ΔPZF +pGED - Mutant A2 | ΔPZF +pGED - Mutant A3 | ΔPZF +pGED - Mutant B1 | ΔPZF +pGED - Mutant B2 | ΔPZF +pGED - Mutant B3 | ΔPZF +pGED - Mutant C1 | ΔPZF +pGED - Mutant C2 | ΔPZF +pGED - Mutant C3 |
|---|---|---|---|---|---|---|---|---|---|---|---|---|---|---|---|---|---|
| Cra | 88,283 | coding region (Q86*) | SNP | C | T | DNA-binding transcriptional dual regulator | | + | + | + | + | + | + | + | + | + | + |
| cydA | 771,121 | intergenic (-336nt) | IS element insertion | - | - | cytochrome bd-I ubiquinol oxidase subunit I | | + | + | + | + | + | + | + | + | + | + |
| rne | 1,142,413 | coding region (1954th nt) | Δ2 bp deletion / frameshift | TC | - | ribonuclease E | | | + | + | + | | | | | | |
| dhaM | 1,248,151 | coding region | IS element insertion | - | - | dihydroxyacetone kinase, phosphotransferase component | | + | + | + | + | + | + | + | + | + | + |
| clsA | 1,307,916 | coding region (D244N) | SNP | C | T | cardiolipin synthase A | | | + | + | + | | | | | | |
| flhD | 1,978,503 | intergenic (-400nt) | IS element insertion | - | - | DNA-binding transcriptional dual regulator | | + | + | + | + | + | + | + | + | + | + |
| gatC | 2,173,363 | coding region | Δ2 bp deletion | CC | - | galactitol-specific PTS enzyme IIC component | | + | + | + | + | + | + | + | + | + | + |
| ptsI | 2,535,329 | coding region (E422*) | SNP | G | T | phosphoenolpyruvate-protein phosphotransferase PTS enzyme I | | + | + | + | + | + | + | + | + | + | + |
| avtA | 3,740,351 | coding region (647th nt) | Δ2 bp deletion | TG | - | valine-pyruvate aminotransferase | | | | | | | | | + | | + |
| avtA | 3,740,466 | coding region (G254GG) | 3bp insertion | - | TGG | valine-pyruvate aminotransferase | | | | | | + | | + | | | |
| glpF / zapB | 4,118,275 | intergenic (-184nt) | IS element insertion | - | - | glycerol facilitator | | + | + | + | + | + | + | + | + | + | + |
| eptA | 4,334,799 | coding region (silent) | SNP | C | T | phosphoethanolamine transferase | | + | + | + | + | + | + | + | + | + | + |
| leuX (tRNA) | 4,496,406 | coding region (2nd nt) | SNP | C | T | tRNA-Leu(CAA) | | | | | | + | + | + | + | + | + |
| fimA | 4,543,603 | coding region (502nd nt) | IS element insertion | - | - | type 1 fimbriae (pili) major subunit | | | | | | | + | | | | |
| yjjY | 4,640,422 | coding region | IS element insertion | - | - | protein of unknown function | | + | + | + | + | + | + | + | + | + | + |
| | | | | | | | | | This strain is refered to as Mutant 'A' in the main text | | | This strain is refered to as Mutant 'B' in the main text | | | This strain is refered to as Mutant 'C' in the main text | | |
| | | | | | | | | | These three colonies were isolated from a single culture evolved for growth on xylose | | | These three colonies were isolated from a single culture evolved for growth on xylose | | | These three colonies were isolated from a single culture evolved for growth on xylose | | |

**Supplementary Table 2:** List of DNA oligo primers used in this study.

| Name | Function | Sequence (5'→3') |
|---|---|---|
| rpe-V1 | Verification of Δrpe | GTTTTACGCTGAGTCGCACGTTCCTG |
| rpe-V2 | | CGAAGGTCTGGTTGAGCGCCATATTCC |
| tktA-V1 | Verification of ΔtktA | ACAACAAATGTCAATACGCATATCGT |
| tktA-V2 | | GTTCCATGTACATGACGCGC |
| tktB-V1 | Verification of ΔtktB | CCACCTTCTCAGACGTTCCC |
| tktB-V2 | | GCTTTGACGGTCAGCGTTTT |
| zwf-V1 | Verification of Δzwf | GCACGAGGCCTGAAAGTGTA |
| zwf-V2 | | AAAGCAGTACAGTGCACCGT |
| sthA-V1 | Verification of ΔsthA (ΔudhA) | GTTTCTGTTTTGAAGCCGGGGC |
| sthA-V2 | | AAACAGACAAAGCAAAGGCCGC |
| gndA | Amplification of gnd from *E.coli* genomic DNA | GAATGCATCATCACCATCACCACTCCAAGCAACAGATCGGCGTAGTCGG |
| gndB | | GCATCGGTGATTTTTTGCAGGAACTGCGC |
| gndC | | GCGCAGTTCCTGCAAAAAATCACCGATGC |
| gndD | | CGCTAGCTCTAGATTAATCCAGCCATTCGGTATGGAACACACCTTC |
| edaA | Amplification of eda from *E.coli* genomic DNA | GAATGCATCATCACCATCACCACAAAAACTGGAAAACAAGTGCAGAATCAATCCTGACCAC |
| edaB | | GAACGGACCCGCAATCGCTTGCAGGGCTTTCAC |
| edaC | | GTGAAAGCCCTGCAAGCGATTGCGGGTCCGTTC |
| edaD | | CGCTAGCTCTAGATTACAGCTTAGCGCCTTCTACAGCTTCACG |
| eddA | Amplification of edd from *E.coli* genomic DNA | GAATGCATCATCACCATCACCACAATCCACAATTGTTACGCGTAACAAATCGAATCATTGAACG |
| eddB | | CGCTAGCTCTAGATTAAAAAGTGATACAGGTTGCGCCCTGTTCGG |
| q-rrsA1 | qPCR of *rrsA* (reference gene) | CTCTTGCCATCGGATGTGCCCA |
| q-rrsA2 | | CCAGTGTGGCTGGTCATCCTCTCA |
| q-pntA1 | qPCR of *pntA* | GCCAATCTGCAACAGTGCTC |
| q-pntA2 | | TTTTTGGCTGGATGGCAAGC |
| CmR-1 | Engineering of the pntAB promotor region | TAATACGACTCACTATAGGGCTCCATATGAATATCCTCCTTAG |
| CmR-2 | | AATTAACCCTCACTAAAGGGCGGAGCTGCTTCGAAGTTCCTA |
| PromW-Fwd | | GAGCCCTATAGTGAGTCGTATTATCCCTTTGATATTGCATCCCGCGTATATAATATG |
| PromM-Fwd | | GAGCCCTATAGTGAGTCGTATTAACCTATTGACAATTAAAGGCTAAAATGCTATAATTCCAC |
| PromS-Fwd | | GAGCCCTATAGTGAGTCGTATTAAATACTTGACATATCACTGTGATTCACATATAATATGCG |
| pntA-Prom1 | | GTACTGGTATTGTTATTAACGAGAAACGTGGCTGATTATTGCATTTAAACAATTAACCCTCACTAAAGGGCG |
| pntA-Prom2 | | GCAACACGGGTTTCATTGGTTAACCGTTCTCTTGGTATGCCAATTCGCATTCTTGCCTCTTAACTTTAAAGTTAAACAAAATTATTTCTATTA |
| pntA-V1 | Sequencing of pntA promotor region | AACCCAGTTTCAGCAGCTG |
| pntA-V2 | | CCAGGAGGGTGTTCTTAAGC |

Supplementary Data

Sequence of the genomic region upstream of the *pntA* gene in the isolated Δ*tktAB* + pGED mutant strain. Binding sites of the primers used for Sanger sequencing are in bold; coding sequence of the pntA gene is underlined; sequence of the IS5 mobile element is shown in red

**aacccagtttcagcagctg**ttccactgtttttggcgttgctgcaacacgggtttcattggttaaccgttct
cttggtatgccaattcgcatgatattcccttccatcggttttattgatgatggtttgcctgtgtcaggagc
cacacaagctgctcatgtacgagctaaatgttactccgttaaaataaattagggaaggtgcgaataagcgg
ggaaattcttctcggctgactcagtcatttcatttcttcatgtttgagccgatttttttctcccgtaaatgc
cttgaatcagcctatttagaccgtttcttcgccatttaaggcgttatccccagttttttagtgagatctctc
ccactgacgtatcatttggtccgcccgaaacaggttggccagcgtgaataacatcgccagttggttatcgt
ttttcagcaaccccttgtatctggctttcacgaagccgaactgtcgcttgatgatgcgaaatgggtgctcc
accctggcccggatgctggctttcatgtattcgatgttgatggccgttttgttcttgcgtggatgctgttt
caaggttcttaccttgccgggggcgctcggcgatcagccagtccacatccacctcggccagctcctcgcgct
gtggcgccccttggtagccggcatcggctgagacaaattgctcctctccatgcagcagattacccagctga
ttgaggtcatgctcgttggccgcggtggtgaccaggctgtgggtcaggccactcttggcatcgacaccaat
gtgggccttcatgccaaagtgccactgattgcctttcttggtctgatgcatctccggatcgcgttgctgct
ctttgttcttggtcgagctgggtgcctcaatgatggtggcatcgaccaaggtgccttgagtcatcatgacg
cctgcttcggccagccagcgattgatggtcttgaacaattggcgggccagttgatgctgctccagcaggtg
gcggaaattcatgatggtggtgcggtccggcaaggcgctatccagggataaccgggcaaacagacgcatgg
aggcgatttcgtacagagcatcttccatcgcgccatcgctcaggttgtaccaatgctgcatgcagtgaatg
cgtagcatggtttccagcggataaggtcgccggccattaccagccttggggtaaaacggctcgatgacttc
caccatgttttgccatggcagaatctgctccatgcgggacaagaaaatctcttttctggtctgacggcgct
tactgctgaattcactgtcggcgaaggtaagttgatgactcatgatgaaccctgttctatggctccagatg
acaaacatgatctcatatcagggacttgttcgcaccttcctaacaaacgccctataacgtactgaaaatta
**tgcctgtgatctagcgccaaaa**

Sequence of the construct (linear dsDNA) used for *pntA* promotor engineering with the λ Red recombinase method. Homologous regions for recombination are in bold lower-case; introduced promotor region is in upper-case; ribosome binding site + *pntA* start codon are underlined; CmR selection marker is shown in red.

**gcaacacgggtttcattggttaaccgttctcttggtatgccaattcgcat**TCTTGCCTCTTAACTTTAAAG
TTAAACAAAATTATTTCTATTAACTAGTGAATTCGGTCAGTGCGTCCTGCGCATATTATATGTGAATCACA
GTGATATGTCAAGTATTTAATACGACTCACTATAGGGCTCCATATGAATATCCTCCTTAGTTCCTATTCCG
AAGTTCCTATTCTCTAGAAAGTATAGGAACTTCGGCGCGCCTACCTGTGACGGAAGATCACTTCGCAGAAT
AAATAAATCCTGGTGTCCCTGTTGATACCGGGAAGCCCTGGGCCAACTTTTGGCGAAAATGAGACGTTGAT
CGGCACGTAAGAGGTTCCAACTTTCACCATAATGAAATAAGATCACTACCGGGCGTATTTTTTGAGTTGTC
GAGATTTTCAGGAGCTAAGGAAGCTAAAATGGAGAAAAAAATCACTGGATATACCACCGTTGATATATCCC
AATGGCATCGTAAAGAACATTTTGAGGCATTTCAGTCAGTTGCTCAATGTACCTATAACCAGACCGTTCAG
CTGGATATTACGGCCTTTTTAAAGACCGTAAAGAAAAATAAGCACAAGTTTTATCCGGCCTTTATTCACAT
TCTTGCCCGCCTGATGAATGCTCATCCGGAATTACGTATGGCAATGAAAGACGGTGAGCTGGTGATATGGG
ATAGTGTTCACCCTTGTTACACCGTTTTCCATGAGCAAACTGAAACGTTTTCATCGCTCTGGAGTGAATAC
CACGACGATTTCCGGCAGTTTCTACACATATATTCGCAAGATGTGGCGTGTTACGGTGAAAACCTGGCCTA
TTTCCCTAAAGGGTTTATTGAGAATATGTTTTTCGTCTCAGCCAATCCCTGGGTGAGTTTCACCAGTTTTG
ATTTAAACGTGGCCAATATGGACAACTTCTTCGCCCCCGTTTTCACCATGGGCAAATATTATACGCAAGGC
GACAAGGTGCTGATGCCGCTGGCGATTCAGGTTCATCATGCCGTTTGTGATGGCTTCCATGTCGGCAGAAT
GCTTAATGAATTACAACAGTACTGCGATGAGTGGCAGGGCGGGGCGTAAGGCGCGCCATTTAAATGAAGTT
CCTATTCCGAAGTTCCTATTCTCTAGAAAGTATAGGAACTTCGAAGCAGCTCCGCCCTTTAGTGAGGGTTA
ATT**gtttaaatgcaataatcagccacgtttctcgttaataacaataccagtac**

### Supplementary Text

#### 1 path-designer: an MILP-based algorithm for metabolic pathway design

We use a similar approach to the recently published OptMDFpathway method [1]. We set up a Mixed Integer Linear Problem (MILP) -based optimization problem which simultaneously looks for solutions that balance an objective reaction (here, 3 $CO_2 \rightarrow$ pyruvate), and maximize the Max-min Driving Force (MDF) [3] while simultaneously minimizing the number of reactions. In section 1.1, we lay out the changes we made to the set of reactions in the iML1515 model [2]. In section 1.2 we describe how the MILP problem is formulated. Finally, in section 1.3 we describe how we use the MILP framework to find also sub-optimal solutions and cover a large part of the feasible solution space of carbon fixation pathways.

##### 1.1 Alterations to the iML1515 model

We made a few changes to the genome-scale model of *E. coli* [2]:

- Removing all non-cytoplasmic reactions (i.e. exchange or transport reactions), except for exchange reactions of inorganic metabolites: protons, water, orthophosphate, ammonium, and oxygen.
- Removing all boundary reactions (i.e. sink reactions needed to allow certain co-factors to leave the system).
- Adding co-factor regenerating reactions: ADP $\rightarrow$ ATP, $NADP^+ \rightarrow$ NADPH, and $NAD^+ \rightarrow$ NADH.
- Replacing all flavoredoxins and thioredoxins with NADP †.
- Removing the Formate-tetrahydrofolate ligase reaction (FTHFLi) ‡.
- Add the objective reaction (OBJ): 3 $CO_2 \rightarrow$ pyruvate.
- Setting the bounds (the range of possible fluxes) of all remaining reactions to be between -10 and 10.

† We replaced all flavoredoxins and thioredoxins with NADP, since we do not have a good estimate of their reduction potential, and therefore the MDF for pathways using them was artificially high. We can assume that the electrons used for reducing $CO_2$ in the carbon fixation cycle ultimately have to pass through NADP, and therefore a simple solution was to replace the electron donor with NADPH. This way, we could keep the flavoredoxins/thioredoxins-dependent reactions in the model while having a more realistic estimate of their thermodynamics.

‡ We found that the reaction formate-tetrahydrofolate ligase (FTHFLi) appears in some of the solutions although the gene associated with this reaction is unknown. FTHFLi was thus excluded from our model altogether. Notably, removing this reaction does not significantly affect the space of solutions, because it can be easily replaced by GAR transformylase-T (GART) and the reverse reaction of Phosphoribosylglycinamide formyltransferase (GARFT).

| Compound | BiGG identifier | Concentration range |
|---|---|---|
| ATP | atp_c | 5 mM |
| ADP | adp_c | 0.5 mM – 2.5 mM |
| AMP | amp_c | 0.5 mM – 2.5 mM |
| $NAD^+$ | nad_c | 1 mM |
| NADH | nadh_c | 10 µM – 100 µM |
| $NADP^+$ | nadp_c | 10 µM |
| NADPH | nadph_c | 10 µM – 100 µM |
| $O_2$ | o2_c | 273 µM |
| $CO_2$ | co2_c | 6.3 mM |
| CoA | coa_c | 1 mM – 5 mM |
| orthophosphate | pi_c | 1 mM – 10 mM |
| pyrophosphate | ppi_c | 0.5 mM – 1.5 mM |
| ammonia | nh4_c | 1 mM – 10 mM |
| alphaketoglutarate | akg_c | 0.5 mM – 5 mM |
| glutamate | glu__L_c | 30 mM – 150 mM |

**Table X1**: The allowed concentration ranges for metabolites in the model. All metabolites that do not appear in this table, were constrained by the default ranges of 1 µM to 10 mM.

#### 1.2 Mixed Integer Linear Problem

$$\underset{B,\mathbf{v},\mathbf{z},\mathbf{x}}{\text{maximize}} \quad B - \textstyle\sum_i z_i \tag{1}$$

such that

$$\mathbf{S}\mathbf{v} = \mathbf{0} \tag{2}$$

$$v_{\text{OBJ}} = -1 \tag{3}$$

$$\mathbf{v} - \beta\mathbf{z} \leq 0 \tag{4}$$

$$B \leq -\mathbf{g}^{\circ} - \mathbf{S}^{\top}\mathbf{x} + M(1-\mathbf{z}) \tag{5}$$

$$\mathbf{z} \in \{0,1\}^n \tag{6}$$

$$\mathbf{0} \leq \mathbf{v} \leq \beta \tag{7}$$

$$\ln(\mathbf{C}_{\min}) \leq \mathbf{x} \leq \ln(\mathbf{C}_{\max}) \tag{8}$$

$$0 \leq B \tag{9}$$

where the vector $\mathbf{v}$ contains the relative reaction rates, $\mathbf{z}$ are the Boolean reaction indicators, $\mathbf{x}$ are the log-scaled metabolite concentrations, and $\mathbf{g}^{\circ}$ is a vector of all the reactions' standard Gibbs free energies in units of $RT$, i.e. $\forall i \; g_i^{\circ} \equiv \Delta_r G_i'^{\circ}/RT$. $\beta$ is a parameter that limits the maximal rate for each single reaction in the pathway (relative to the objective reaction, i.e. 3 $CO_2$ → pyruvate), and was set arbitrarily to 10. The rate of the objective reaction ($v_{\text{OBJ}}$) is set to be exactly -1 (Equation 3). This ensures that any pathway solution would exactly balance it, i.e. the overall reaction in the pathway would be 3 $CO_2$ → pyruvate. $M$ is a parameter which has a large value, much higher than any of the values in $\mathbf{g}^{\circ}$. The lower and upper bounds on the concentrations of most metabolites were set to 1 $\mu M$ and 10 mM. Only 14 central metabolites and co-factors were confined to more specific ranges based on physiological data (see Table X1).

Note that as a pre-processing step, we split all reactions to a forward and backward reaction and therefore all (uni-directional) rates must be positive. Equation 4 ensures that a reaction indicator ($z_i$) can be equal to 0, only if the rate $v_i$ is 0. We don't need to care about $z_i$ being equal to 1 even if a reaction is not active, since the optimization goal (which maximizes the sum of all indicators) will prevent that.

Equation 5 ensures that every active reaction has a positive driving force (which is given by $-g_i^{\circ} - \sum_j S_{ij} x_j$). If $z_i = 1$, the driving force must be larger than $B$, which is a positive number that represents a margin. We add $B$ to the optimization function, in order to maximize this margin. This approach is based on the Max-min Driving Force [3] and aims to prioritize pathways that can be operated as far from equilibrium as possible. The method for using Max-min Driving Force optimization for finding feasible pathways in the genome-scale *E. coli* model was introduced by Hädicke et al. [1], and denoted OptMDFpathway.

Our MILP objective function (Equation 1) is the margin ($B$) minus the sum of all indicators (which is equal to the number of reactions in the pathway). Maximizing this function will simultaneously maximize the MDF and minimize the pathway length. Importantly, when combining two optimization functions, the relative weight given to each one is very important. Since the MDF is given in units of $RT$, and the pathway length is an integer, using equal weights is an arbitrary choice. For example, giving a much higher weight to the MDF (by changing the units, or multiplying it by a large pre-factor) would likely change the MILP solution. In this work, we wanted to avoid tuning the relative optimization weights. Instead, we iterate the space of sub-optimal solutions and try to identify pathways that are Pareto-optimal (i.e., no other solution outperforms them in both MDF and length). This procedure is explained in detail in the next section.

##### 1.3 Iterating the space of solutions

In order to cover the space of thermodynamically feasible solutions (i.e. pathways with MDF $> 0$), we iteratively use integer-cuts to eliminate all previous solutions and find the next optimal one [4]. Formally, if $P_0, \ldots, P_m$ are the set of solutions already discovered by our algorithm (where $P_j \subset \{0, \ldots, n\}$) then the added constraints will be:

$$\forall j \quad \sum_{i \in P_j} z_i < |P_j| \tag{10}$$

where $|P_j|$ is the size of the pathway (i.e. the number of reactions). Each one of these constraints eliminates $P_j$ and any pathway which is a superset of $P_j$ from the solution space.

Using IBM's CPLEX solver, we could recover only $\sim$100 solutions per day, on an Intel Core i7-4770S CPU (with 8 cores). However, when running the iterative search for about 3 days, we noticed that even though the solutions were different by at least one reaction, the overlap between them was quite large and running the search exhaustively would take a significant amount of time. Therefore, in order to increase the diversity of the solutions and shortening the run-time, we changed the solution elimination process, so that each time all solutions within a radius of 3 reactions would also be eliminated. To achieve that, we subtract 3 from the right-hand side of each constraint:

$$\forall j \quad \sum_{i \in P_j} z_i < |P_j| - 3\,. \tag{11}$$

After this modification, the diversity of the pathways within the first 50 was much larger, which we measure by counting which carboxylating enzymes were used in each pathway (see Figure X1 and Table X2). Since we optimize both the MDF and number of reactions in each iteration, it is very unlikely that pathways that are Pareto-optimal would be excluded from the results due to the 3-radius rule. Nevertheless, we verified that the set of Pareto-optimal solutions is not affected by the exclusion radius.

We find that only 2 pathways are Pareto-optimal in terms of MDF and pathway length. The first one (pathway 0) is the GED cycle, with 17 reactions and an MDF of 3.3 kJ/mol. This is the shortest possible $CO_2$ fixating pathway in *E. coli*. The only other Pareto-optimal pathway, comprising 20 reactions and an MDF of 5.1 kJ/mol, uses the reverse glycine cleavage system as its carboxylating mechanism.

Graphical depictions of these pathways can be found in section 1.5.

##### 1.4 Implementation

The source code and all input and output files can be found on GitLab under an open-source license (MIT). Specifically, a Jupyter Notebook which was used to create Figure X1 can be found here.

#### References

[1] Oliver Hädicke, Axel von Kamp, Timur Aydogan, and Steffen Klamt. OptMDFpathway: Identification of metabolic pathways with maximal thermodynamic driving force and its application for analyzing the endogenous CO2 fixation potential of Escherichia coli. *PLOS Computational Biology*, 14(9):e1006492, September 2018.

[2] Jonathan M Monk, Colton J Lloyd, Elizabeth Brunk, Nathan Mih, Anand Sastry, Zachary King, Rikiya Takeuchi, Wataru Nomura, Zhen Zhang, Hirotada Mori, Adam M Feist, and Bernhard O Palsson. iML1515, a knowledgebase that computes Escherichia coli traits. *Nature Biotechnology*, 35(10):904–908, October 2017.

[3] Elad Noor, Arren Bar-Even, Avi Flamholz, Ed Reznik, Wolfram Liebermeister, and Ron Milo. Pathway Thermodynamics Highlights Kinetic Obstacles in Central Metabolism. *PLOS Computational Biology*, 10(2):e1003483, February 2014.

[4] Priti Pharkya, Anthony P. Burgard, and Costas D. Maranas. OptStrain: A computational framework for redesign of microbial production systems. *Genome Research*, 14(11):2367–2376, November 2004.

| Pathway | MDF [kJ/mol] | No. reactions | Carboxylating reactions |
|---|---|---|---|
| 0 (GED) | 3.29 | 17.0 | GND |
| 1 | 2.93 | 19.0 | ME1 + GND |
| 2 | 5.13 | 20.0 | GLYCL |
| 3 | 0.17 | 18.0 | GND |
| 4 | 2.19 | 19.0 | GND |
| 5 | 1.02 | 19.0 | GND |
| 6 | 4.27 | 21.0 | GLYCL |
| 7 | 0.15 | 20.0 | ME1 + GND |
| 8 | 0.84 | 21.0 | GND |
| 9 | 5.13 | 23.0 | GLYCL |
| 10 | 0.15 | 21.0 | GND |
| 11 | 1.11 | 22.0 | ME1 + GND |
| 12 | 0.83 | 22.0 | GND |
| 13 | 3.12 | 23.0 | ME1 + GLYCL |
| 14 | 3.12 | 23.0 | GLYCL + PPC |
| 15 | 0.55 | 22.0 | GND |
| 16 | 2.87 | 23.0 | GLYCL |
| 17 | 5.13 | 24.0 | GLYCL |
| 18 | 0.17 | 22.0 | ME1 + GND |
| 19 | 2.19 | 23.0 | ME1 + GND |
| 20 | 2.19 | 23.0 | ME1 + GND |
| 21 | 0.39 | 23.0 | GND |
| 22 | 5.13 | 25.0 | GLYCL |
| 23 | 4.94 | 25.0 | GLYCL |
| 24 | 1.87 | 24.0 | GLYCL |
| 25 | 4.27 | 25.0 | GLYCL |
| 26 | 1.61 | 24.0 | ME1 + GND |
| 27 | 1.53 | 24.0 | GND |
| 28 | 1.43 | 24.0 | GND |
| 29 | 1.32 | 24.0 | ME1 |
| 30 | 0.9 | 24.0 | GND |
| 31 | 3.13 | 25.0 | PPCK + GLYCL |
| 32 | 0.55 | 24.0 | GND |
| 33 | 0.44 | 24.0 | GND |
| 34 | 0.44 | 24.0 | GND |
| 35 | 0.33 | 24.0 | GND |
| 36 | 2.64 | 25.0 | PPCK + GLYCL |
| 37 | 2.64 | 25.0 | ME1 + GLYCL |
| 38 | 0.07 | 24.0 | GLYCL + PPC |
| 39 | 0.03 | 24.0 | GND |
| 40 | 1.53 | 25.0 | ME1 + GND |
| 41 | 3.92 | 26.0 | ME1 + GLYCL |
| 42 | 1.32 | 25.0 | ME1 |
| 43 | 1.32 | 25.0 | ME1 |
| 44 | 0.95 | 25.0 | GND |
| 45 | 0.89 | 25.0 | GND |
| 46 | 0.83 | 25.0 | ME1 + GND |
| 47 | 0.83 | 25.0 | GND |
| 48 | 3.17 | 26.0 | GLYCL |
| 49 | 3.12 | 26.0 | PPCK |

**Table X2**: The Max-min Driving Force and number of reactions of all 50 top pathways. A graph depiction of these pathways can be found in section 1.5.

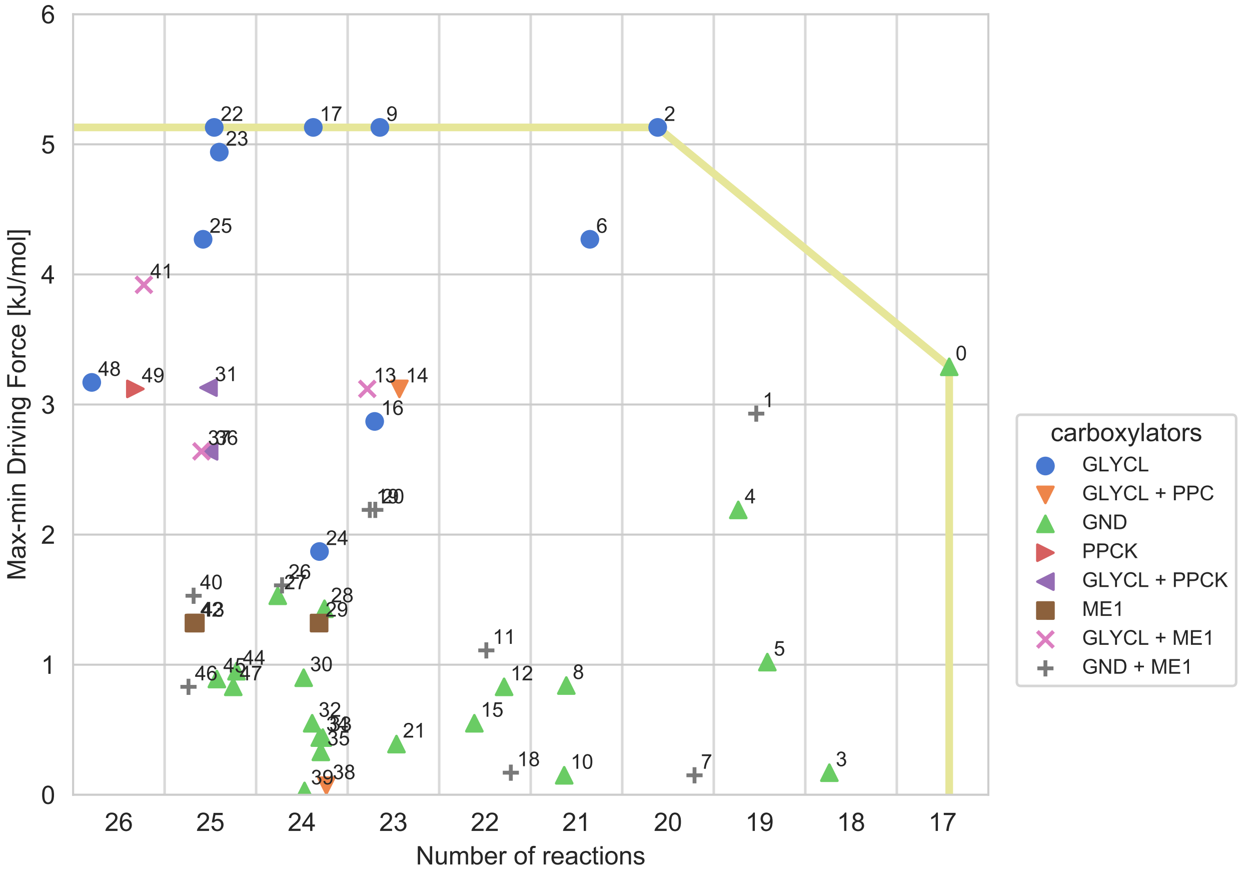


**Figure X1**: The MDF versus the number of reactions for all the first 50 solutions. The only two pathways on the Pareto front are 0 and 2. The radius of exclusion around each solution is 3 reactions. The full list of pathways can be found in in section 1.5 or on the GitLab repository. The carboxylating enzymes we found in our pathways are: PPCK (phosphoenolpyruvate carboxykinase), GND (phosphogluconate dehydrogenase), PPC (phosphoenolpyruvate carboxylase), GLYCL (glycine cleavage system), ME1 (malic enyzme NAD-dependent). The Pareto front is marked by a yellow line.

#### 1.5 Graphical depictions of the top 50 $CO_2$ fixation pathways

Pathway 0 (the GED cycle)

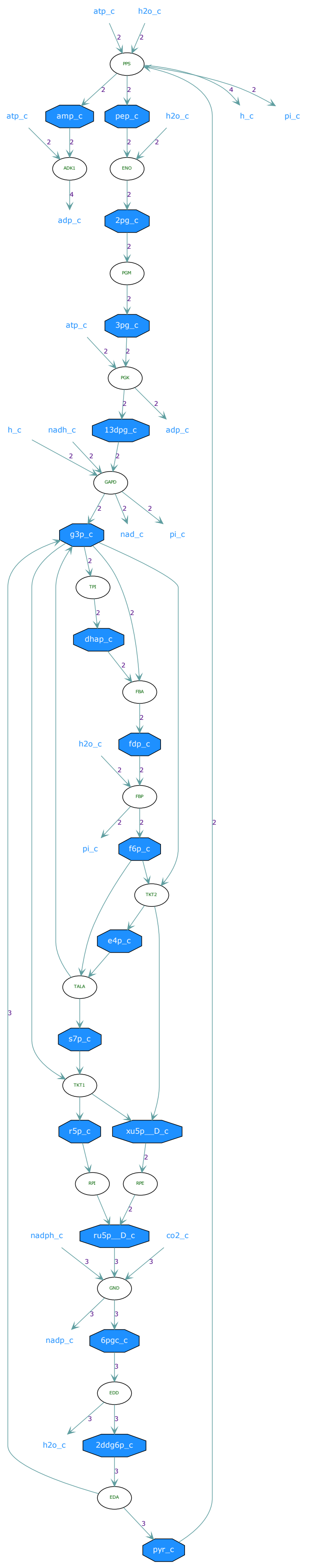

### Pathway 1

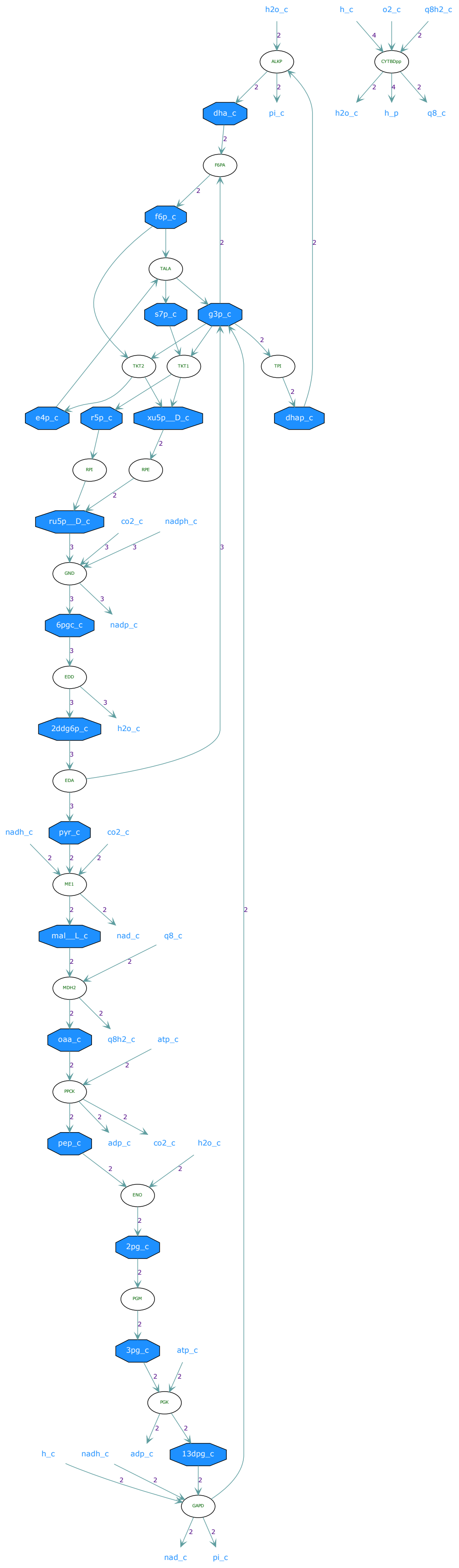
h2o_c
h_c
o2_c
q8h2_c
ALKP
CYTBDpp
dha_c
pi_c
h2o_c
h_p
q8_c
F6PA
f6p_c
TALA
s7p_c
g3p_c
TKT2
TKT1
TPI
e4p_c
r5p_c
xu5p__D_c
dhap_c
RPI
RPE
ru5p__D_c
co2_c
nadph_c
GND
6pgc_c
nadp_c
EDD
2ddg6p_c
h2o_c
EDA
nadh_c
pyr_c
co2_c
ME1
mal__L_c
nad_c
q8_c
MDH2
oaa_c
q8h2_c
atp_c
PPCK
pep_c
adp_c
co2_c
h2o_c
ENO
2pg_c
PGM
3pg_c
atp_c
PGK
h_c
nadh_c
adp_c
13dpg_c
GAPD
nad_c
pi_c

#### Pathway 2

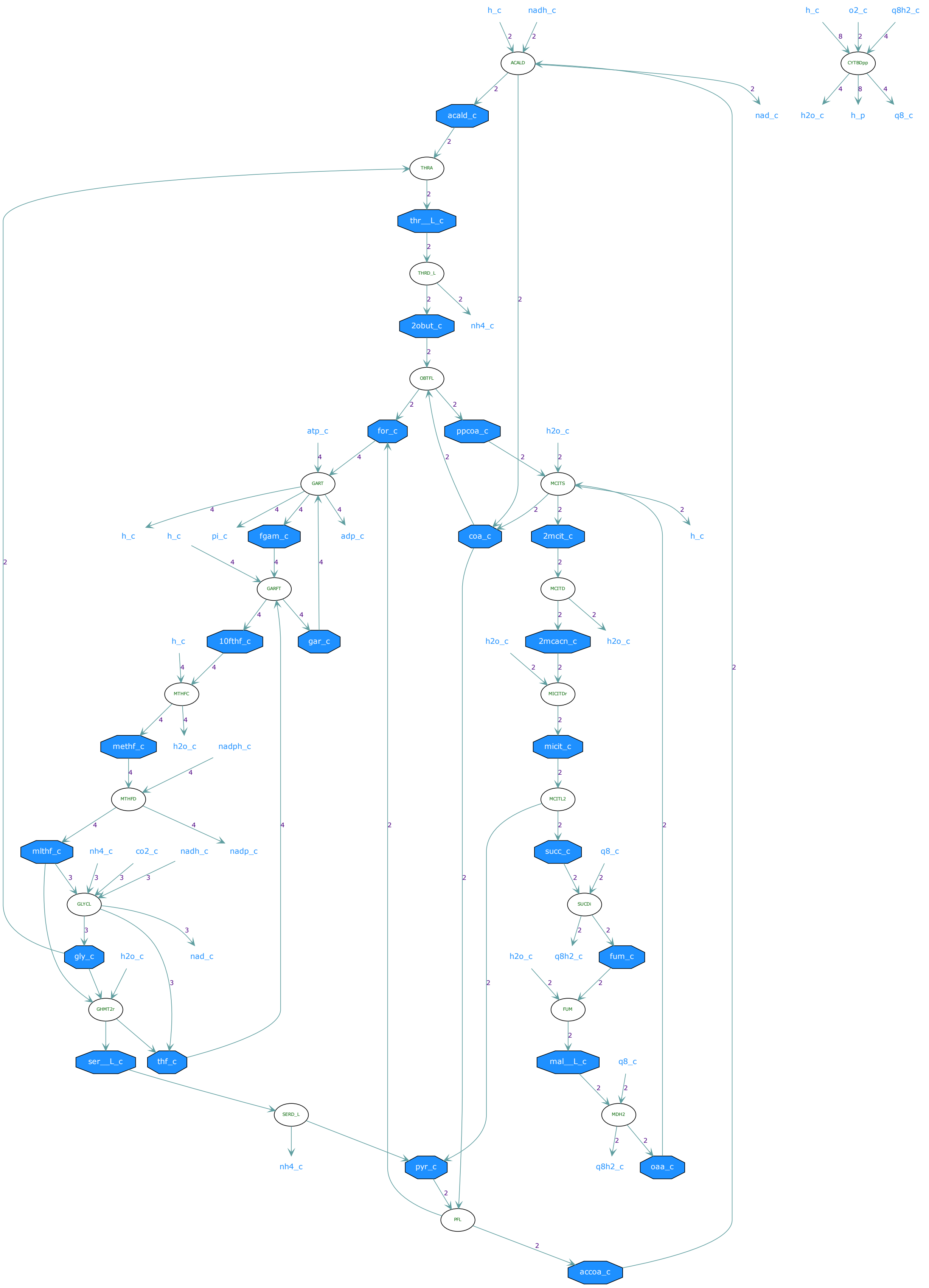
h_c
nadh_c
ACALD
acald_c
nad_c
h_c
o2_c
q8h2_c
CYTBDpp
h2o_c
h_p
q8_c
THRA
thr__L_c
THRD_L
2obut_c
nh4_c
OBTFL
for_c
ppcoa_c
atp_c
h2o_c
GART
MCITS
h_c
h_c
pi_c
fgam_c
adp_c
coa_c
2mcit_c
h_c
GARFT
MCITD
10fthf_c
gar_c
h_c
h2o_c
2mcacn_c
h2o_c
MTHFC
MICITDr
methf_c
h2o_c
nadph_c
micit_c
MTHFD
MCITL2
mlthf_c
nh4_c
co2_c
nadh_c
nadp_c
succ_c
q8_c
GLYCL
SUCDi
gly_c
h2o_c
nad_c
h2o_c
q8h2_c
fum_c
GHMT2r
FUM
ser__L_c
thf_c
mal__L_c
q8_c
SERD_L
MDH2
nh4_c
pyr_c
q8h2_c
oaa_c
PFL
accoa_c

Pathway 3

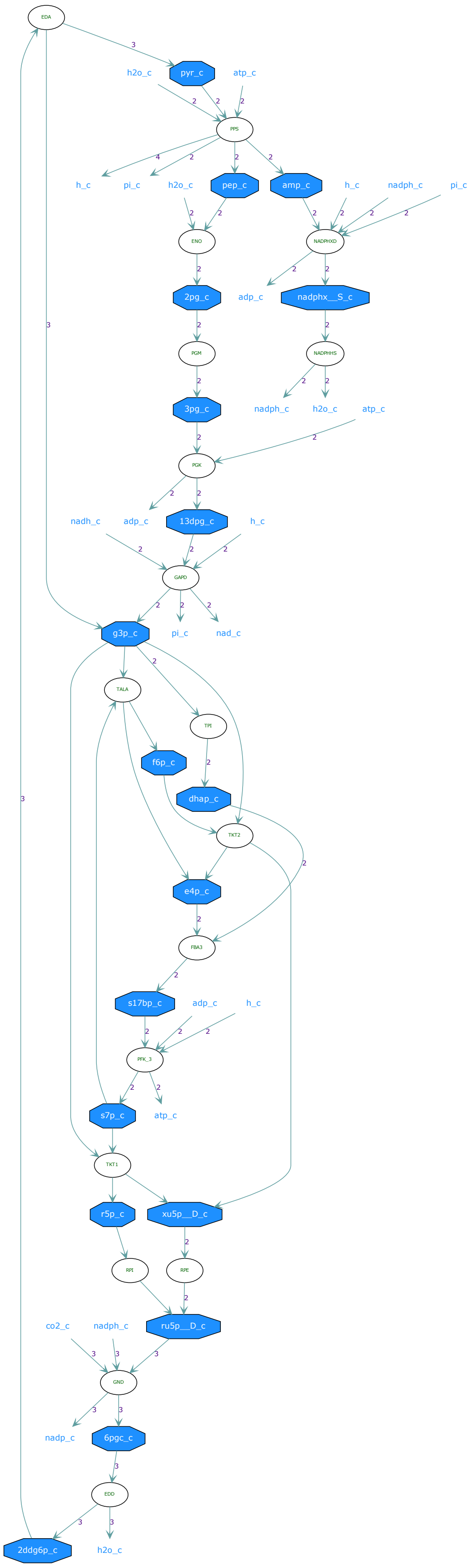
EDA
pyr_c
h2o_c
atp_c
PPS
h_c
pi_c
h2o_c
pep_c
amp_c
h_c
nadph_c
pi_c
ENO
NADPHXD
2pg_c
adp_c
nadphx__S_c
PGM
NADPHHS
3pg_c
nadph_c
h2o_c
atp_c
PGK
nadh_c
adp_c
13dpg_c
h_c
GAPD
g3p_c
pi_c
nad_c
TALA
TPI
f6p_c
dhap_c
TKT2
e4p_c
FBA3
s17bp_c
adp_c
h_c
PFK_3
s7p_c
atp_c
TKT1
r5p_c
xu5p__D_c
RPI
RPE
co2_c
nadph_c
ru5p__D_c
GND
nadp_c
6pgc_c
EDD
2ddg6p_c
h2o_c

Pathway 4

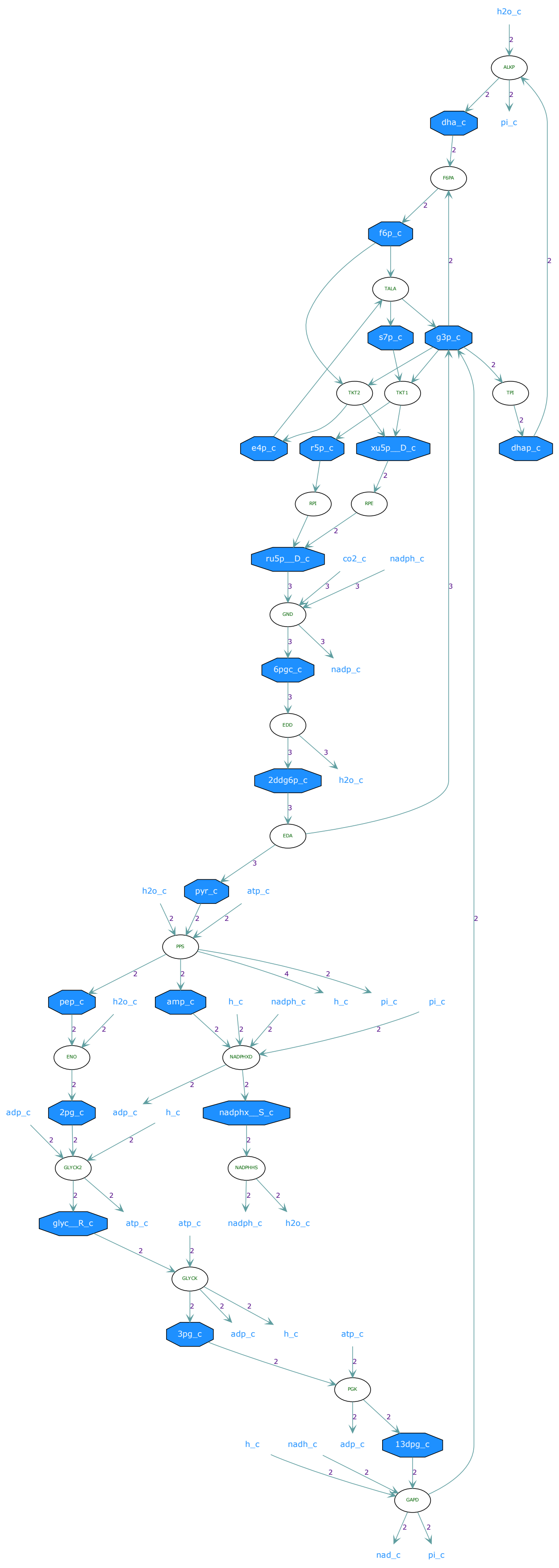
h2o_c
ALKP
dha_c
pi_c
F6PA
f6p_c
TALA
s7p_c
g3p_c
TKT2
TKT1
TPI
e4p_c
r5p_c
xu5p__D_c
dhap_c
RPI
RPE
ru5p__D_c
co2_c
nadph_c
GND
6pgc_c
nadp_c
EDD
2ddg6p_c
h2o_c
EDA
h2o_c
pyr_c
atp_c
PPS
pep_c
h2o_c
amp_c
h_c
nadph_c
h_c
pi_c
pi_c
ENO
NADPHXD
adp_c
2pg_c
adp_c
h_c
nadphx__S_c
GLYCK2
NADPHHS
glyc__R_c
atp_c
atp_c
nadph_c
h2o_c
GLYCK
3pg_c
adp_c
h_c
atp_c
PGK
h_c
nadh_c
adp_c
13dpg_c
GAPD
nad_c
pi_c

### Pathway 5

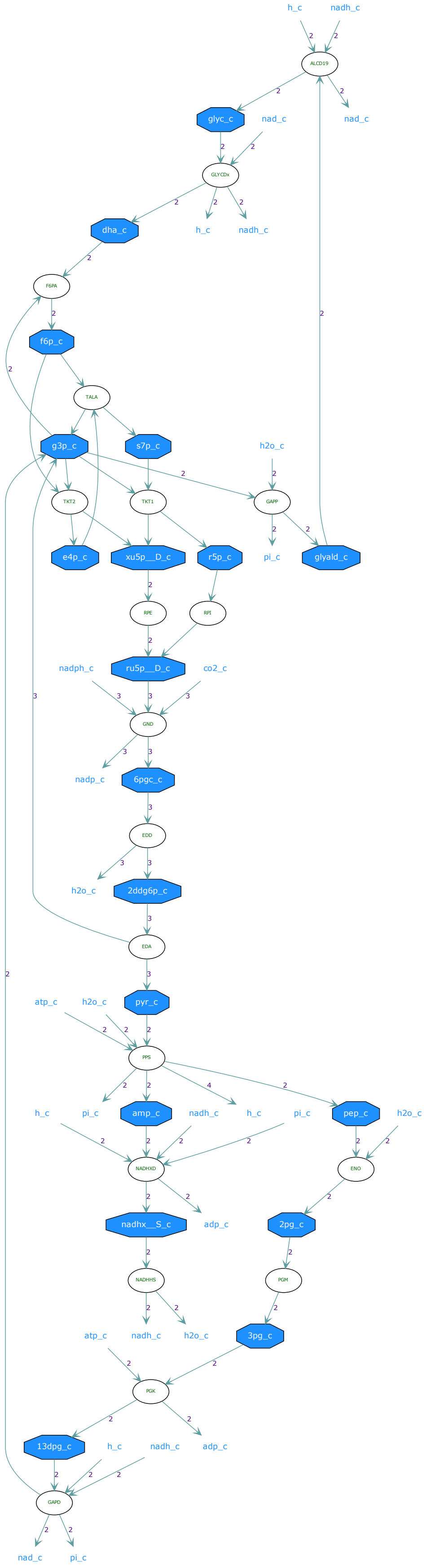
h_c
nadh_c
ALCD19
glyc_c
nad_c
nad_c
GLYCDx
dha_c
h_c
nadh_c
F6PA
f6p_c
TALA
g3p_c
s7p_c
h2o_c
TKT2
TKT1
GAPP
e4p_c
xu5p__D_c
r5p_c
pi_c
glyald_c
RPE
RPI
nadph_c
ru5p__D_c
co2_c
GND
nadp_c
6pgc_c
EDD
h2o_c
2ddg6p_c
EDA
atp_c
h2o_c
pyr_c
PPS
h_c
pi_c
amp_c
nadh_c
h_c
pi_c
pep_c
h2o_c
NADHXD
ENO
nadhx__S_c
adp_c
2pg_c
NADHHS
PGM
atp_c
nadh_c
h2o_c
3pg_c
PGK
13dpg_c
h_c
nadh_c
adp_c
GAPD
nad_c
pi_c

### Pathway 6

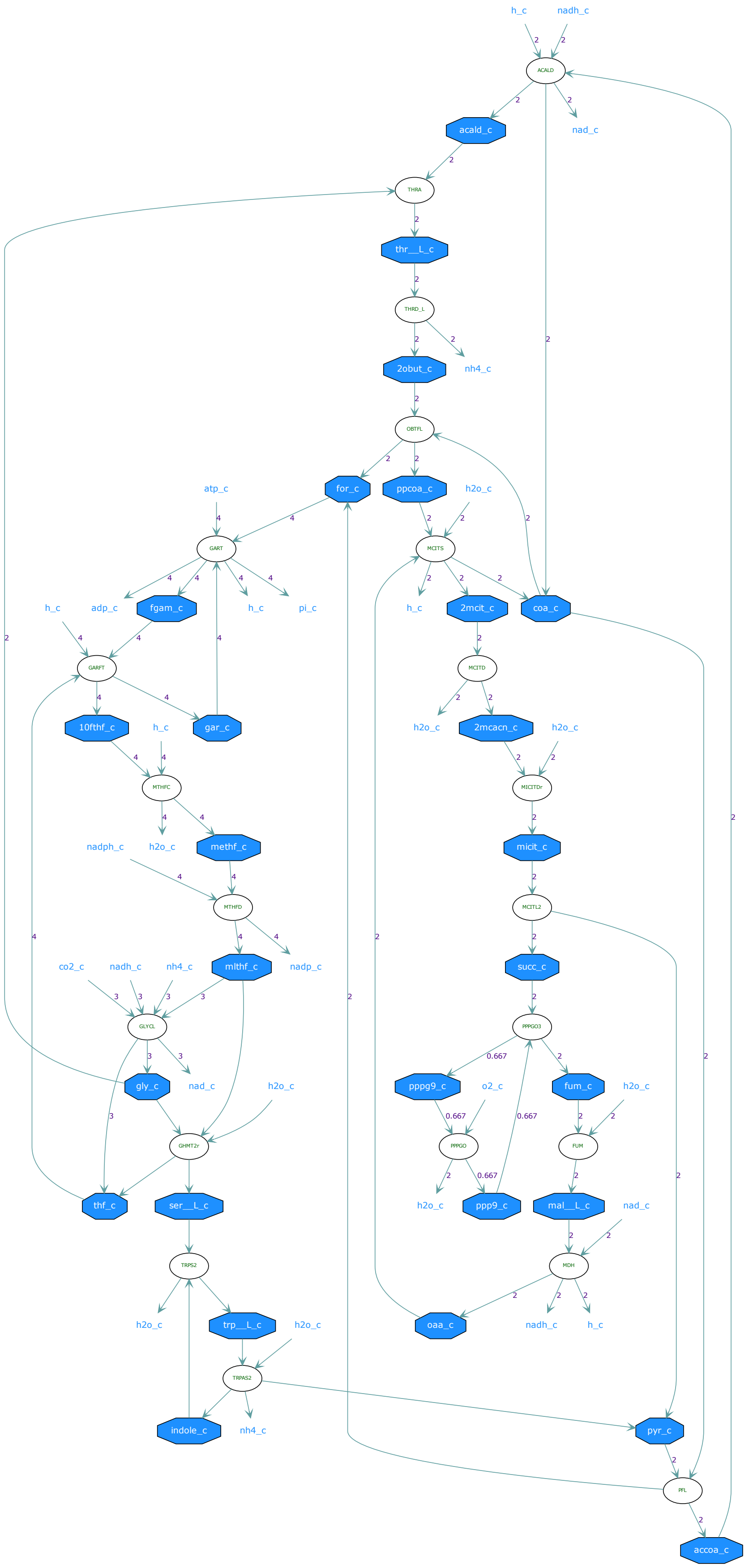

h_c
nadh_c
ACALD
acald_c
nad_c
THRA
thr__L_c
THRD_L
2obut_c
nh4_c
OBTFL
for_c
ppcoa_c
h2o_c
atp_c
GART
MCITS
adp_c
fgam_c
pi_c
2mcit_c
coa_c
GARFT
MCITD
10fthf_c
gar_c
2mcacn_c
MTHFC
MICITDr
nadph_c
methf_c
micit_c
MTHFD
MCITL2
co2_c
mlthf_c
nadp_c
succ_c
GLYCL
PPPGO3
gly_c
pppg9_c
o2_c
fum_c
GHMT2r
PPPGO
FUM
thf_c
ser__L_c
ppp9_c
mal__L_c
TRPS2
MDH
trp__L_c
oaa_c
TRPAS2
indole_c
pyr_c
PFL
accoa_c

### Pathway 7

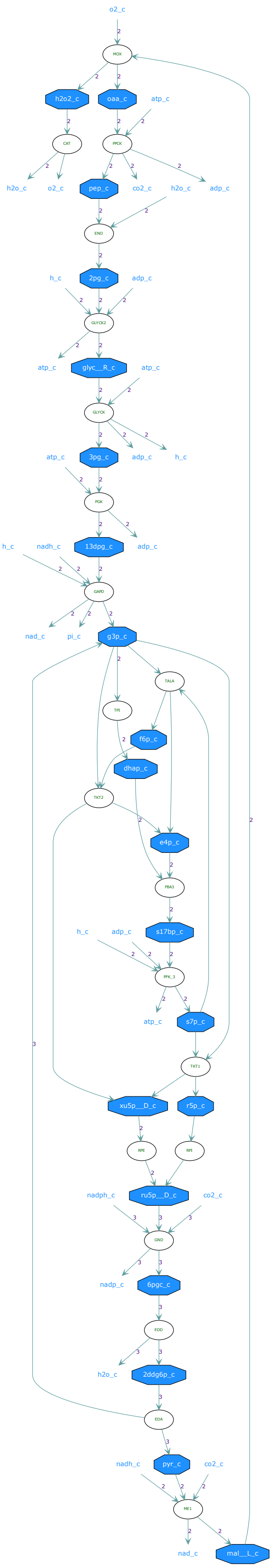
o2_c
MOX
h2o2_c
oaa_c
atp_c
CAT
PPCK
h2o_c
o2_c
pep_c
co2_c
h2o_c
adp_c
ENO
2pg_c
h_c
adp_c
GLYCK2
atp_c
glyc__R_c
atp_c
GLYCK
3pg_c
atp_c
adp_c
h_c
PGK
13dpg_c
h_c
nadh_c
adp_c
GAPD
nad_c
pi_c
g3p_c
TALA
TPI
f6p_c
dhap_c
TKT2
e4p_c
FBA3
s17bp_c
h_c
adp_c
PFK_3
atp_c
s7p_c
TKT1
xu5p__D_c
r5p_c
RPE
RPI
ru5p__D_c
nadph_c
co2_c
GND
nadp_c
6pgc_c
EDD
h2o_c
2ddg6p_c
EDA
pyr_c
nadh_c
co2_c
ME1
nad_c
mal__L_c

### Pathway 8

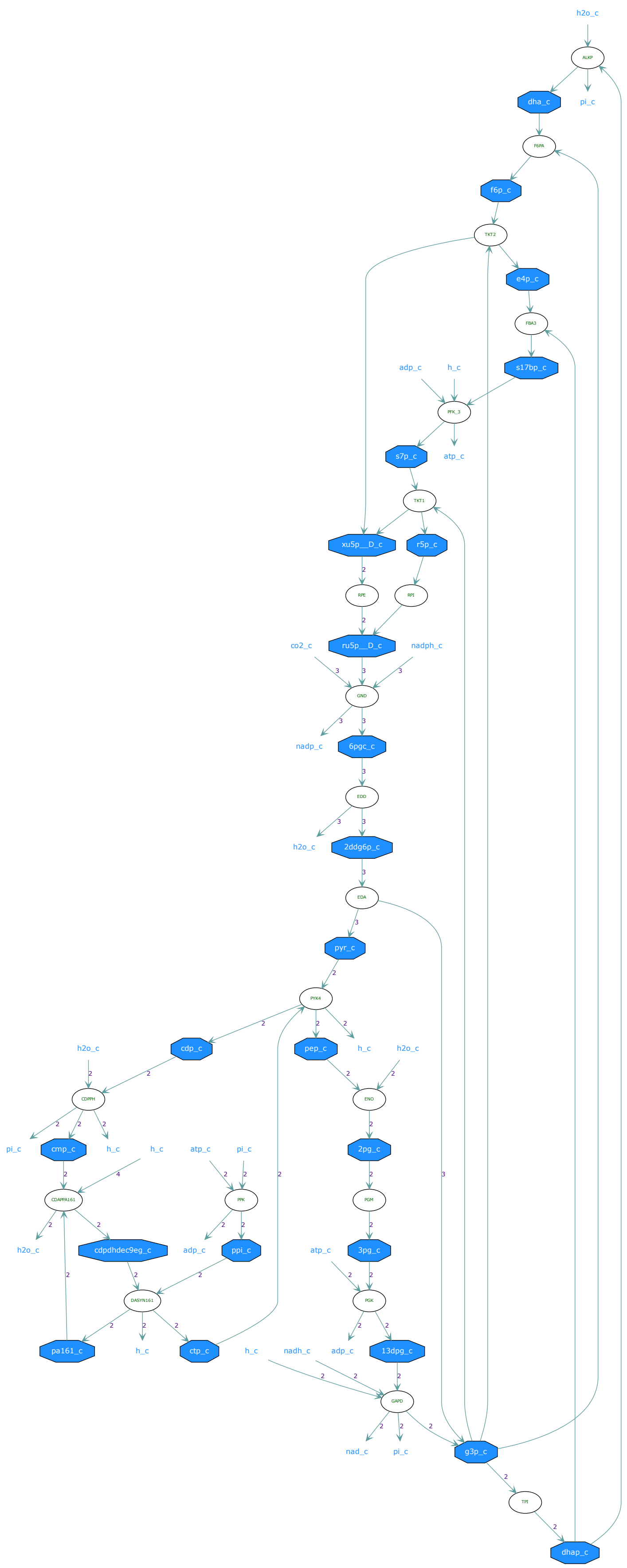
h2o_c
ALKP
dha_c
pi_c
F6PA
f6p_c
TKT2
e4p_c
FBA3
adp_c
h_c
s17bp_c
PFK_3
s7p_c
atp_c
TKT1
xu5p__D_c
r5p_c
RPE
RPI
co2_c
ru5p__D_c
nadph_c
GND
nadp_c
6pgc_c
EDD
h2o_c
2ddg6p_c
EDA
pyr_c
PYK4
h2o_c
cdp_c
pep_c
h_c
h2o_c
CDPPH
ENO
pi_c
cmp_c
h_c
h_c
atp_c
pi_c
2pg_c
CDAPPA161
PPK
PGM
h2o_c
cdpdhdec9eg_c
adp_c
ppi_c
atp_c
3pg_c
DASYN161
PGK
pa161_c
h_c
ctp_c
h_c
nadh_c
adp_c
13dpg_c
GAPD
nad_c
pi_c
g3p_c
TPI
dhap_c

### Pathway 9

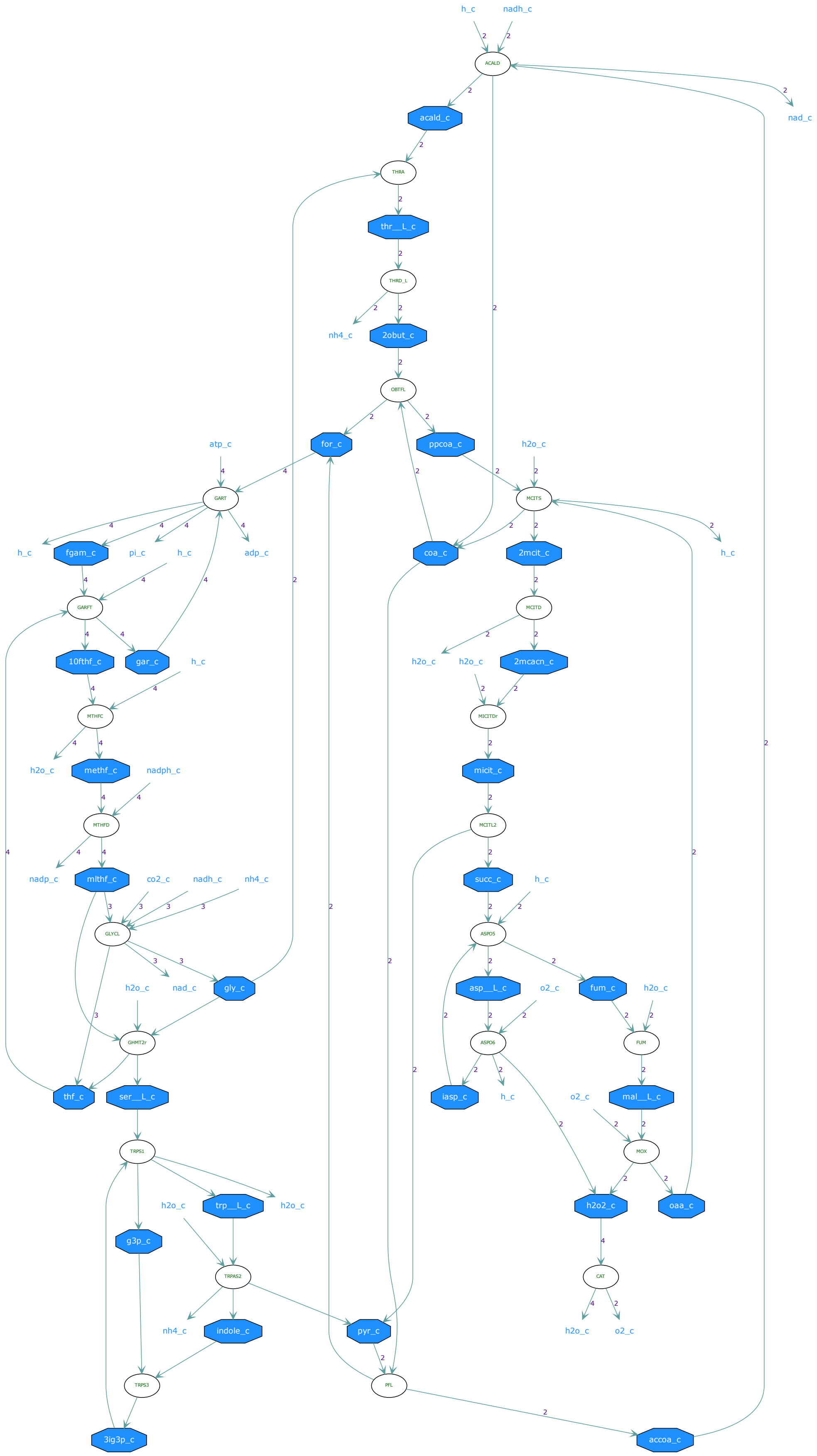

h_c
nadh_c
ACALD
nad_c
acald_c
THRA
thr__L_c
THRD_L
nh4_c
2obut_c
OBTFL
for_c
ppcoa_c
atp_c
h2o_c
GART
MCITS
h_c
fgam_c
pi_c
h_c
adp_c
coa_c
2mcit_c
h_c
GARFT
MCITD
10fthf_c
gar_c
h_c
h2o_c
h2o_c
2mcacn_c
MTHFC
MICITDr
h2o_c
methf_c
nadph_c
micit_c
MTHFD
MCITL2
nadp_c
mlthf_c
co2_c
nadh_c
nh4_c
succ_c
h_c
GLYCL
ASPO5
h2o_c
nad_c
gly_c
asp__L_c
o2_c
fum_c
h2o_c
GHMT2r
ASPO6
FUM
thf_c
ser__L_c
iasp_c
h_c
o2_c
mal__L_c
TRPS1
MOX
h2o_c
trp__L_c
h2o_c
h2o2_c
oaa_c
g3p_c
TRPAS2
CAT
nh4_c
indole_c
pyr_c
h2o_c
o2_c
TRPS3
PFL
3ig3p_c
accoa_c

Pathway 10

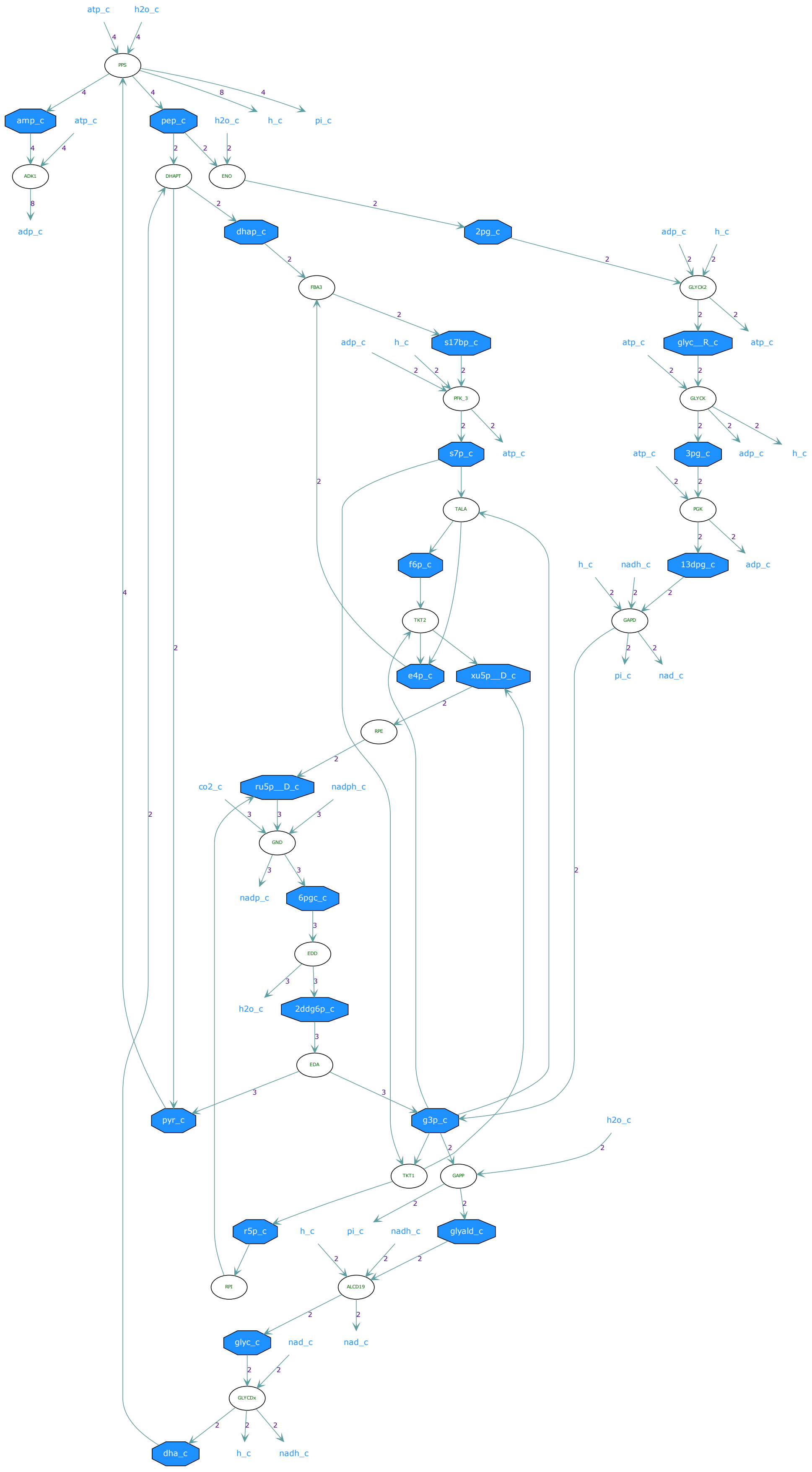
atp_c
h2o_c
PPS
amp_c
atp_c
pep_c
h2o_c
h_c
pi_c
ADK1
DHAPT
ENO
adp_c
dhap_c
2pg_c
adp_c
h_c
FBA3
GLYCK2
adp_c
h_c
s17bp_c
atp_c
glyc__R_c
atp_c
PFK_3
GLYCK
s7p_c
atp_c
atp_c
3pg_c
adp_c
h_c
TALA
PGK
f6p_c
h_c
nadh_c
13dpg_c
adp_c
TKT2
GAPD
e4p_c
xu5p__D_c
pi_c
nad_c
RPE
co2_c
ru5p__D_c
nadph_c
GND
nadp_c
6pgc_c
EDD
h2o_c
2ddg6p_c
EDA
pyr_c
g3p_c
h2o_c
TKT1
GAPP
r5p_c
h_c
pi_c
nadh_c
glyald_c
RPI
ALCD19
glyc_c
nad_c
nad_c
GLYCDx
dha_c
h_c
nadh_c

### Pathway 11

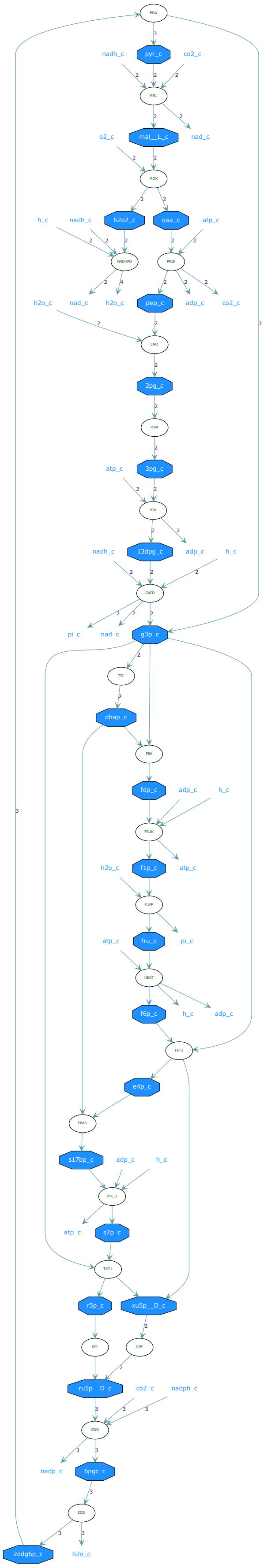

#### Pathway 12

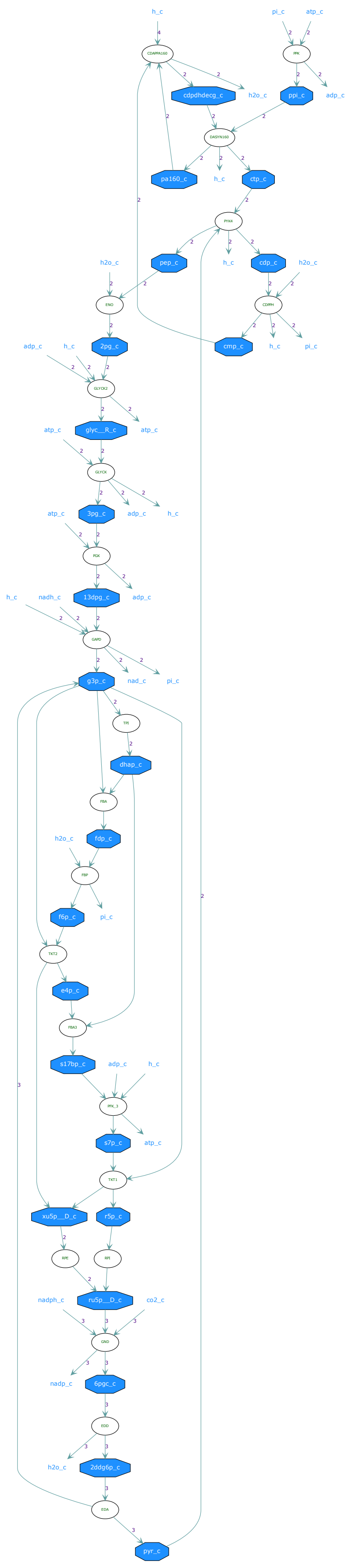
h_c
pi_c
atp_c
cdpdhdecg_c
h2o_c
ppi_c
adp_c
pa160_c
ctp_c
pep_c
cdp_c
2pg_c
cmp_c
glyc__R_c
3pg_c
13dpg_c
nadh_c
g3p_c
nad_c
dhap_c
fdp_c
f6p_c
e4p_c
s17bp_c
s7p_c
xu5p__D_c
r5p_c
ru5p__D_c
nadph_c
co2_c
nadp_c
6pgc_c
2ddg6p_c
pyr_c

### Pathway 13

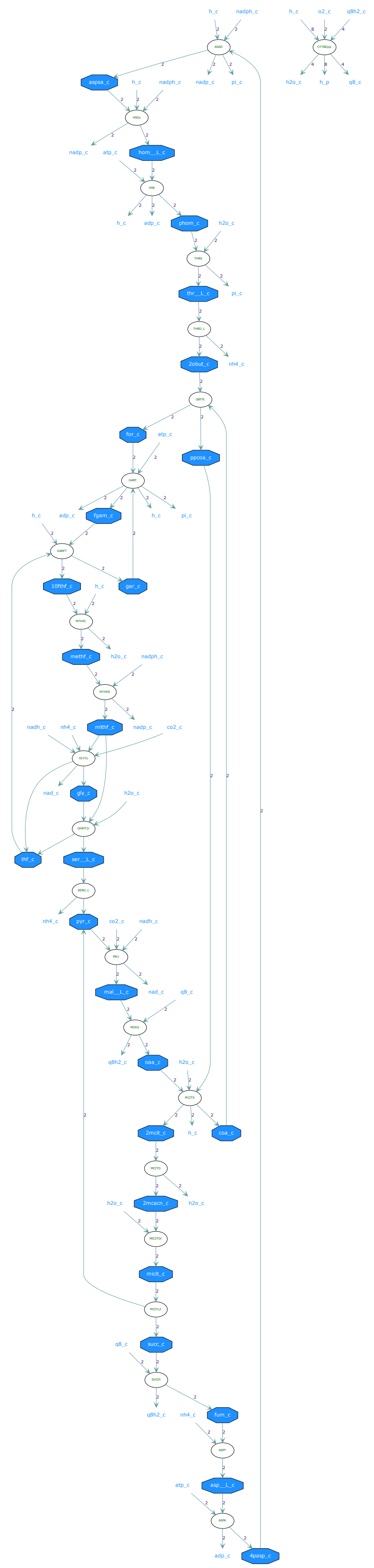
h_c
nadph_c
h_c
o2_c
q8h2_c
aspsa_c
nadp_c
pi_c
h2o_c
h_p
q8_c
nadp_c
atp_c
hom__L_c
h_c
adp_c
phom_c
h2o_c
thr__L_c
pi_c
2obut_c
nh4_c
for_c
atp_c
ppcoa_c
adp_c
fgam_c
pi_c
10fthf_c
gar_c
methf_c
h2o_c
nadph_c
nadh_c
nh4_c
mlthf_c
nadp_c
co2_c
nad_c
gly_c
thf_c
ser__L_c
pyr_c
co2_c
nadh_c
mal__L_c
nad_c
q8h2_c
oaa_c
2mcit_c
coa_c
2mcacn_c
micit_c
succ_c
fum_c
asp__L_c
adp_c
4pasp_c

### Pathway 14

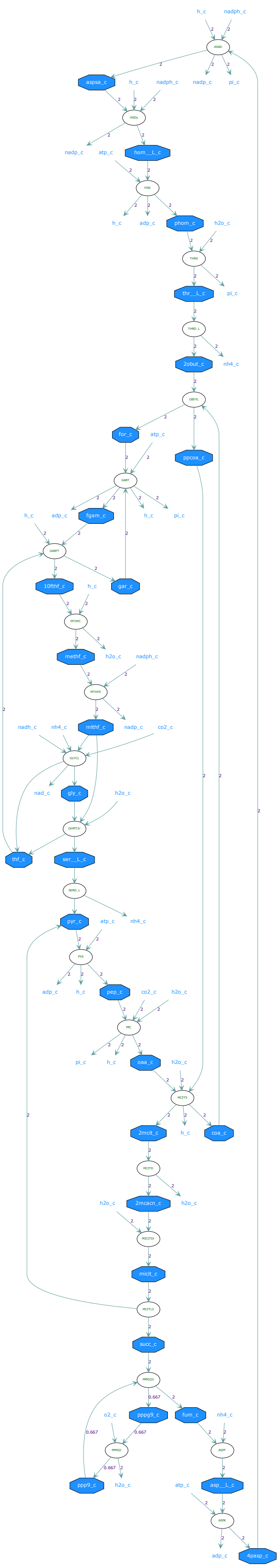
h_c
nadph_c
ASAD
aspsa_c
nadp_c
pi_c
HSDy
hom__L_c
atp_c
HSK
adp_c
phom_c
h2o_c
THRS
thr__L_c
THRD_L
2obut_c
nh4_c
OBTFL
for_c
ppcoa_c
GART
fgam_c
GARFT
10fthf_c
gar_c
MTHFC
methf_c
MTHFD
mlthf_c
nadh_c
co2_c
GLYCL
nad_c
gly_c
GHMT2r
thf_c
ser__L_c
SERD_L
pyr_c
PYK
pep_c
PPC
oaa_c
MCITS
2mcit_c
coa_c
MCITD
2mcacn_c
MICITDr
micit_c
MCITL2
succ_c
PPPGO3
0.667
pppg9_c
fum_c
o2_c
PPPGO
ASPT
ppp9_c
asp__L_c
ASPK
4pasp_c

### Pathway 15

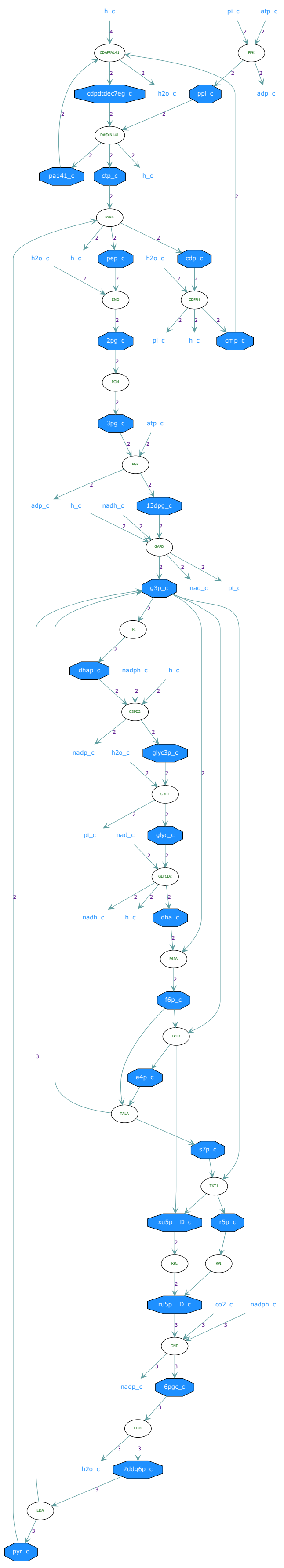
h_c
pi_c
atp_c
cdpdtdec7eg_c
h2o_c
ppi_c
adp_c
pa141_c
ctp_c
pep_c
cdp_c
2pg_c
cmp_c
3pg_c
13dpg_c
nadh_c
g3p_c
nad_c
dhap_c
nadph_c
glyc3p_c
nadp_c
glyc_c
dha_c
f6p_c
e4p_c
s7p_c
xu5p__D_c
r5p_c
ru5p__D_c
co2_c
6pgc_c
2ddg6p_c
pyr_c

### Pathway 16

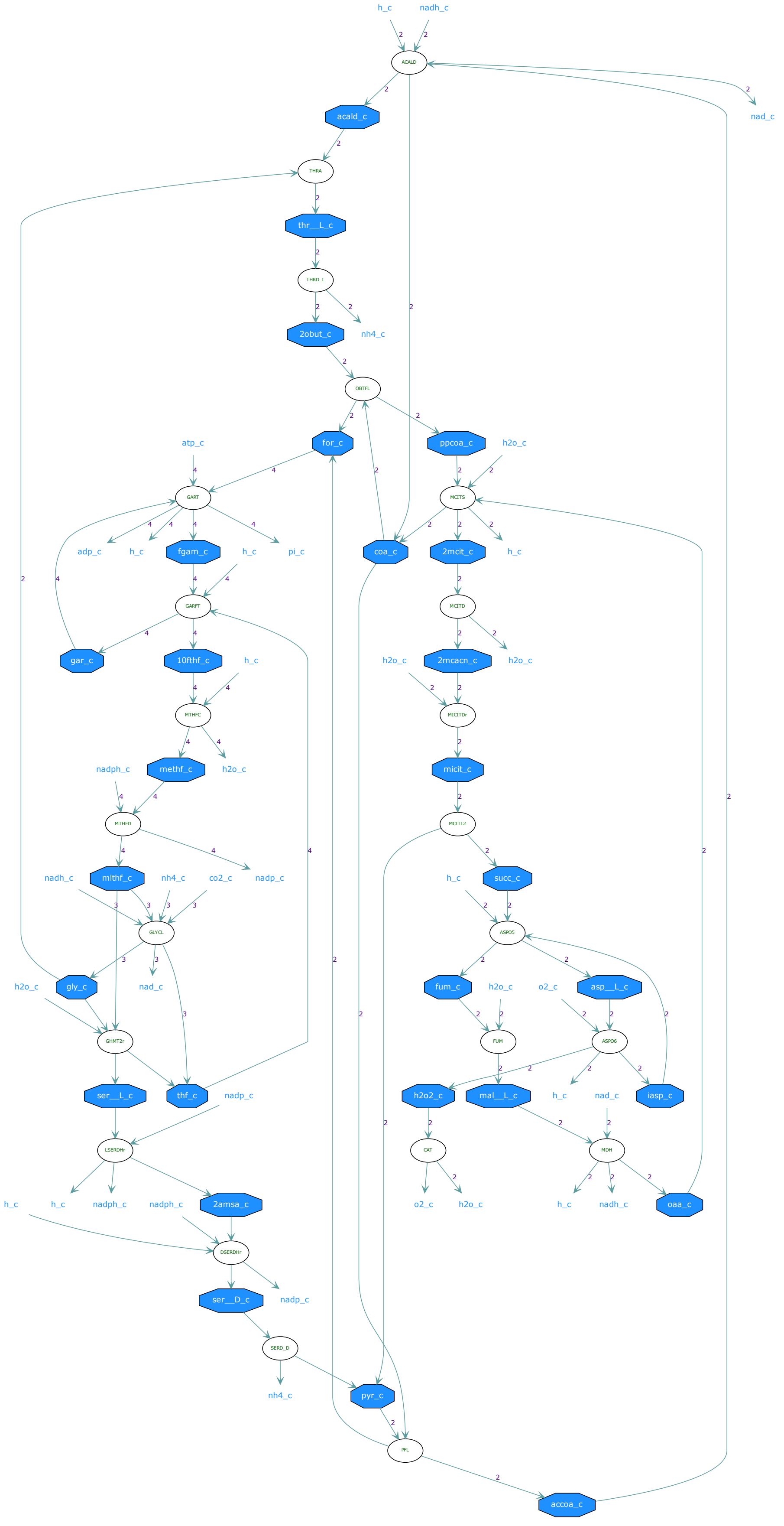

h_c
nadh_c
ACALD
nad_c
acald_c
THRA
thr__L_c
THRD_L
2obut_c
nh4_c
OBTFL
atp_c
for_c
ppcoa_c
h2o_c
GART
MCITS
adp_c
h_c
fgam_c
h_c
pi_c
coa_c
2mcit_c
h_c
GARFT
MCITD
gar_c
10fthf_c
h_c
h2o_c
2mcacn_c
h2o_c
MTHFC
MICITDr
nadph_c
methf_c
h2o_c
micit_c
MTHFD
MCITL2
nadh_c
mlthf_c
nh4_c
co2_c
nadp_c
h_c
succ_c
GLYCL
ASPO5
h2o_c
gly_c
nad_c
fum_c
h2o_c
o2_c
asp__L_c
GHMT2r
FUM
ASPO6
ser__L_c
thf_c
nadp_c
h2o2_c
mal__L_c
h_c
nad_c
iasp_c
LSERDHr
CAT
MDH
h_c
h_c
nadph_c
nadph_c
2amsa_c
o2_c
h2o_c
h_c
nadh_c
oaa_c
DSERDHr
ser__D_c
nadp_c
SERD_D
nh4_c
pyr_c
PFL
accoa_c

### Pathway 17

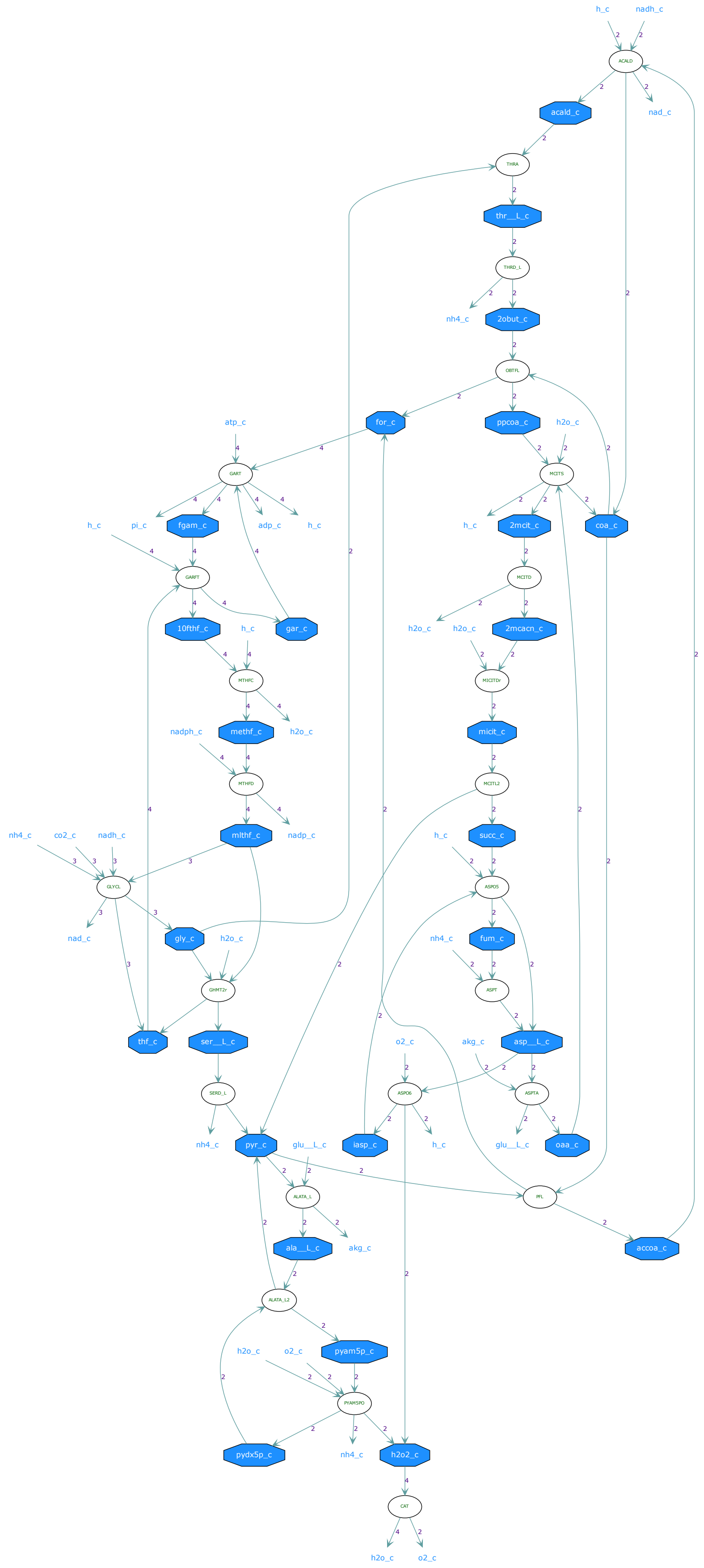

h_c
nadh_c
ACALD
acald_c
nad_c
THRA
thr__L_c
THRD_L
nh4_c
2obut_c
OBTFL
for_c
ppcoa_c
h2o_c
atp_c
GART
MCITS
h_c
pi_c
fgam_c
adp_c
h_c
h_c
2mcit_c
coa_c
GARFT
MCITD
10fthf_c
h_c
gar_c
h2o_c
h2o_c
2mcacn_c
MTHFC
MICITDr
nadph_c
methf_c
h2o_c
micit_c
MTHFD
MCITL2
nh4_c
co2_c
nadh_c
mlthf_c
nadp_c
h_c
succ_c
GLYCL
ASPO5
nad_c
gly_c
h2o_c
nh4_c
fum_c
GHMT2r
ASPT
thf_c
ser__L_c
o2_c
akg_c
asp__L_c
SERD_L
ASPO6
ASPTA
nh4_c
pyr_c
glu__L_c
iasp_c
h_c
glu__L_c
oaa_c
ALATA_L
PFL
ala__L_c
akg_c
accoa_c
ALATA_L2
h2o_c
o2_c
pyam5p_c
PYAM5PO
pydx5p_c
nh4_c
h2o2_c
CAT
h2o_c
o2_c

### Pathway 18

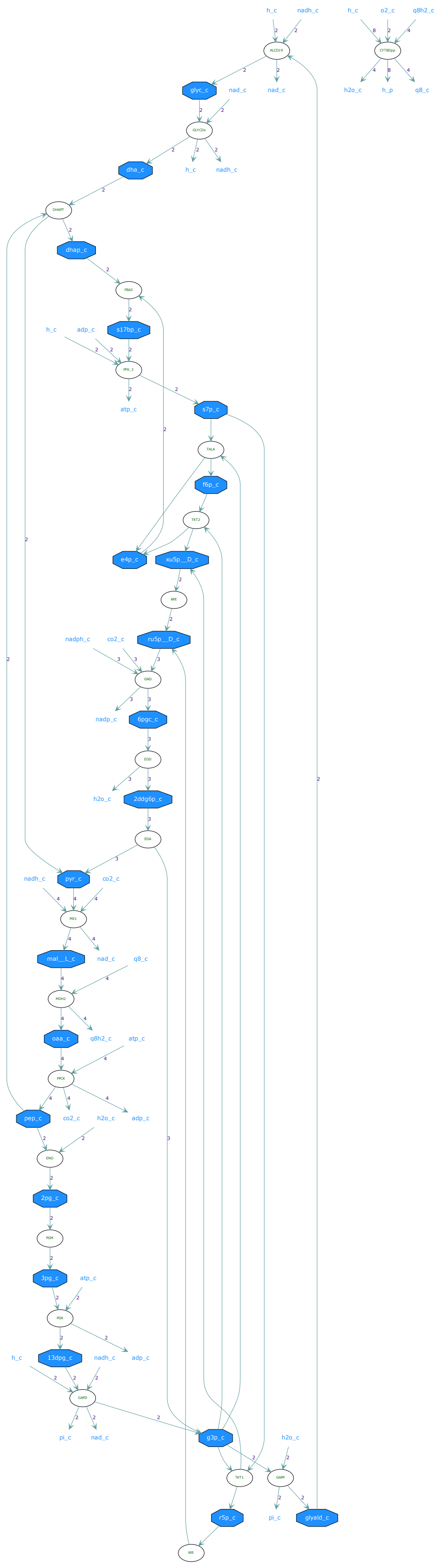
h_c
nadh_c
o2_c
q8h2_c
glyc_c
nad_c
h2o_c
h_p
q8_c
dha_c
dhap_c
s17bp_c
adp_c
atp_c
s7p_c
f6p_c
e4p_c
xu5p__D_c
ru5p__D_c
nadph_c
co2_c
nadp_c
6pgc_c
2ddg6p_c
pyr_c
mal__L_c
oaa_c
pep_c
2pg_c
3pg_c
13dpg_c
pi_c
g3p_c
r5p_c
glyald_c

### Pathway 19

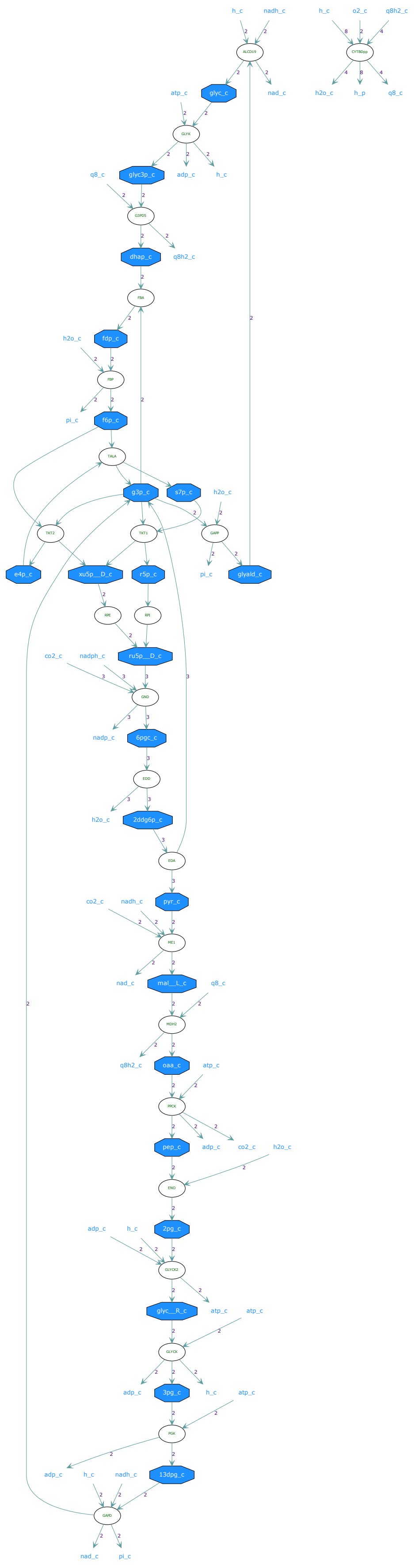

### Pathway 20

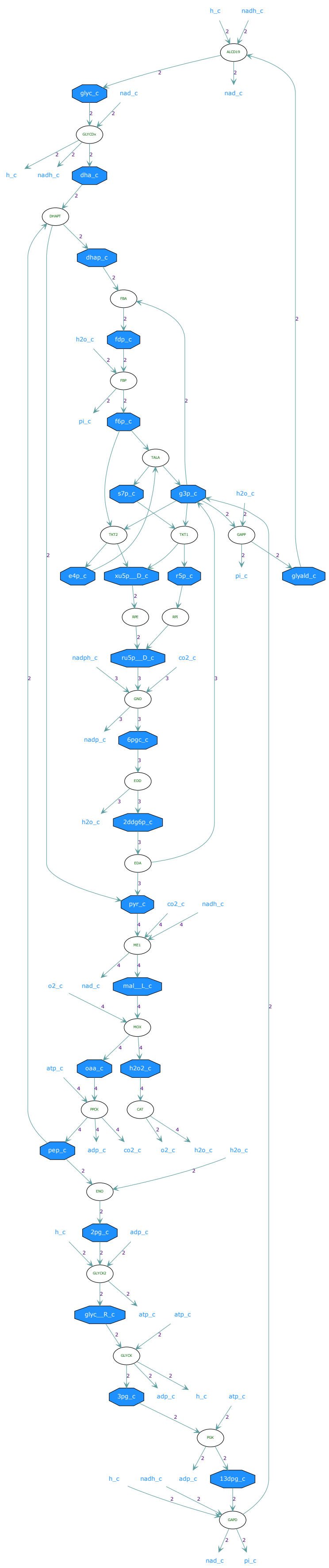

### Pathway 21

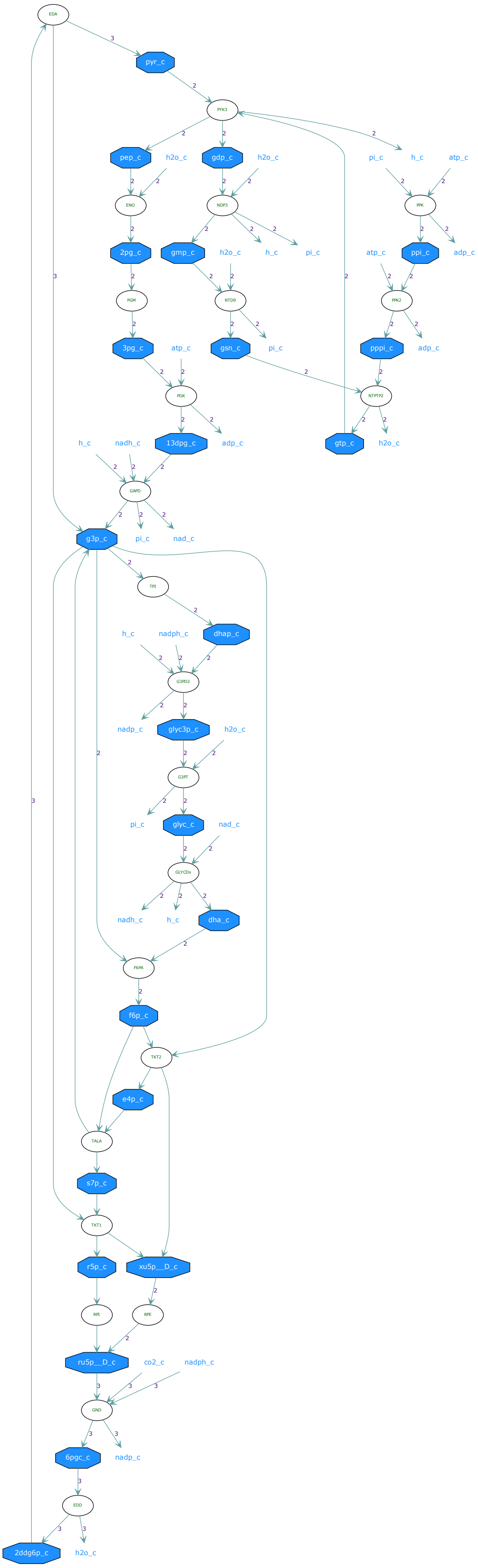
EDA
pyr_c
PYK3
pep_c
h2o_c
gdp_c
h2o_c
pi_c
h_c
atp_c
ENO
NDP3
PPK
2pg_c
gmp_c
h2o_c
h_c
pi_c
atp_c
ppi_c
adp_c
PGM
NTD9
PPK2
3pg_c
atp_c
gsn_c
pi_c
pppi_c
adp_c
PGK
NTPTP2
h_c
nadh_c
13dpg_c
adp_c
gtp_c
h2o_c
GAPD
g3p_c
pi_c
nad_c
TPI
h_c
nadph_c
dhap_c
G3PD2
nadp_c
glyc3p_c
h2o_c
G3PT
pi_c
glyc_c
nad_c
GLYCDx
nadh_c
h_c
dha_c
FBPA
f6p_c
TKT2
e4p_c
TALA
s7p_c
TKT1
r5p_c
xu5p__D_c
RPI
RPE
ru5p__D_c
co2_c
nadph_c
GND
6pgc_c
nadp_c
EDD
2ddg6p_c
h2o_c

### Pathway 22

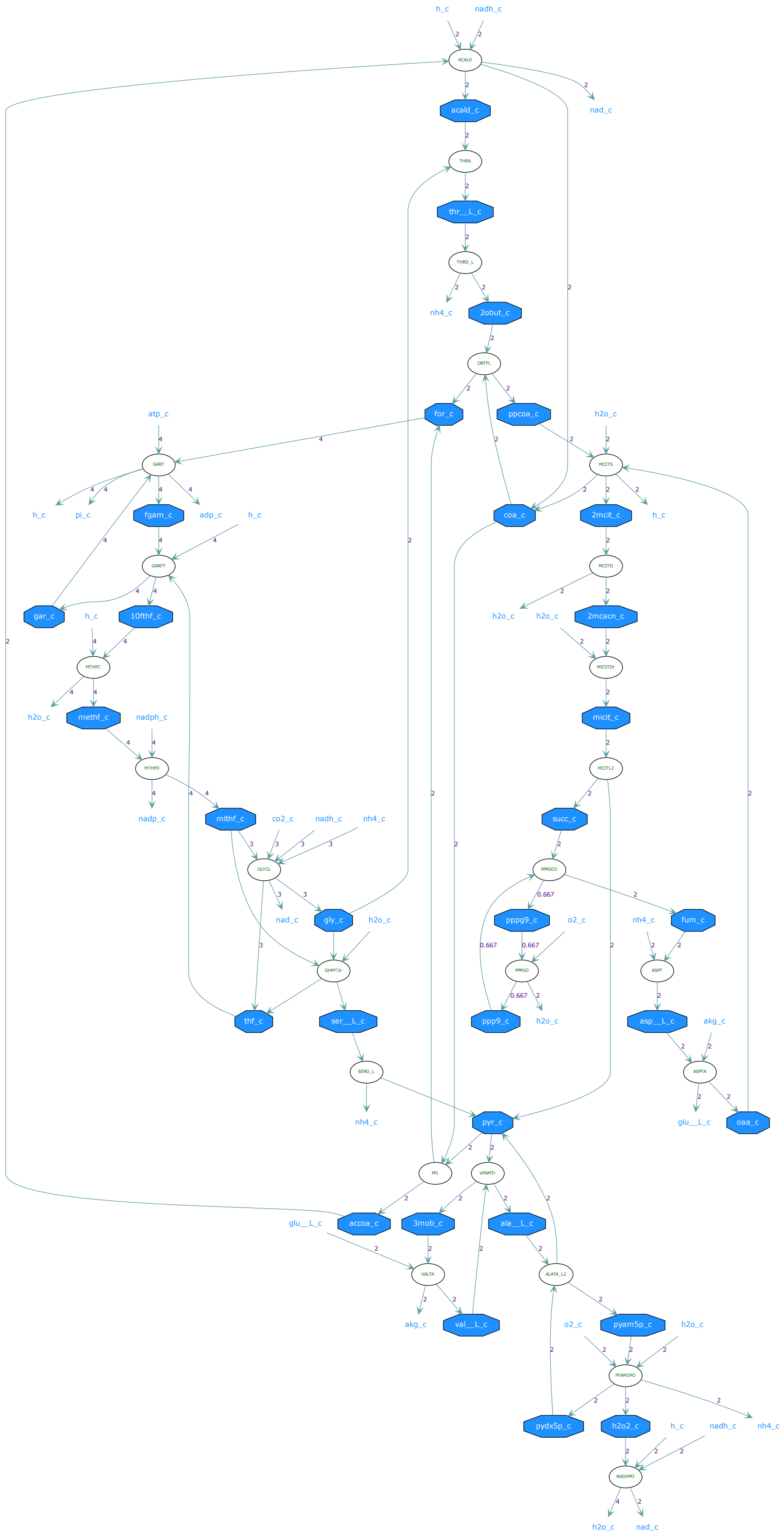

h_c
nadh_c
ACALD
nad_c
acald_c
THRA
thr__L_c
THRD_L
nh4_c
2obut_c
OBTFL
for_c
ppcoa_c
atp_c
h2o_c
GART
MCITS
h_c
pi_c
fgam_c
adp_c
h_c
coa_c
2mcit_c
h_c
GARFT
MCITD
gar_c
h_c
10fthf_c
h2o_c
h2o_c
2mcacn_c
MTHFC
MICITDr
h2o_c
methf_c
nadph_c
micit_c
MTHFD
MCITL2
nadp_c
mlthf_c
co2_c
nadh_c
nh4_c
succ_c
GLYCL
PPPGO3
nad_c
gly_c
h2o_c
pppg9_c
o2_c
nh4_c
fum_c
GHMT2r
PPPGO
ASPT
thf_c
ser__L_c
ppp9_c
h2o_c
asp__L_c
akg_c
SERD_L
ASPTA
nh4_c
pyr_c
glu__L_c
oaa_c
PFL
VPAMTr
glu__L_c
accoa_c
3mob_c
ala__L_c
VALTA
ALATA_L2
akg_c
val__L_c
o2_c
pyam5p_c
h2o_c
PYAM5PO
pydx5p_c
h2o2_c
h_c
nadh_c
nh4_c
NADH9O
h2o_c
nad_c

### Pathway 23

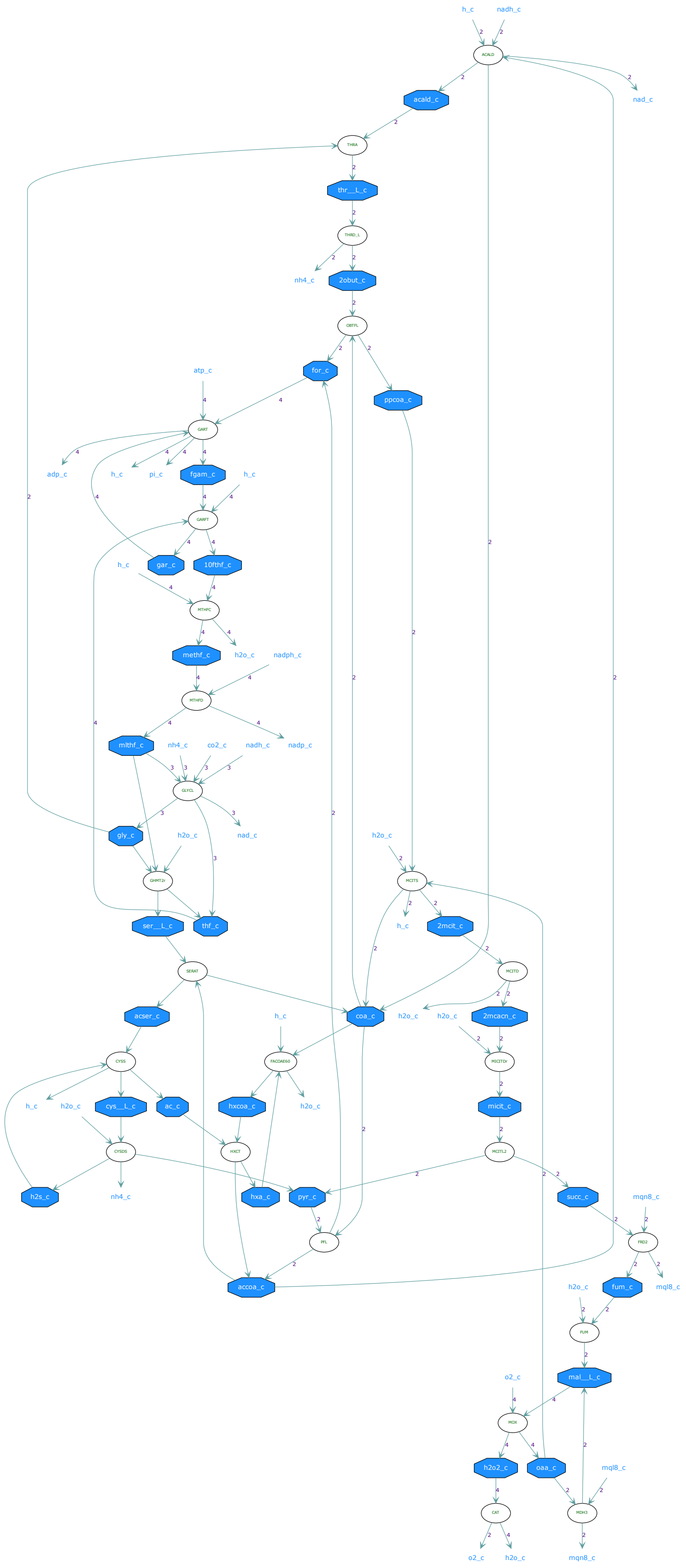
h_c
nadh_c
ACALD
nad_c
acald_c
THRA
thr__L_c
THRD_L
nh4_c
2obut_c
OBTFL
for_c
ppcoa_c
atp_c
GART
adp_c
h_c
pi_c
fgam_c
h_c
GARFT
gar_c
10fthf_c
h_c
MTHFC
methf_c
h2o_c
nadph_c
MTHFD
mlthf_c
nh4_c
co2_c
nadh_c
nadp_c
GLYCL
gly_c
h2o_c
nad_c
h2o_c
MCITS
GHMT2r
ser__L_c
thf_c
h_c
2mcit_c
SERAT
MCITD
acser_c
h_c
coa_c
h2o_c
h2o_c
2mcacn_c
CYSS
FACOAE60
MICITDr
h_c
h2o_c
cys__L_c
ac_c
hxcoa_c
h2o_c
micit_c
CYSDS
HXCT
MCITL2
h2s_c
nh4_c
hxa_c
pyr_c
succ_c
mqn8_c
PFL
FRD2
accoa_c
h2o_c
fum_c
mql8_c
FUM
mal__L_c
o2_c
MOX
h2o2_c
oaa_c
mql8_c
CAT
MDH3
o2_c
h2o_c
mqn8_c

### Pathway 24

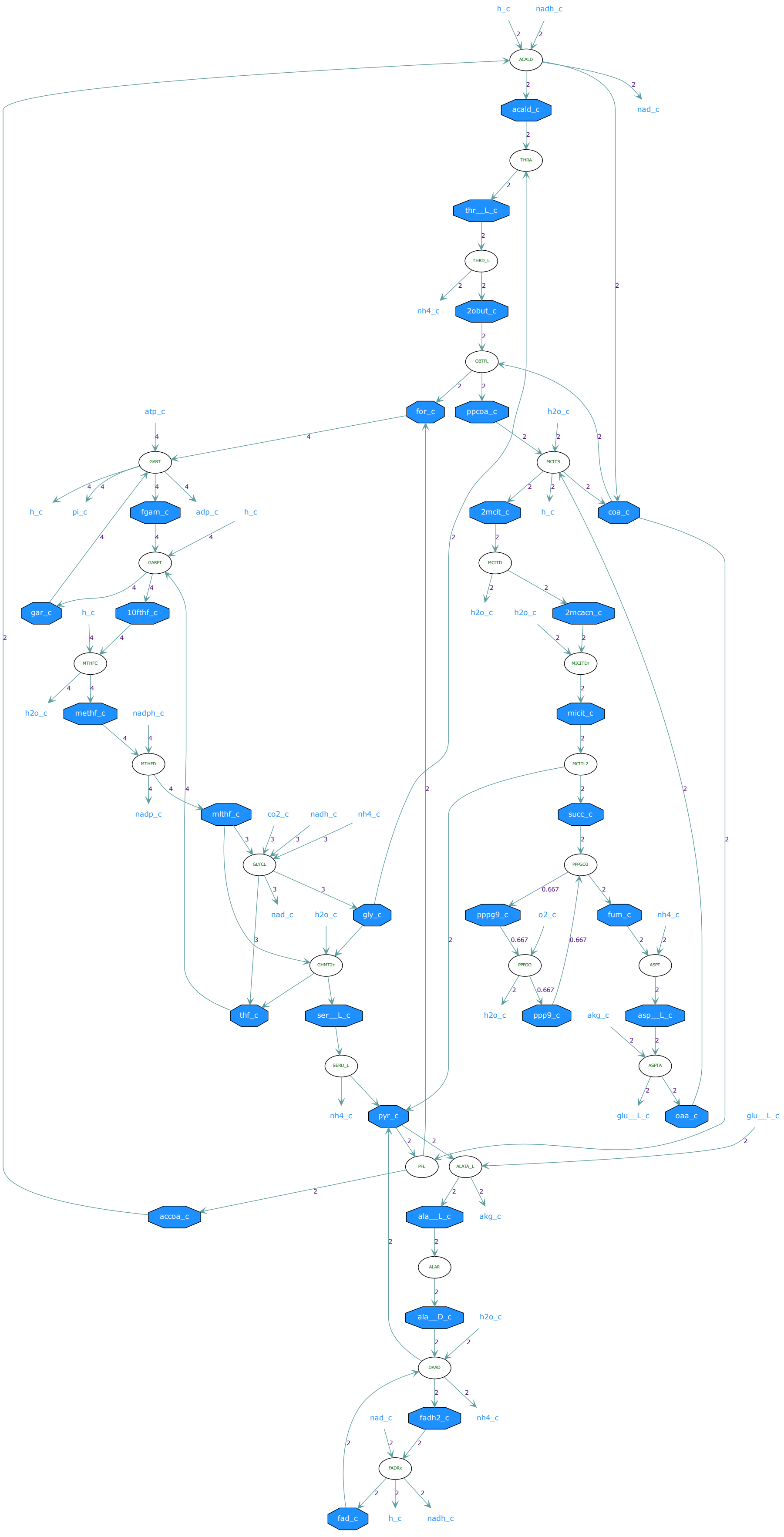
h_c
nadh_c
ACALD
nad_c
acald_c
THRA
thr__L_c
THRD_L
nh4_c
2obut_c
OBTFL
for_c
ppcoa_c
h2o_c
atp_c
GART
MCITS
h_c
pi_c
fgam_c
adp_c
h_c
2mcit_c
h_c
coa_c
GARFT
MCITD
gar_c
h_c
10fthf_c
h2o_c
h2o_c
2mcacn_c
MTHFC
MICITDr
h2o_c
methf_c
nadph_c
micit_c
MTHFD
MCITL2
nadp_c
mlthf_c
co2_c
nadh_c
nh4_c
succ_c
GLYCL
PPPGO3
nad_c
h2o_c
gly_c
pppg9_c
o2_c
fum_c
nh4_c
GHMT2r
PPPGO
ASPT
thf_c
ser__L_c
h2o_c
ppp9_c
akg_c
asp__L_c
SERD_L
ASPTA
nh4_c
pyr_c
glu__L_c
oaa_c
glu__L_c
PFL
ALATA_L
accoa_c
ala__L_c
akg_c
ALAR
ala__D_c
h2o_c
DAAD
nad_c
fadh2_c
nh4_c
FADRx
fad_c
h_c
nadh_c

### Pathway 25

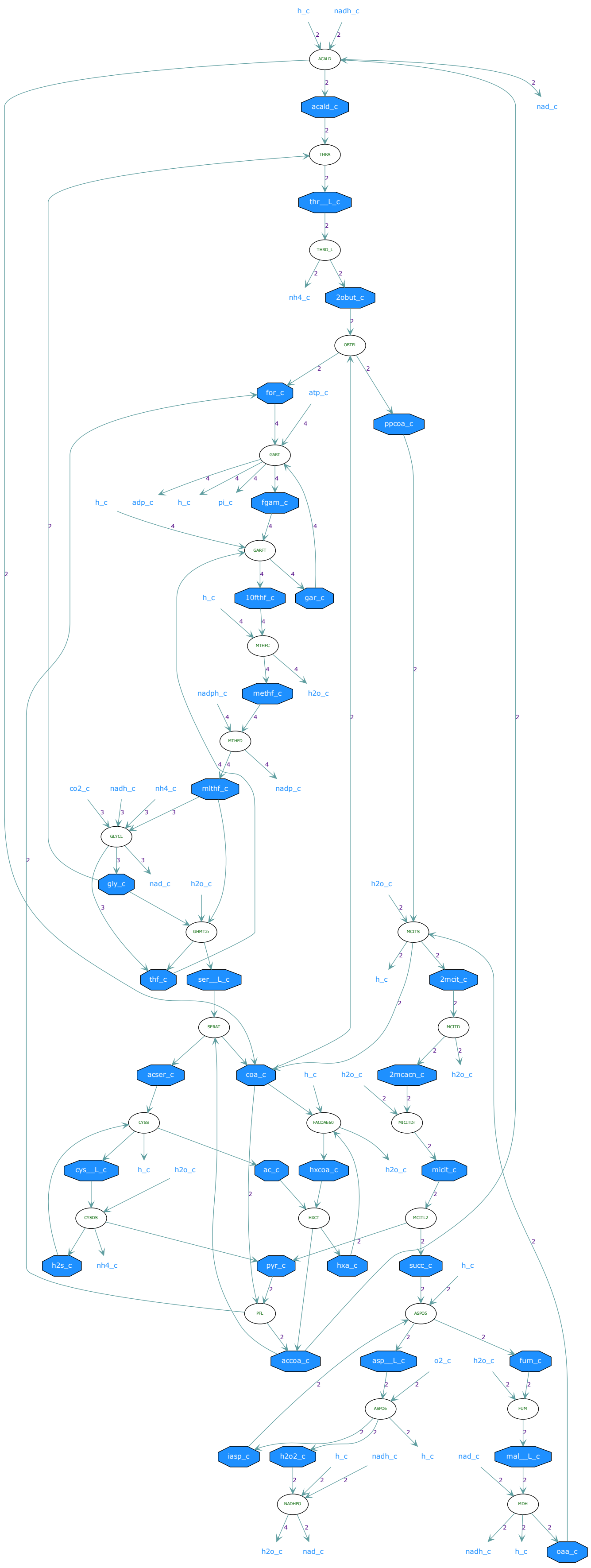
h_c
nadh_c
ACALD
nad_c
acald_c
THRA
thr__L_c
THRD_L
nh4_c
2obut_c
OBTFL
for_c
atp_c
ppcoa_c
GART
h_c
adp_c
h_c
pi_c
fgam_c
GARFT
h_c
10fthf_c
gar_c
MTHFC
nadph_c
methf_c
h2o_c
MTHFD
co2_c
nadh_c
nh4_c
mlthf_c
nadp_c
GLYCL
gly_c
nad_c
h2o_c
h2o_c
MCITS
GHMT2r
h_c
2mcit_c
thf_c
ser__L_c
SERAT
MCITD
acser_c
coa_c
h_c
h2o_c
2mcacn_c
h2o_c
CYSS
FACOAE60
MICITDr
cys__L_c
h_c
h2o_c
ac_c
hxcoa_c
h2o_c
micit_c
CYSDS
HXCT
MCITL2
h2s_c
nh4_c
pyr_c
hxa_c
succ_c
h_c
PFL
ASPO5
accoa_c
asp__L_c
o2_c
h2o_c
fum_c
ASPO6
FUM
iasp_c
h2o2_c
h_c
nadh_c
h_c
nad_c
mal__L_c
NADHPO
MDH
h2o_c
nad_c
nadh_c
h_c
oaa_c

### Pathway 26

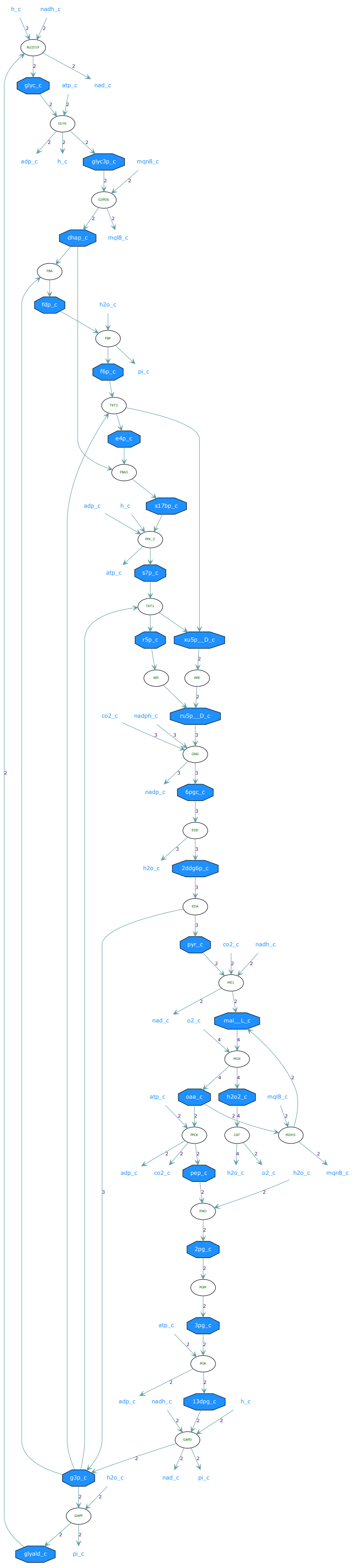

### Pathway 27

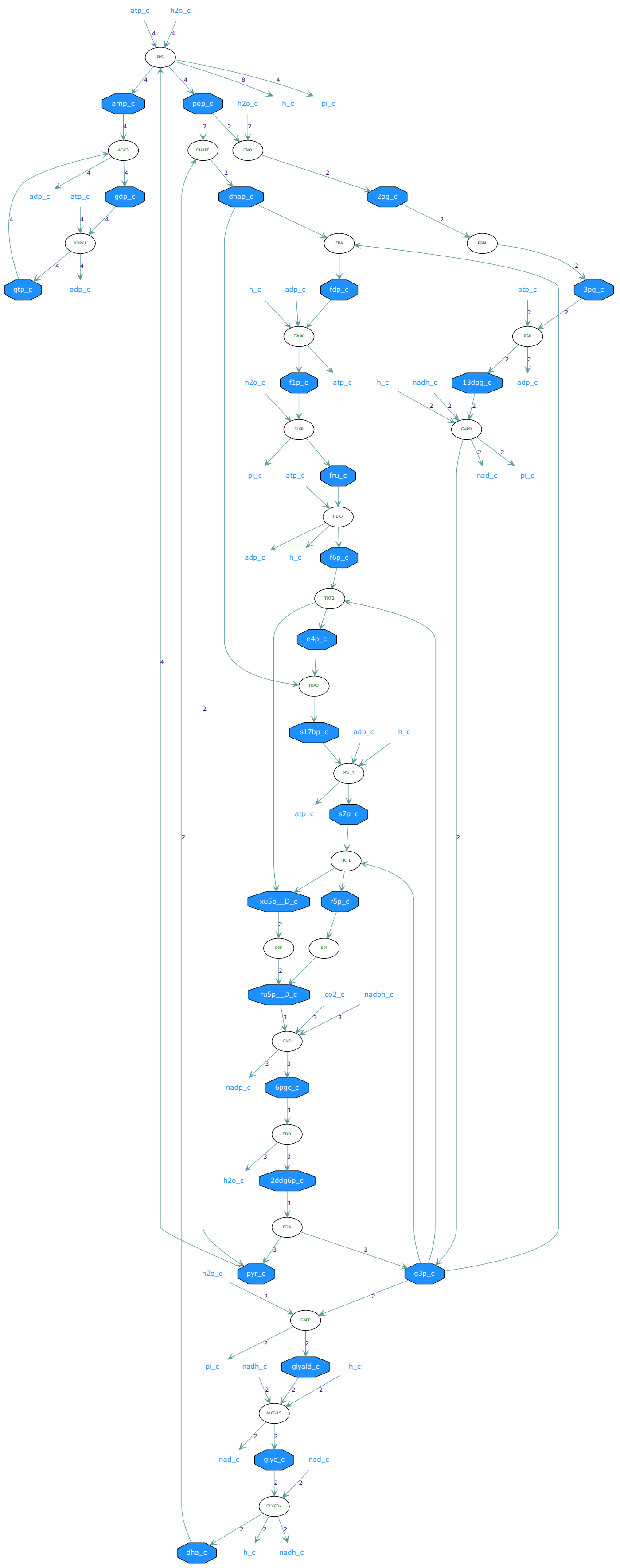
atp_c
h2o_c
PPS
amp_c
pep_c
h2o_c
h_c
pi_c
ADK3
DHAPT
ENO
adp_c
atp_c
gdp_c
dhap_c
2pg_c
NDPK1
FBA
PGM
gtp_c
adp_c
h_c
adp_c
fdp_c
atp_c
3pg_c
FRUK
PGK
h2o_c
f1p_c
atp_c
h_c
nadh_c
13dpg_c
adp_c
F1PP
GAPD
pi_c
atp_c
fru_c
nad_c
pi_c
HEX7
adp_c
h_c
f6p_c
TKT2
e4p_c
FBA3
s17bp_c
adp_c
h_c
PFK_3
atp_c
s7p_c
TKT1
xu5p__D_c
r5p_c
RPE
RPI
ru5p__D_c
co2_c
nadph_c
GND
nadp_c
6pgc_c
EDD
h2o_c
2ddg6p_c
EDA
h2o_c
pyr_c
g3p_c
GAPP
pi_c
nadh_c
glyald_c
h_c
ALCD19
nad_c
glyc_c
nad_c
GLYCDx
dha_c
h_c
nadh_c

### Pathway 28

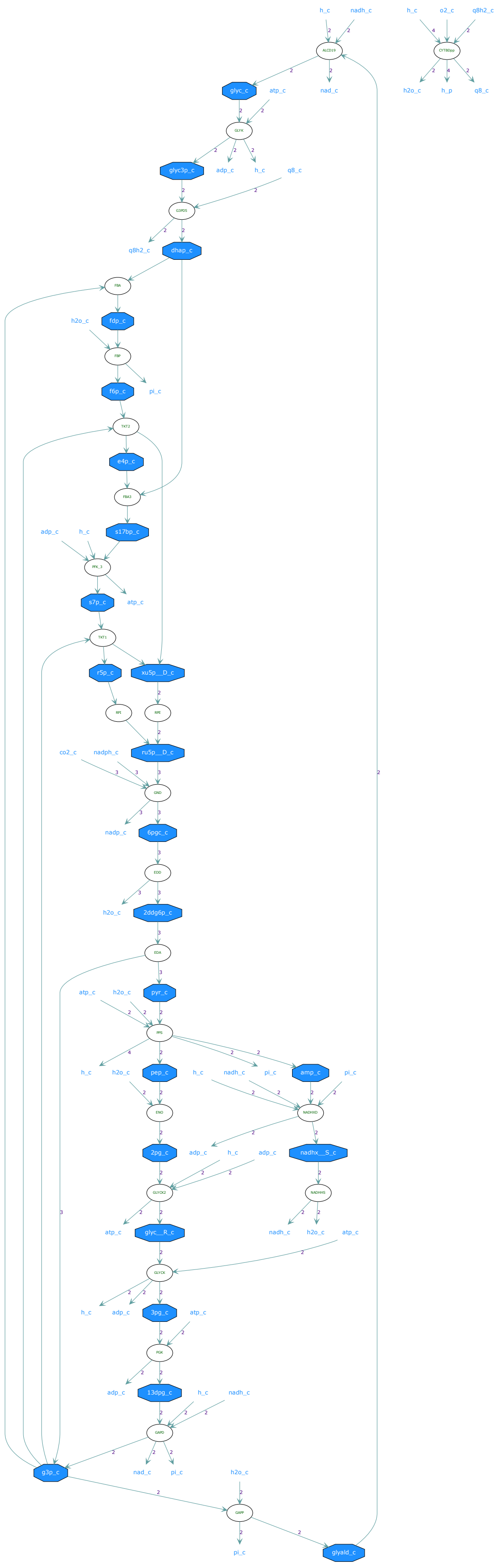
h_c
nadh_c
o2_c
q8h2_c
ALDD19
CYTBDpp
glyc_c
atp_c
nad_c
h2o_c
h_p
q8_c
GLYK
glyc3p_c
adp_c
G3PD5
q8h2_c
dhap_c
FBA
fdp_c
FBP
f6p_c
pi_c
TKT2
e4p_c
FBA3
s17bp_c
PFK_3
s7p_c
TKT1
r5p_c
xu5p__D_c
RPI
RPE
co2_c
nadph_c
ru5p__D_c
GND
nadp_c
6pgc_c
EDD
2ddg6p_c
EDA
pyr_c
PPS
pep_c
amp_c
ENO
NADH5
2pg_c
nadhx__S_c
GLYCK2
NADHHS
glyc__R_c
GLYCK
3pg_c
PGK
13dpg_c
GAPD
g3p_c
GAPP
glyald_c

### Pathway 29

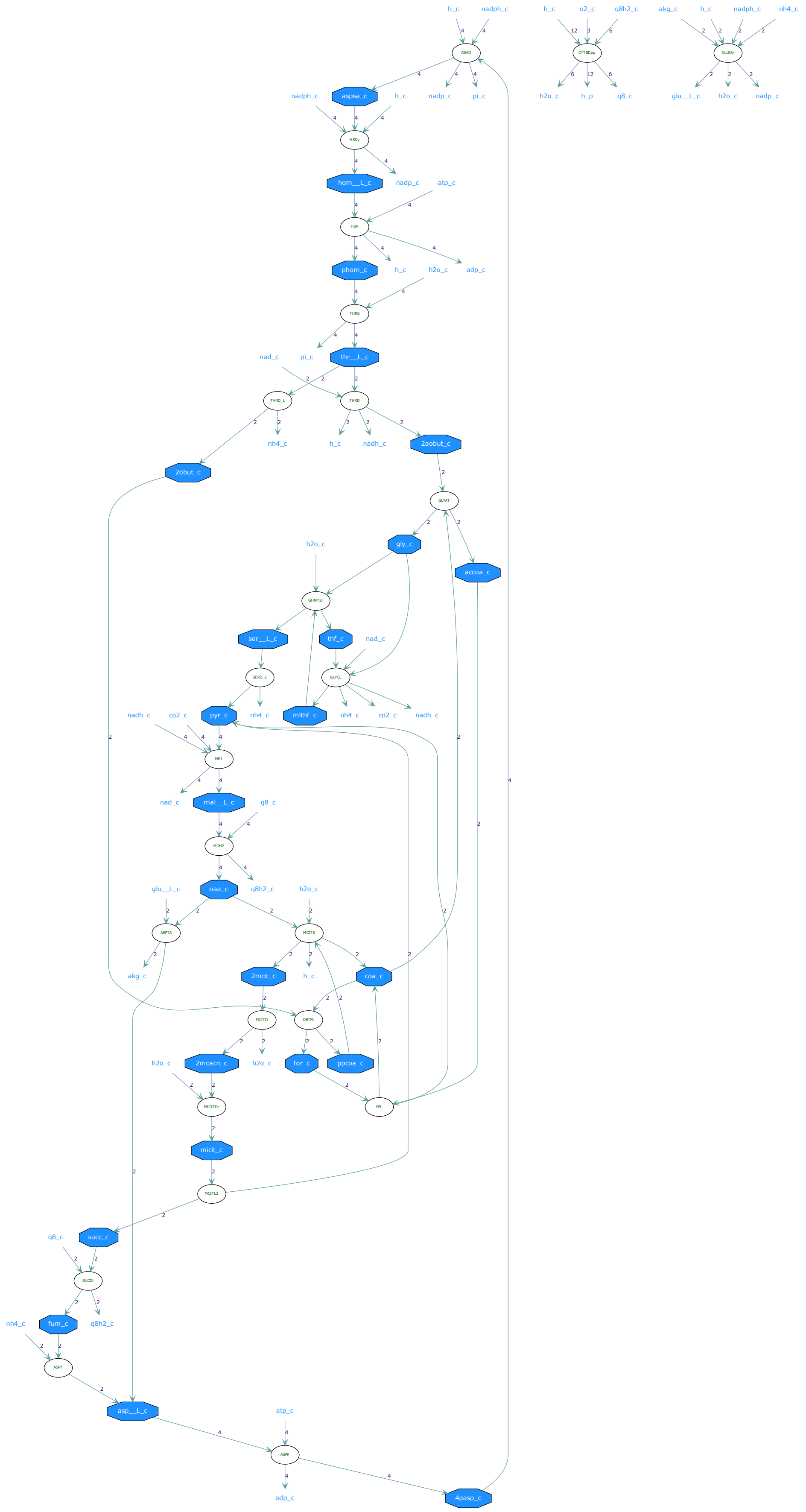
h_c
nadph_c
ASAD
aspsa_c
nadp_c
pi_c
HSDy
hom__L_c
atp_c
HSK
phom_c
h2o_c
adp_c
THRS
nad_c
thr__L_c
THRD_L
THRD
nh4_c
nadh_c
2aobut_c
2obut_c
GLYAT
gly_c
accoa_c
GHMT2r
ser__L_c
thf_c
SERD_L
GLYCL
pyr_c
mlthf_c
co2_c
ME1
nad_c
mal__L_c
q8_c
MDH2
oaa_c
q8h2_c
glu__L_c
ASPTA
MCITS
akg_c
2mcit_c
coa_c
MCITD
OBTFL
2mcacn_c
for_c
ppcoa_c
MCITDr
PFL
micit_c
MICITL2
succ_c
SUCDi
fum_c
ASPT
asp__L_c
ASPK
4pasp_c
o2_c
CYTBDpp
h_p
GLUDy

### Pathway 30

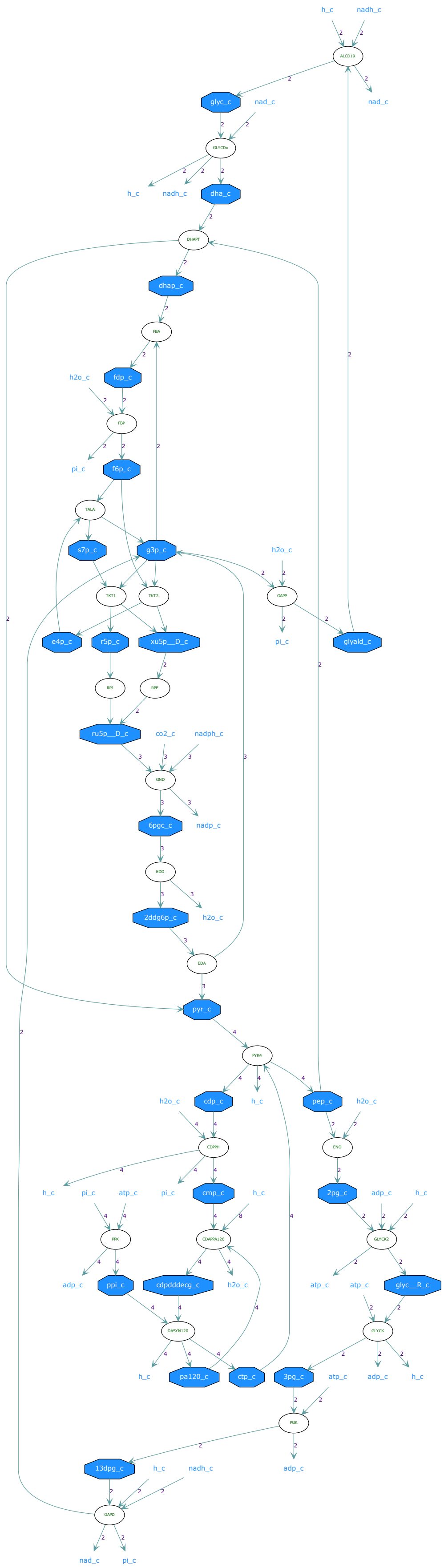

h_c
nadh_c
ALCD19
glyc_c
nad_c
GLYCDx
dha_c
DHAPT
dhap_c
FBA
fdp_c
h2o_c
FBP
pi_c
f6p_c
TALA
s7p_c
g3p_c
TKT1
TKT2
GAPP
e4p_c
r5p_c
xu5p__D_c
glyald_c
RPI
RPE
ru5p__D_c
co2_c
nadph_c
GND
6pgc_c
nadp_c
EDD
2ddg6p_c
EDA
pyr_c
PYK4
cdp_c
pep_c
CDPPH
ENO
cmp_c
2pg_c
atp_c
adp_c
PPK
CDAPPA120
GLYCK2
ppi_c
cdpdddecg_c
glyc__R_c
DASYN120
GLYCK
pa120_c
ctp_c
3pg_c
PGK
13dpg_c
GAPD

### Pathway 31

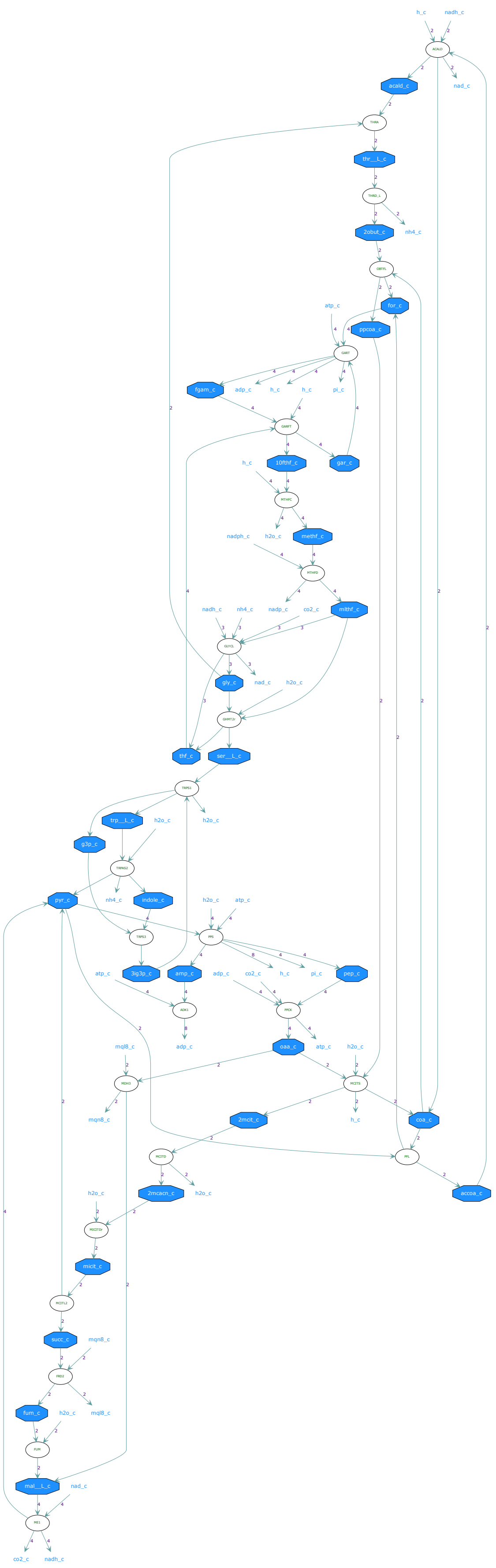

h_c
nadh_c
ACALD
acald_c
nad_c
THRA
thr__L_c
THRD_L
2obut_c
nh4_c
OBTFL
for_c
ppcoa_c
atp_c
GART
fgam_c
adp_c
h_c
h_c
pi_c
GARFT
10fthf_c
gar_c
h_c
MTHFC
h2o_c
methf_c
nadph_c
MTHFD
mlthf_c
nadh_c
nh4_c
nadp_c
co2_c
GCVCL
gly_c
nad_c
h2o_c
GHMT2r
thf_c
ser__L_c
TRPS1
trp__L_c
h2o_c
h2o_c
g3p_c
TRPAS2
pyr_c
nh4_c
indole_c
TRPS3
PPS
h2o_c
atp_c
3ig3p_c
amp_c
atp_c
adp_c
co2_c
h_c
pi_c
pep_c
ADK1
PPCK
adp_c
oaa_c
atp_c
mql8_c
h2o_c
MDH3
MCITS
mqn8_c
h_c
2mcit_c
coa_c
MCITD
PFL
2mcacn_c
h2o_c
accoa_c
h2o_c
MICITDr
micit_c
MCITL2
succ_c
mqn8_c
FRD2
fum_c
h2o_c
mql8_c
FUM
mal__L_c
nad_c
ME1
co2_c
nadh_c

### Pathway 32

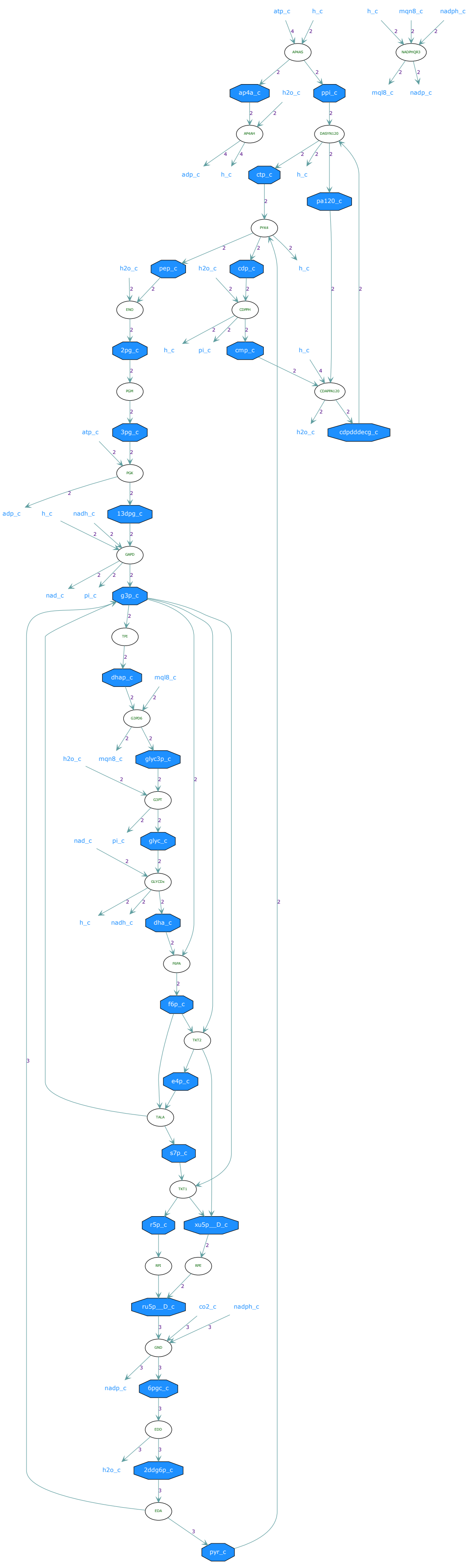
atp_c
h_c
h_c
mqn8_c
nadph_c
ap4a_c
h2o_c
ppi_c
mql8_c
nadp_c
adp_c
h_c
ctp_c
h_c
pa120_c
h2o_c
pep_c
h2o_c
cdp_c
h_c
h_c
pi_c
2pg_c
cmp_c
h_c
3pg_c
atp_c
h2o_c
cdpdddecg_c
adp_c
h_c
nadh_c
13dpg_c
nad_c
pi_c
g3p_c
dhap_c
mql8_c
h2o_c
mqn8_c
glyc3p_c
nad_c
pi_c
glyc_c
h_c
nadh_c
dha_c
f6p_c
e4p_c
s7p_c
r5p_c
xu5p__D_c
ru5p__D_c
co2_c
nadph_c
nadp_c
6pgc_c
h2o_c
2ddg6p_c
pyr_c

### Pathway 33

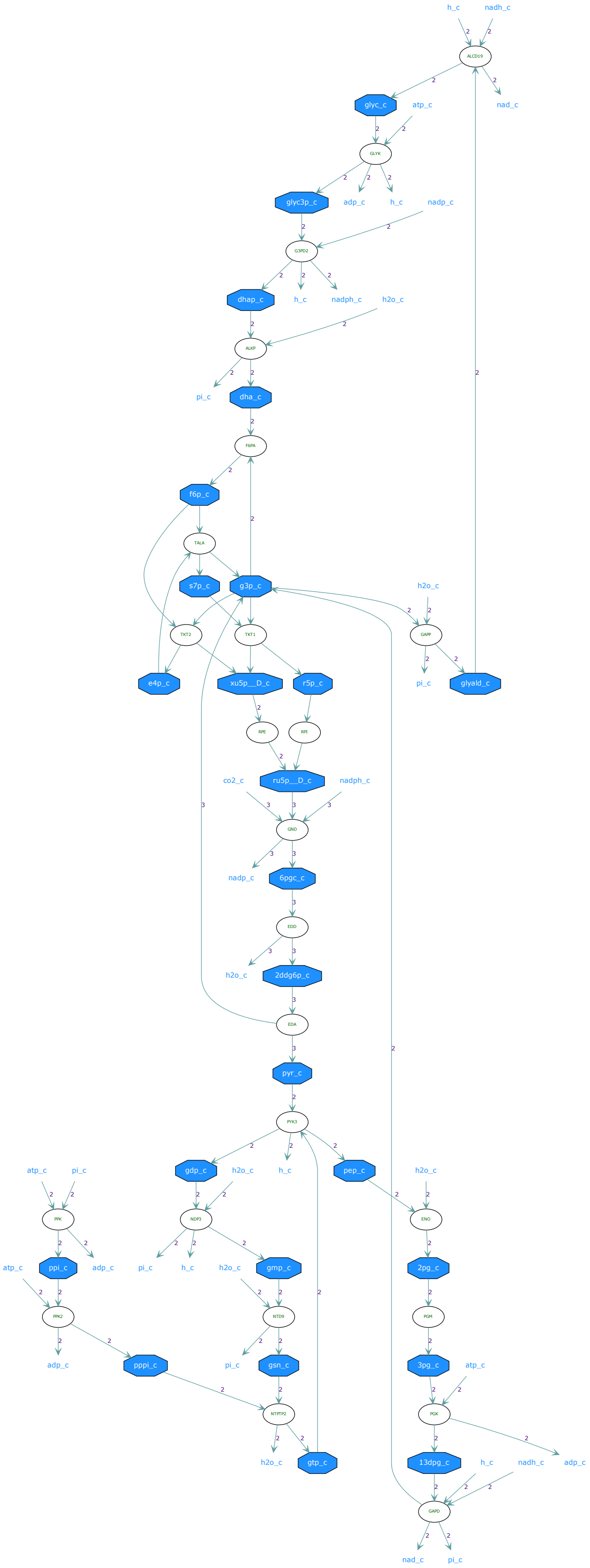

h_c
nadh_c
ALCD19
nad_c
glyc_c
atp_c
GLYK
glyc3p_c
adp_c
h_c
nadp_c
G3PD2
dhap_c
h_c
nadph_c
h2o_c
ALKP
pi_c
dha_c
F6PA
f6p_c
TALA
s7p_c
g3p_c
h2o_c
TKT2
TKT1
GAPP
e4p_c
xu5p__D_c
r5p_c
pi_c
glyald_c
RPE
RPI
co2_c
ru5p__D_c
nadph_c
GND
nadp_c
6pgc_c
EDD
h2o_c
2ddg6p_c
EDA
pyr_c
PYK3
atp_c
pi_c
gdp_c
h2o_c
h_c
pep_c
h2o_c
PPK
NDP3
ENO
atp_c
ppi_c
adp_c
pi_c
h_c
h2o_c
gmp_c
2pg_c
PPK2
NTD9
PGM
adp_c
pppi_c
pi_c
gsn_c
3pg_c
atp_c
NTPTP2
PGK
h2o_c
gtp_c
13dpg_c
h_c
nadh_c
adp_c
GAPD
nad_c
pi_c

### Pathway 34

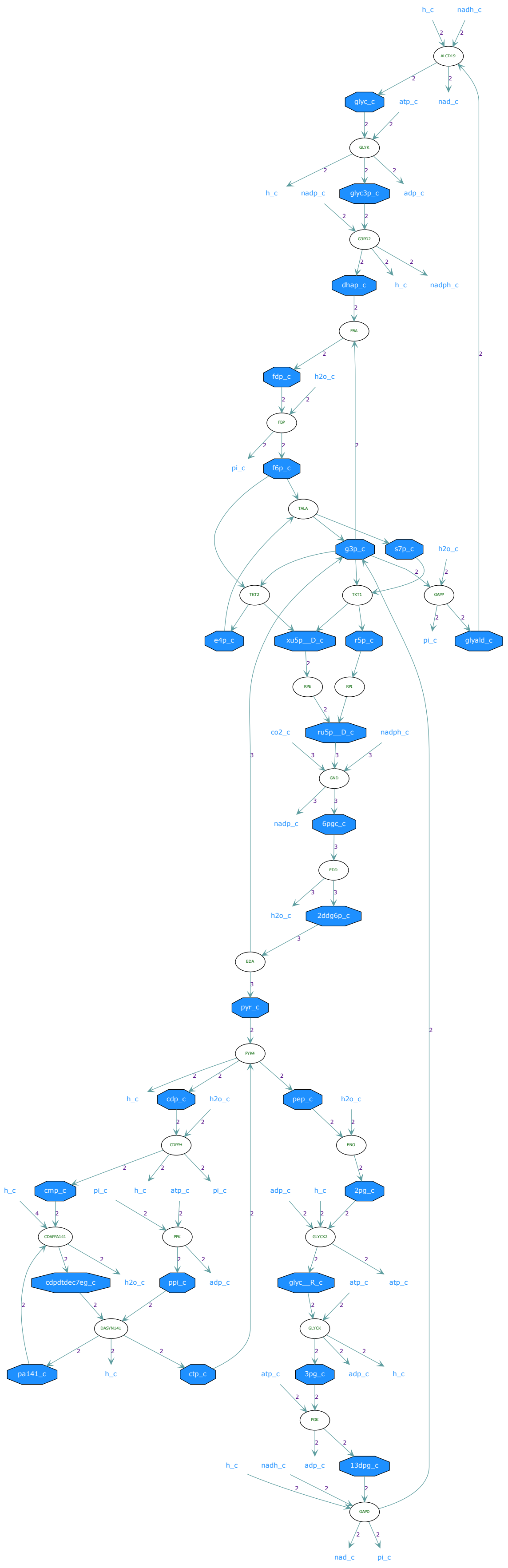
h_c
nadh_c
ALCD19
glyc_c
atp_c
nad_c
GLYK
h_c
nadp_c
glyc3p_c
adp_c
G3PD2
dhap_c
h_c
nadph_c
FBA
fdp_c
h2o_c
FBP
pi_c
f6p_c
TALA
g3p_c
s7p_c
h2o_c
TKT2
TKT1
GAPP
e4p_c
xu5p__D_c
r5p_c
pi_c
glyald_c
RPE
RPI
co2_c
ru5p__D_c
nadph_c
GND
nadp_c
6pgc_c
EDD
h2o_c
2ddg6p_c
EDA
pyr_c
PYK4
h_c
cdp_c
h2o_c
pep_c
h2o_c
CDPPH
ENO
h_c
cmp_c
pi_c
h_c
atp_c
pi_c
adp_c
h_c
2pg_c
CDAPPA141
PPK
GLYCK2
cdpdtdec7eg_c
h2o_c
ppi_c
adp_c
glyc__R_c
atp_c
atp_c
DASYN141
GLYCK
pa141_c
h_c
ctp_c
atp_c
3pg_c
adp_c
h_c
PGK
h_c
nadh_c
adp_c
13dpg_c
GAPD
nad_c
pi_c

### Pathway 35

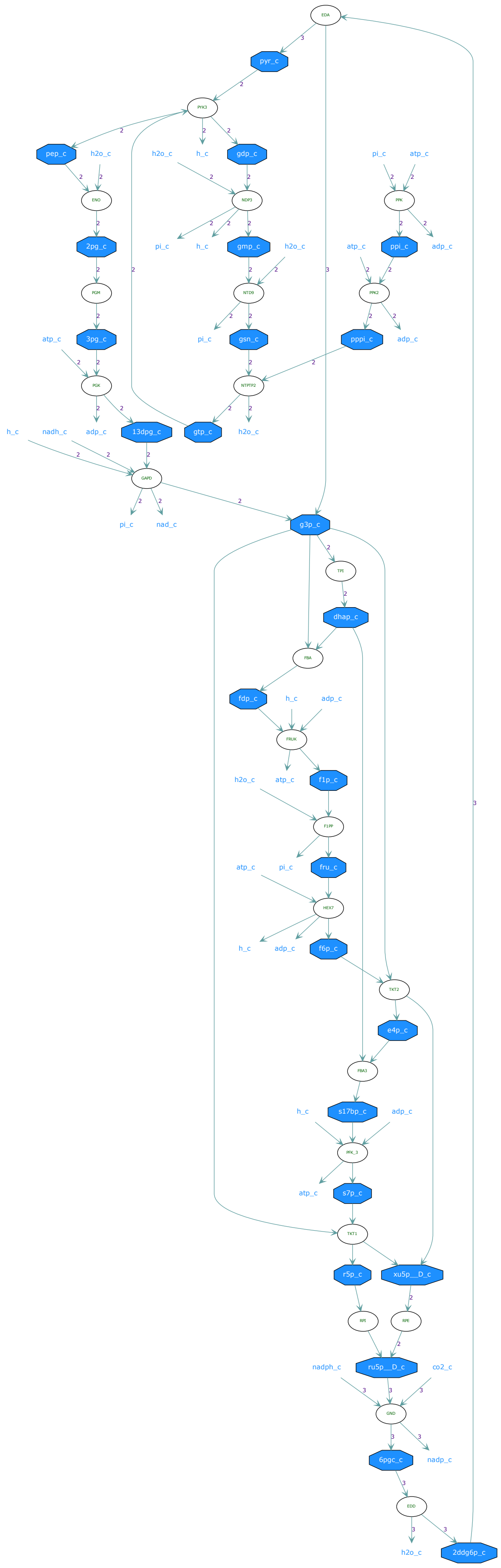
EDA
pyr_c
PYK3
pep_c
h2o_c
h_c
gdp_c
pi_c
atp_c
ENO
NDP3
PPK
2pg_c
gmp_c
ppi_c
adp_c
PGM
NTD9
PPK2
3pg_c
gsn_c
pppi_c
PGK
NTPTP2
nadh_c
13dpg_c
gtp_c
GAPD
nad_c
g3p_c
TPI
dhap_c
FBA
fdp_c
FRUK
f1p_c
F1PP
fru_c
HEX7
f6p_c
TKT2
e4p_c
FBA3
s17bp_c
PFK_3
s7p_c
TKT1
r5p_c
xu5p__D_c
RPI
RPE
nadph_c
ru5p__D_c
co2_c
GND
6pgc_c
nadp_c
EDD
2ddg6p_c

### Pathway 36

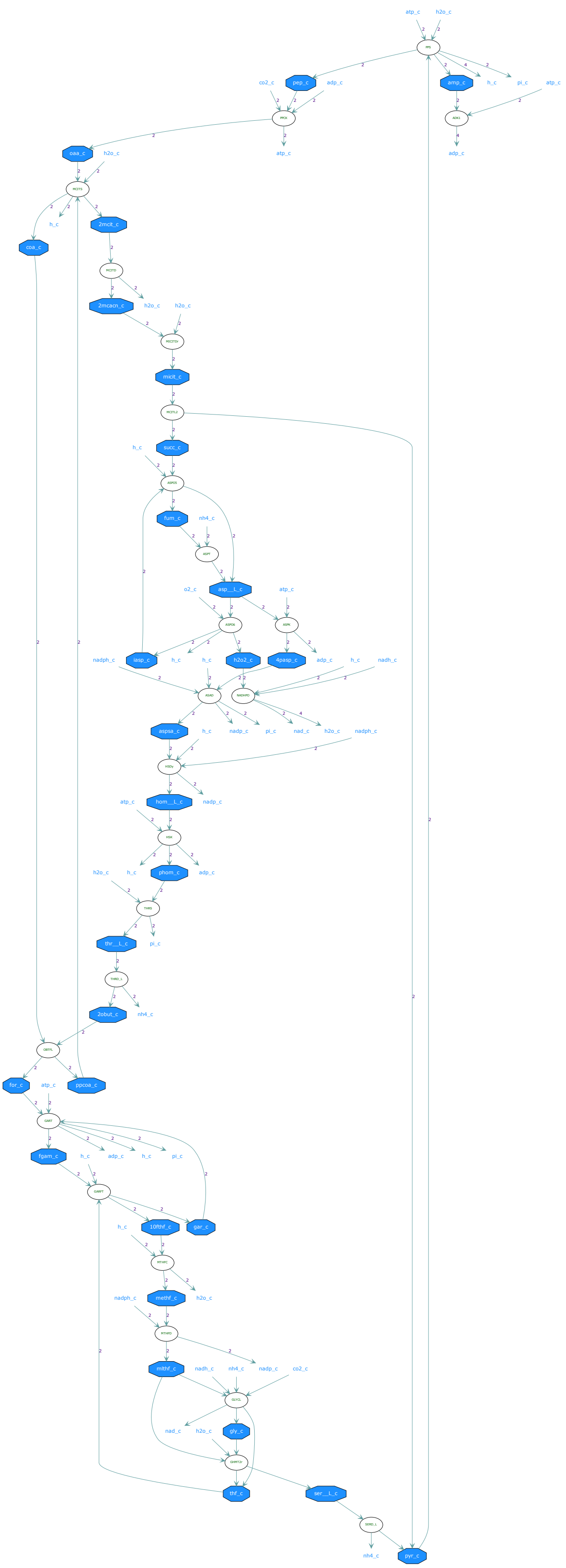

### Pathway 37

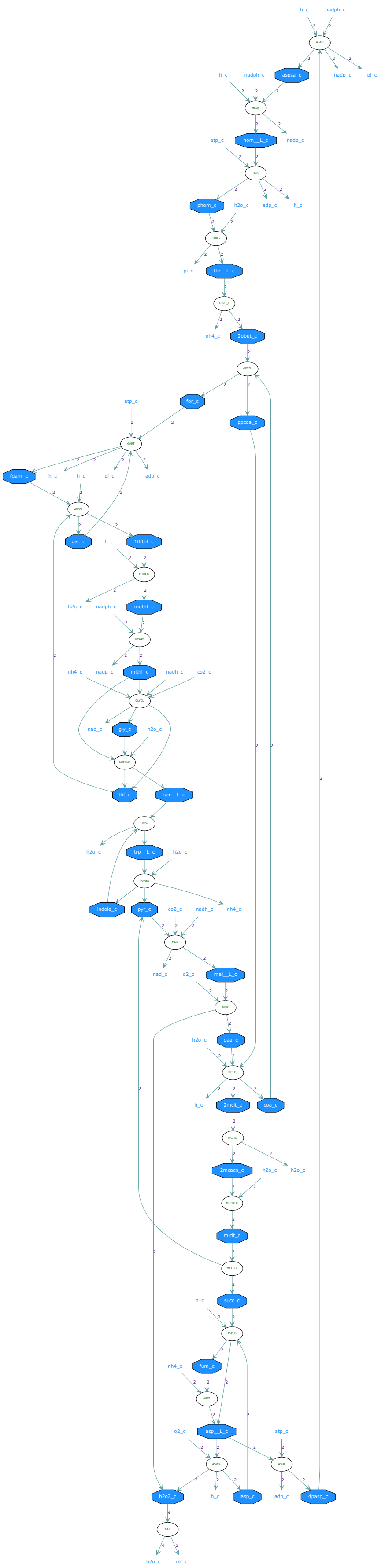

h_c
nadph_c
aspsa_c
nadp_c
pi_c
hom__L_c
atp_c
phom_c
h2o_c
adp_c
thr__L_c
nh4_c
2obut_c
for_c
ppcoa_c
fgam_c
gar_c
10fthf_c
methf_c
mlthf_c
nadh_c
co2_c
nad_c
gly_c
thf_c
ser__L_c
trp__L_c
indole_c
pyr_c
o2_c
mal__L_c
oaa_c
2mcit_c
coa_c
2mcacn_c
micit_c
succ_c
fum_c
asp__L_c
iasp_c
4pasp_c
h2o2_c

### Pathway 38

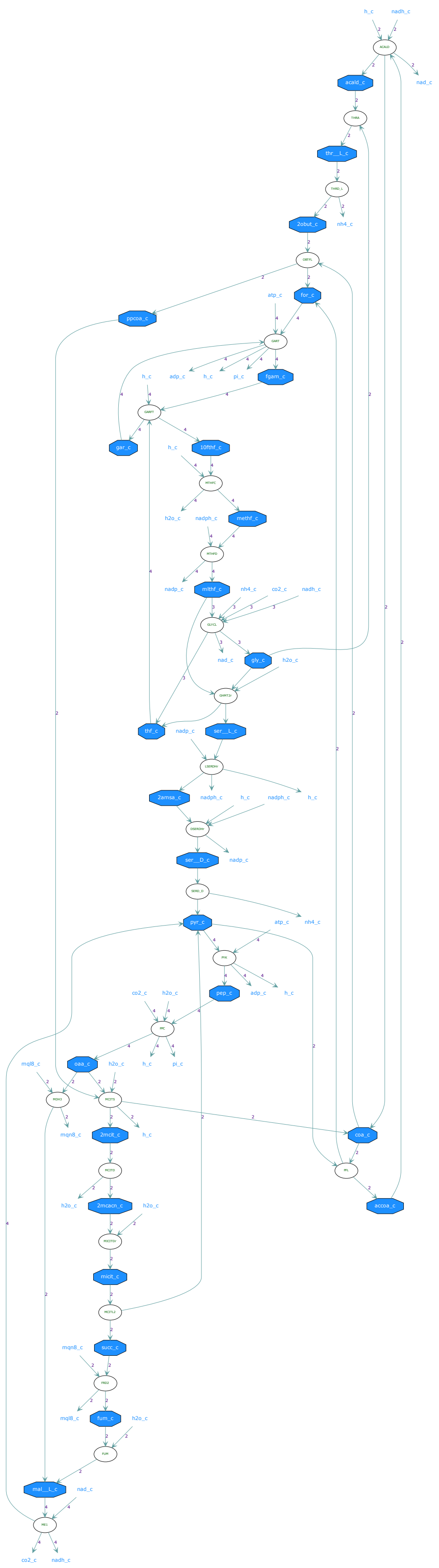

### Pathway 39

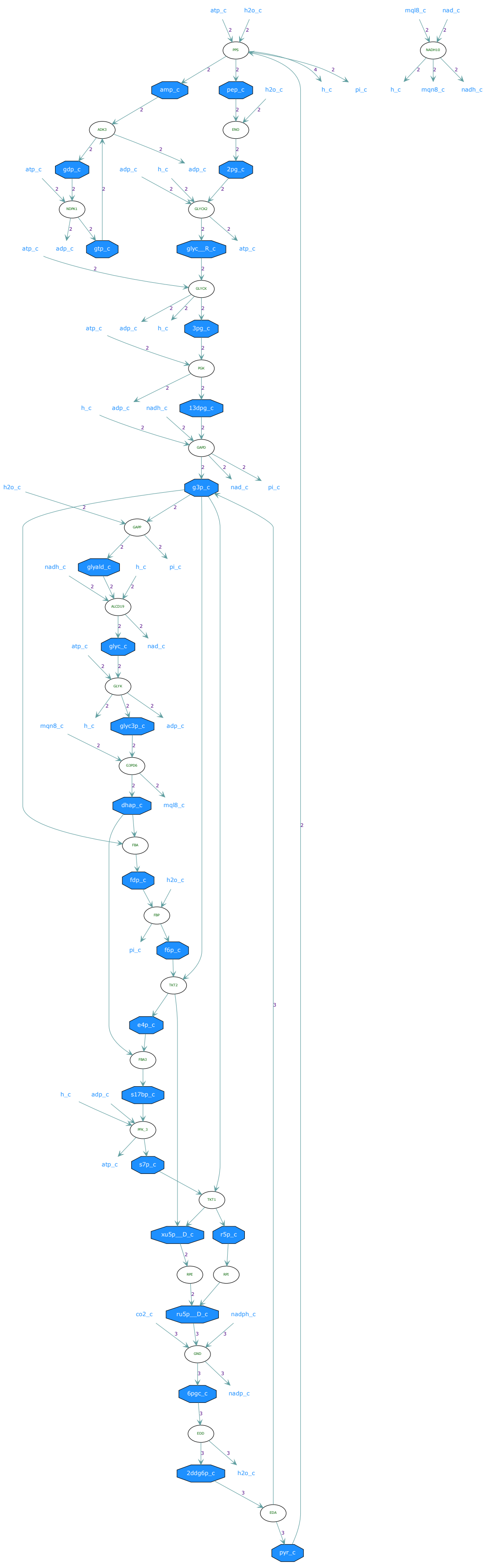

### Pathway 40

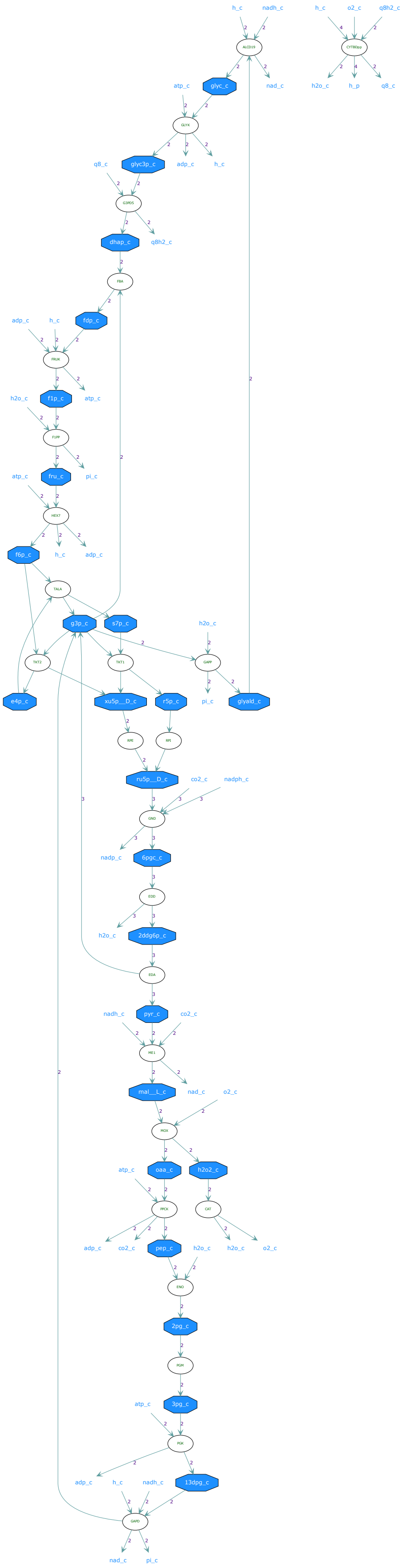

h_c
nadh_c
o2_c
q8h2_c
nad_c
glyc_c
atp_c
h2o_c
h_p
q8_c
glyc3p_c
adp_c
dhap_c
q8h2_c
fdp_c
f1p_c
pi_c
fru_c
f6p_c
g3p_c
s7p_c
e4p_c
xu5p__D_c
r5p_c
glyald_c
ru5p__D_c
co2_c
nadph_c
nadp_c
6pgc_c
2ddg6p_c
pyr_c
mal__L_c
oaa_c
h2o2_c
pep_c
2pg_c
3pg_c
13dpg_c

### Pathway 41

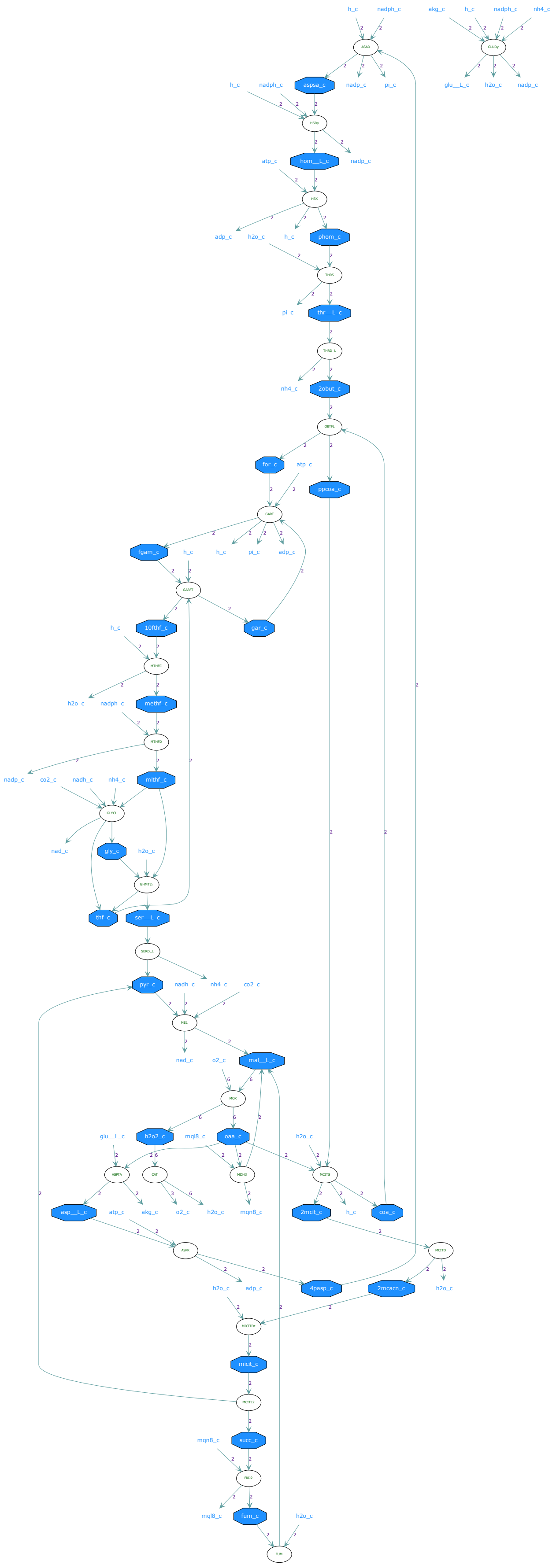

h_c
nadph_c
akg_c
h_c
nadph_c
nh4_c
aspsa_c
nadp_c
pi_c
glu__L_c
h2o_c
nadp_c
hom__L_c
atp_c
adp_c
h2o_c
h_c
phom_c
pi_c
thr__L_c
nh4_c
2obut_c
for_c
atp_c
ppcoa_c
fgam_c
h_c
pi_c
adp_c
10fthf_c
gar_c
methf_c
h2o_c
nadph_c
nadp_c
co2_c
nadh_c
nh4_c
mlthf_c
nad_c
gly_c
h2o_c
thf_c
ser__L_c
pyr_c
nadh_c
nh4_c
co2_c
nad_c
o2_c
mal__L_c
h2o2_c
mql8_c
oaa_c
h2o_c
glu__L_c
asp__L_c
atp_c
akg_c
o2_c
h2o_c
mqn8_c
2mcit_c
h_c
coa_c
h2o_c
adp_c
4pasp_c
2mcacn_c
h2o_c
micit_c
succ_c
mqn8_c
mql8_c
fum_c
h2o_c

### Pathway 42

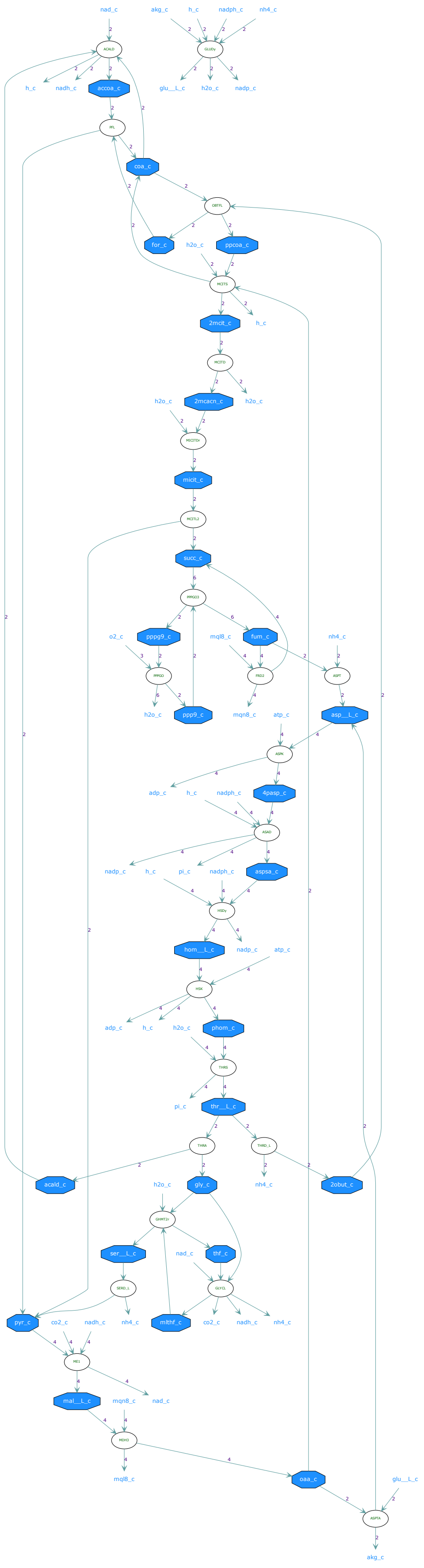

Pathway 43

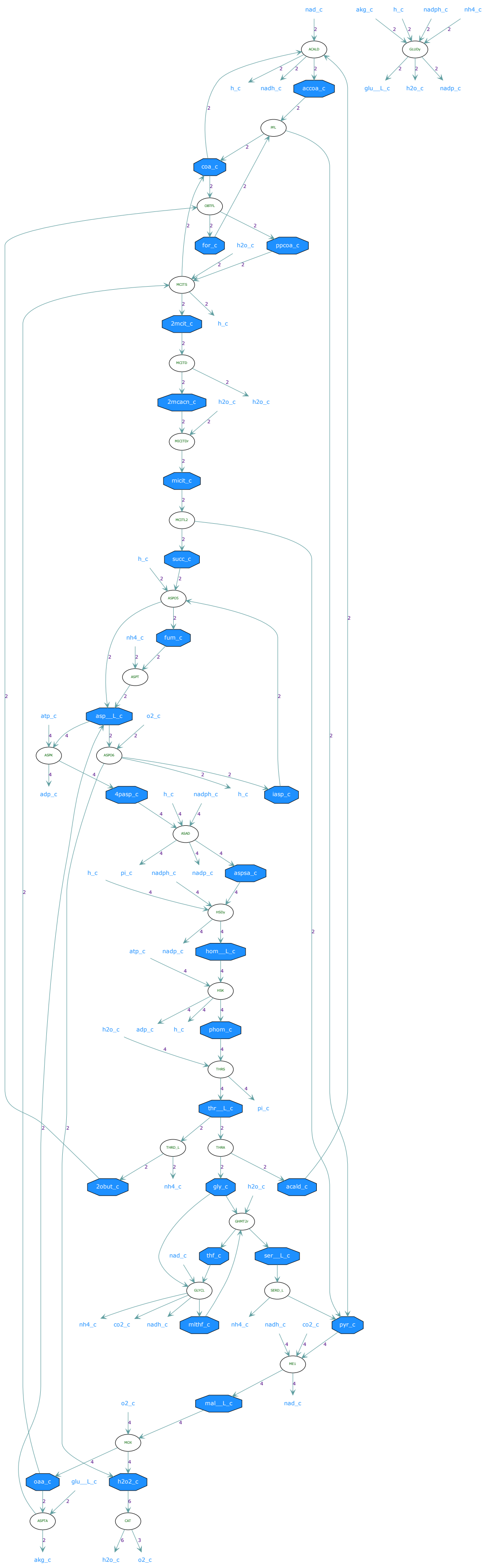
nad_c
akg_c
h_c
nadph_c
nh4_c
h_c
nadh_c
accoa_c
glu__L_c
h2o_c
nadp_c
coa_c
for_c
h2o_c
ppcoa_c
2mcit_c
h_c
2mcacn_c
h2o_c
h2o_c
micit_c
succ_c
h_c
fum_c
nh4_c
asp__L_c
atp_c
o2_c
adp_c
4pasp_c
h_c
nadph_c
h_c
iasp_c
h_c
pi_c
nadph_c
nadp_c
aspsa_c
atp_c
nadp_c
hom__L_c
h2o_c
adp_c
h_c
phom_c
thr__L_c
pi_c
2obut_c
nh4_c
gly_c
h2o_c
acald_c
nad_c
thf_c
ser__L_c
nh4_c
co2_c
nadh_c
mlthf_c
nh4_c
nadh_c
co2_c
pyr_c
o2_c
mal__L_c
nad_c
oaa_c
glu__L_c
h2o2_c
akg_c
h2o_c
o2_c

### Pathway 44

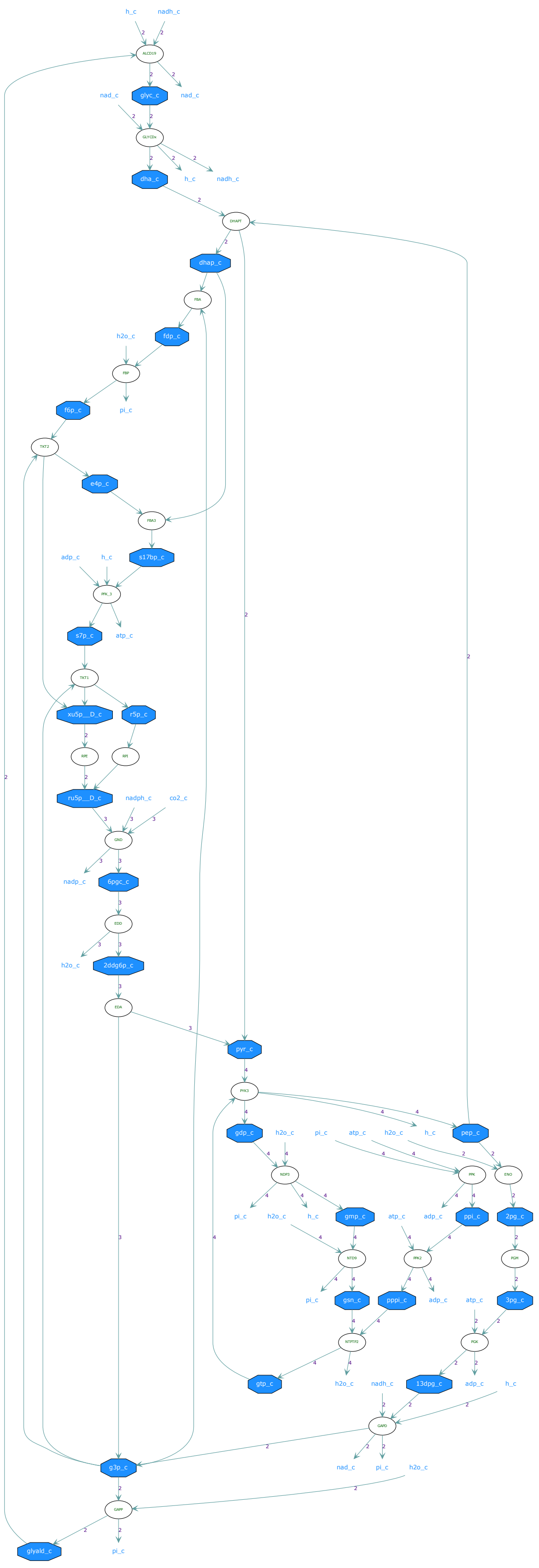

Pathway 45

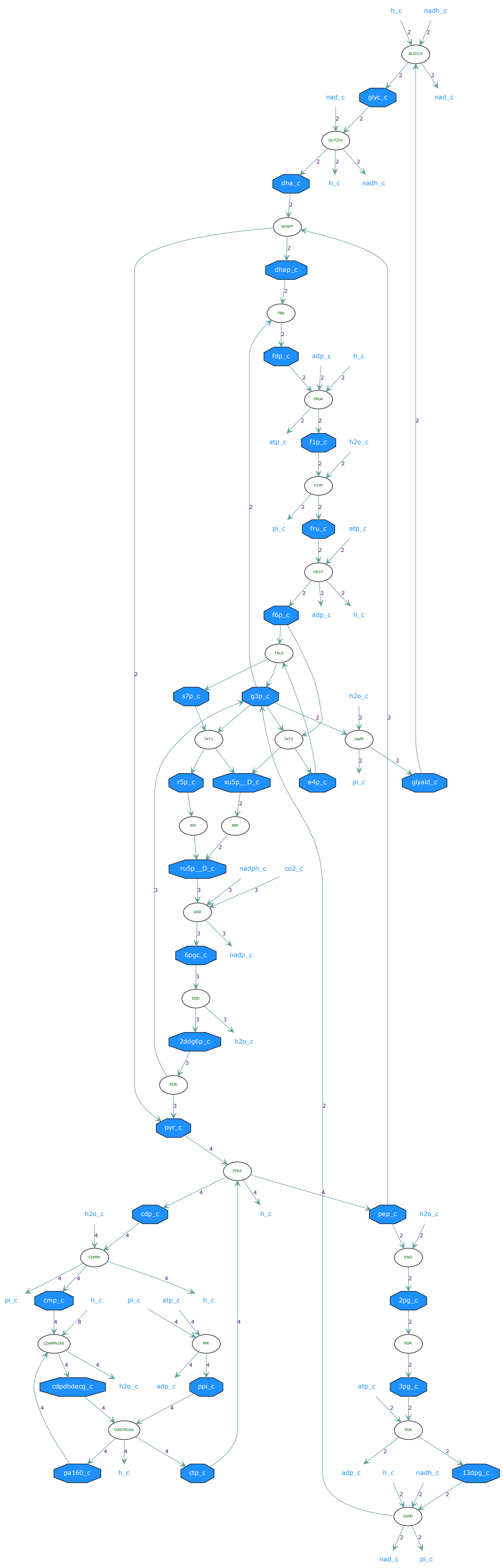

### Pathway 46

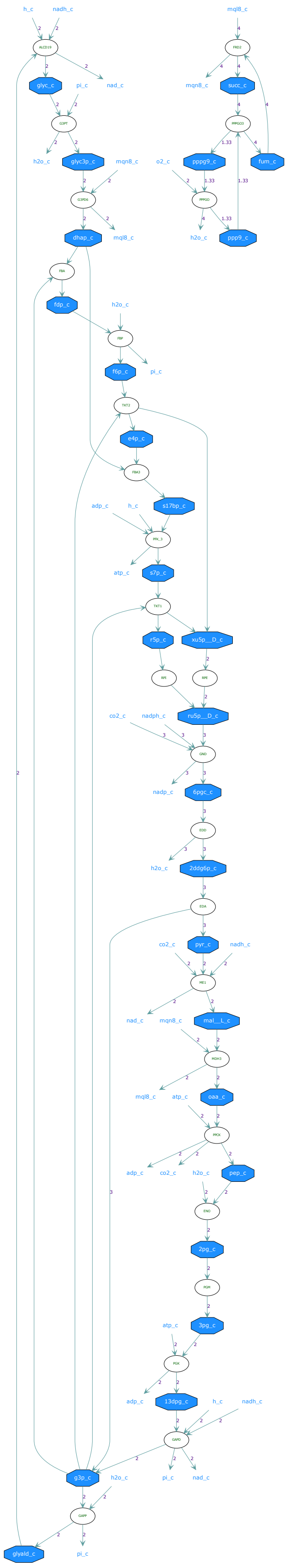

### Pathway 47

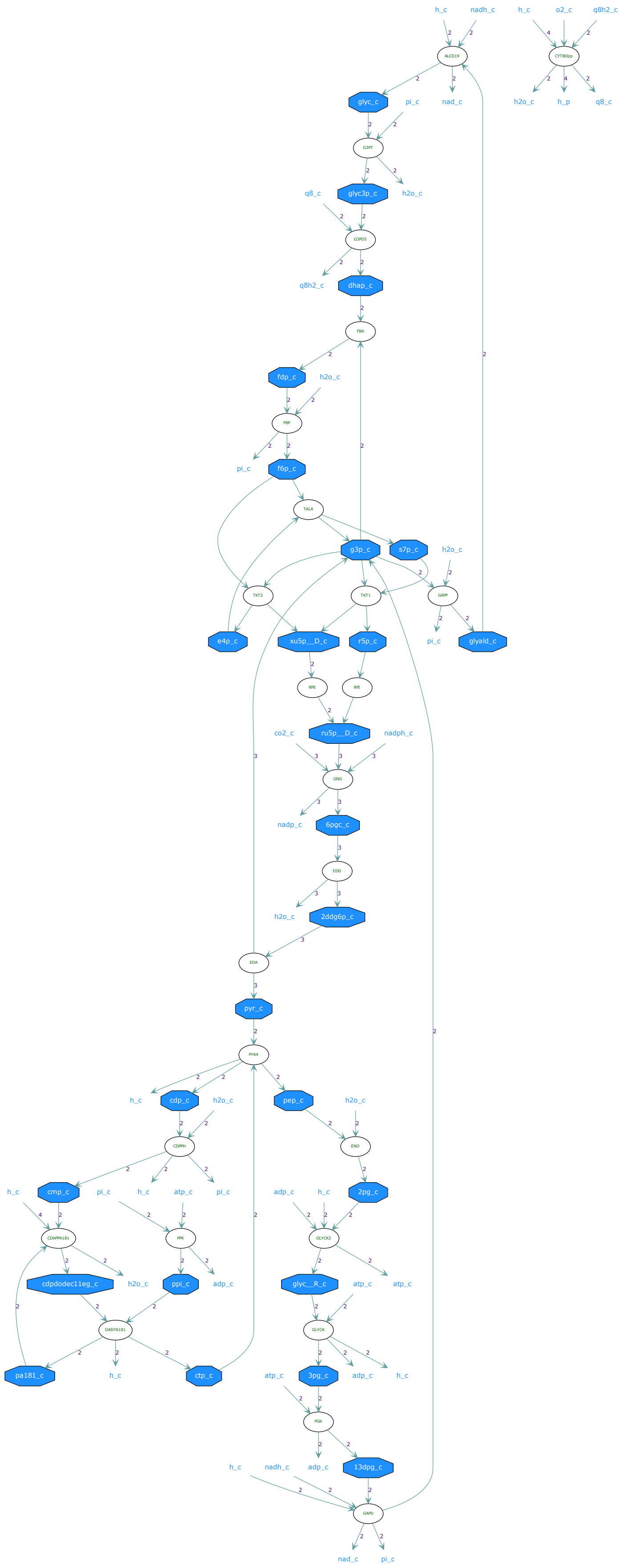
h_c
nadh_c
h_c
o2_c
q8h2_c
ALCD19
CYTBDpp
glyc_c
pi_c
nad_c
h2o_c
h_p
q8_c
G3PT
q8_c
glyc3p_c
h2o_c
G3PD5
q8h2_c
dhap_c
FBA
fdp_c
h2o_c
FBP
pi_c
f6p_c
TALA
g3p_c
s7p_c
h2o_c
TKT2
TKT1
GAPP
e4p_c
xu5p__D_c
r5p_c
pi_c
glyald_c
RPE
RPE
co2_c
ru5p__D_c
nadph_c
GND
nadp_c
6pgc_c
EDD
h2o_c
2ddg6p_c
EDA
pyr_c
PYK4
h_c
cdp_c
h2o_c
pep_c
h2o_c
CDPPH
ENO
h_c
cmp_c
pi_c
h_c
atp_c
pi_c
adp_c
h_c
2pg_c
CDAPPA181
PPK
GLYCK2
cdpdodec11eg_c
h2o_c
ppi_c
adp_c
glyc__R_c
atp_c
atp_c
DASYN181
GLYCK
pa181_c
h_c
ctp_c
atp_c
3pg_c
adp_c
h_c
PGK
h_c
nadh_c
adp_c
13dpg_c
GAPD
nad_c
pi_c

Pathway 48

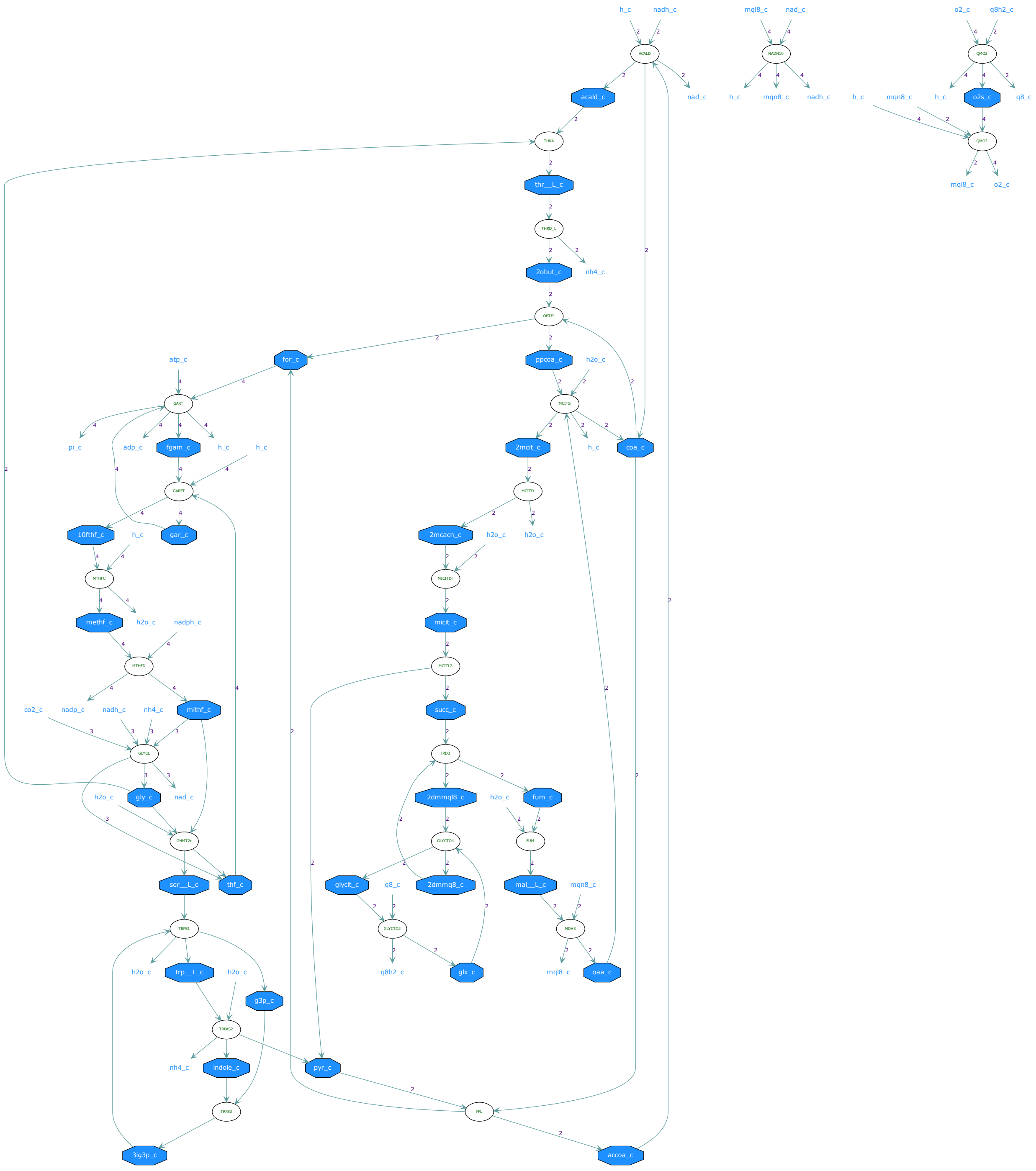
h_c
nadh_c
ACALD
acald_c
nad_c
mql8_c
nad_c
NADH10
h_c
mqn8_c
nadh_c
o2_c
q8h2_c
QMO2
o2s_c
q8_c
QMO3
mql8_c
THRA
thr__L_c
THRD_L
2obut_c
nh4_c
OBTFL
for_c
ppcoa_c
h2o_c
MCITS
2mcit_c
coa_c
MCITD
2mcacn_c
MICITDr
micit_c
MCITL2
succ_c
FRD3
2dmmql8_c
fum_c
FUM
GLYCTO4
2dmmq8_c
glyclt_c
q8_c
GLYCTO2
q8h2_c
glx_c
mal__L_c
mqn8_c
MDH3
oaa_c
atp_c
GART
pi_c
adp_c
fgam_c
GARFT
10fthf_c
gar_c
MTHFC
methf_c
nadph_c
MTHFD
co2_c
nadp_c
nh4_c
mlthf_c
GLYCL
gly_c
GHMT2r
ser__L_c
thf_c
TRPS1
trp__L_c
g3p_c
TRPAS2
indole_c
pyr_c
TRPS3
3ig3p_c
PFL
accoa_c

### Pathway 49

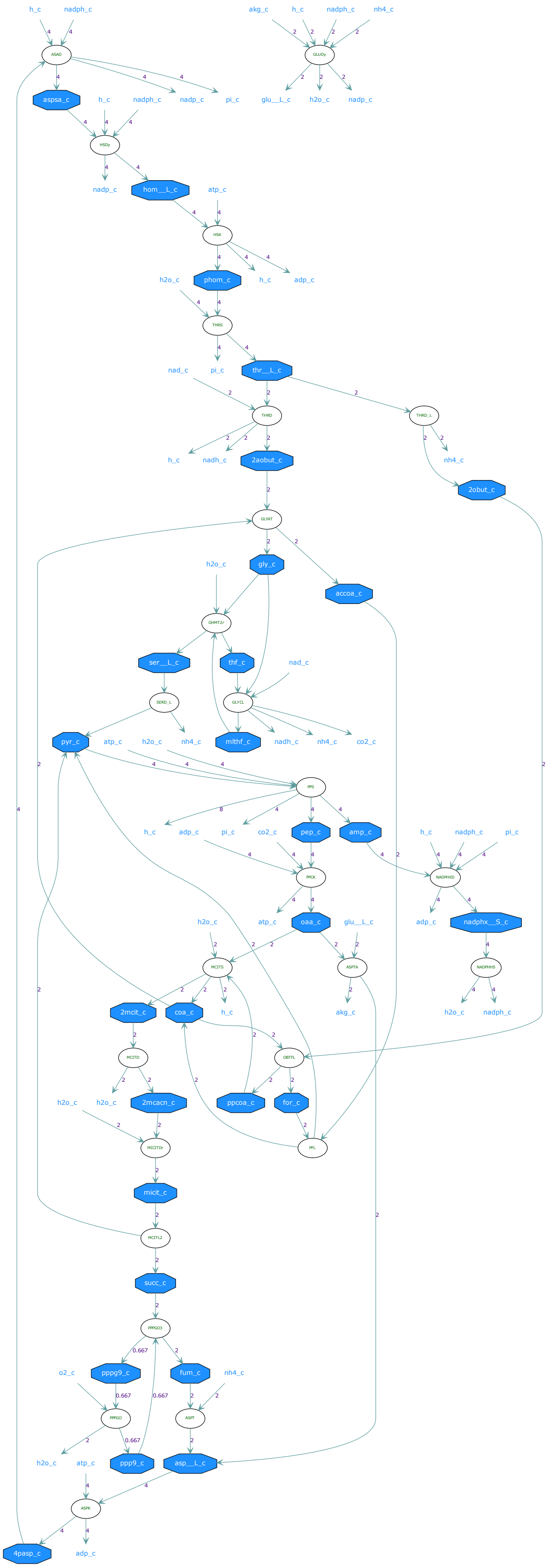
h_c
nadph_c
akg_c
h_c
nadph_c
nh4_c
ASAD
GLUDy
aspsa_c
h_c
nadph_c
nadp_c
pi_c
glu__L_c
h2o_c
nadp_c
HSDy
nadp_c
hom__L_c
atp_c
HSK
h2o_c
phom_c
h_c
adp_c
THRS
nad_c
pi_c
thr__L_c
THRD
THRD_L
h_c
nadh_c
2aobut_c
nh4_c
2obut_c
GLYAT
h2o_c
gly_c
accoa_c
GHMT2r
ser__L_c
thf_c
nad_c
SERD_L
GLYCL
pyr_c
atp_c
h2o_c
nh4_c
mlthf_c
nadh_c
nh4_c
co2_c
PPS
h_c
adp_c
pi_c
co2_c
pep_c
amp_c
h_c
nadph_c
pi_c
PPCK
NADPHXD
h2o_c
atp_c
oaa_c
glu__L_c
adp_c
nadphx__S_c
MCITS
ASPTA
NADPHHS
2mcit_c
coa_c
h_c
akg_c
h2o_c
nadph_c
MCITD
OBTFL
h2o_c
h2o_c
2mcacn_c
ppcoa_c
for_c
MICITDr
PFL
micit_c
MCITL2
succ_c
PPPGO3
o2_c
pppg9_c
fum_c
nh4_c
PPPGO
ASPT
h2o_c
atp_c
ppp9_c
asp__L_c
ASPK
4pasp_c
adp_c
